# On the origin and evolution of the mosquito male-determining factor *Nix*

**DOI:** 10.1101/2022.12.07.519398

**Authors:** James K. Biedler, Azadeh Aryan, Yumin Qi, Aihua Wang, Ellen O. Martinson, Daniel A. Hartman, Fan Yang, Atashi Sharma, Katherine S. Morton, Mark Potters, Chujia Chen, Stephen L. Dobson, Gregory D. Ebel, Rebekah C. Kading, Sally Paulson, Rui-De Xue, Michael R. Strand, Zhijian Tu

**Affiliations:** Department of Biochemistry, Virginia Tech, Blacksburg, VA 24061, USA; Fralin Life Sciences Institute, Virginia Tech, Blacksburg, VA 24061, USA; Department of Entomology, University of Georgia, Athens, GA 30602; Center for Vector-borne Infectious Diseases, Department of Microbiology Immunology and Pathology, Colorado State University, Fort Collins, CO 80523, USA; Department of Entomology, Virginia Tech, Blacksburg, VA 24061, USA; Genetics Bioinformatics and Computational Biology PhD program, Virginia Tech, Blacksburg, VA 24061, USA; Department of Entomology, University of Kentucky, Lexington, KY 40503, USA; MosquitoMate, Inc., Lexington, KY 40502, USA; Anastasia Mosquito Control District, St. Augustine, FL 32092

**Author notes:** Equal contribution (James K. Biedler, Azadeh Aryan, Yumin Qi). Please send correspondence to: Jim Biedler or Zhijian Jake Tu. Department of Biology, University of New Mexico (Ellen O. Martinson). Department of Entomology, Cornell University (Daniel A. Hartman).

## Abstract

The mosquito family Culicidae is divided into two subfamilies named the Culicinae and Anophelinae. *Nix*, the dominant male-determining factor, has only been found in the culicines *Aedes aegypti* and *Ae. albopictus*, two important arboviral vectors that belong to the subgenus Stegomyia. Here we performed sex-specific whole-genome sequencing and RNAseq of divergent mosquito species and explored additional male-inclusive datasets to investigate the distribution of *Nix*. Except for the Culex genus, *Nix* homologs were found in all species surveyed from the Culicinae subfamily, including 12 additional species from three highly divergent tribes comprising 4 genera, suggesting *Nix* originated at least 133-165 MYA. Heterologous expression of one of three divergent *Nix* ORFs in *Ae. aegypti* resulted in partial masculinization of genetic females as evidenced by morphology and *doublesex* splicing. It is not clear whether insufficient transgene expression or sequence divergence or both are responsible for the lack of phenotype for the other two. Phylogenetic analysis suggests *Nix* is related to *femaleless* (*fle*), a recently described intermediate sex-determining factor found exclusively in anopheline mosquitoes. *Nix* from all species has a conserved structure, including three RNA-recognition motifs (RRMs), as does *fle*. However, *Nix* has evolved at a much faster rate than *fle*. The RRM3 of both *Nix* and *fle* are related to the single RRM of a widely distributed and conserved splicing factor *transformer-2* (*tra2*). RRM3-based phylogenetic analysis suggests this domain in *Nix* and *fle* may have evolved from *tra2* in a common ancestor of mosquitoes. Our results provide insights into the evolution of sex-determination and homomorphic sex chromosomes in mosquitoes, and will inform broad applications of mosquito-control strategies based on manipulating sex ratios towards the non-biting males.

## Introduction

The primary signals that initiate sex determination in insects are highly diverse (reviewed in (Bachtrog, et al. 2014)). In the vinegar fruit fly *Drosophila melanogaster,* the double dosage of X differentiates XX from XY embryos and triggers female development (Salz and Erickson 2010). However, it is the presence or absence of a dominant male-determining factor (M factor) that distinguishes the two sexes in many other insects including mosquitoes. All mosquitoes belong to the monophyletic family Culicidae, which is subdivided into two subfamilies: the Anophelinae and Culicinae. The Anophelinae consists of a single genus (*Anopheles*) with well-differentiated X and Y sex chromosomes. Males are also the heterogametic sex that contains the repeat-rich and gene-poor Y chromosome. The Culicinae consists of many, highly divergent genera that form several tribes including the: 1) Aedini that contains the genera *Aedes* and *Psorophora*; 2) Culicini that contains the genus *Culex*; 3) Sabethini that includes the genus *Wyeomyia*; and 4) Toxorhynchitini that contains the genus *Toxorhynchites* (Reidenbach, et al. 2009; da Silva, et al. 2020; Zadra, et al. 2021). The sex-determining chromosomes in culicines are homomorphic or autosome-like except for a pair of sex loci: the male-determining locus or the M locus, and its counterpart the m locus. Similar to *Anopheles*, the Mm male is the heterogametic sex. The M locus is approximately 1.3 Mb in length and resides on chromosome 1 (∼310 Mb) in the yellow fever mosquito *Aedes aegypti* (Matthews, et al. 2018). Classic evolutionary theory suggests homomorphic sex chromosomes will eventually evolve into well differentiated sex chromosomes through a process that involves the accumulation of sexually antagonistic genes, suppressed recombination, and decay of the heterogametic sex chromosome (Charlesworth, et al. 2005; Toups and Hahn 2010; Bachtrog 2013). However, this hypothesis is the subject of recent debate (Lenormand and Roze 2022). It is not clear whether the M- and m-bearing chromosomes in the Culicinae mosquitoes represent nascent proto-Y and proto-X, respectively, or ancient sex-determining chromosomes that maintained their homomorphic form (Toups and Hahn 2010).

Early genetic evidence suggests that the M factor resides in an anopheline Y chromosome and a culicine M locus (Newton, et al. 1974; Baker and Sakai 1976, 1979). *Nix*, an M-linked gene that encodes a predicted RNA-binding protein distantly related to a conserved splicing factor transformer2 (*tra2*), fulfills the role of the M factor in *Ae. aegypti* as it is required for male development and sufficient to convert females into fertile males (Hall, et al. 2015; Aryan, et al. 2020). A *Nix* homolog also functions as the M factor in the Asian tiger mosquito *Aedes albopictus* (Liu, et al. 2020; Lutrat, et al. 2022; Zhao, et al. 2022). In the African malaria mosquito *Anopheles gambiae*, a Y-linked gene, *gYG2/Yob*, encodes the M factor which is a 56 amino acid protein that is apparently not related to *Nix* (Hall, et al. 2016; Krzywinska, et al. 2016). These highly plastic primary signals eventually result in sex-specific splicing of the pre-mRNAs of two conserved transcription factors, *doublesex* (*dsx*) and *fruitless (fru),* producing sex-specific Dsx and Fru protein isoforms that program sexual differentiation. Thus, the sex-determination pathway is thought to evolve in an inverse pattern, with rapid changes of the primary signals that control the conserved downstream factors (Bopp, et al. 2014). Recently, another *tra2*-related gene named *femaleless (fle)* was found exclusively in *Anopheles* mosquitoes (Krzywinska, et al. 2021). In contrast to *Nix* which shifts *dsx* and *fru* splicing towards the male isoforms in *Ae. aegypti* (Hall, et al. 2015; Aryan, et al. 2020), *fle* is required for female-specific splicing of *dsx* and *fru* in *An. gambiae* (Krzywinska, et al. 2021).

Although *Ae. aegypti* and *Ae. albopictus* belong to the same subgenus (Stegomyia), their Nix proteins are only 52% identical over 96% of the protein length. Low conservation and the presence of up to 100 kb-sized introns (Matthews, et al. 2018; Liu, et al. 2020) present significant challenges to identify *Nix* homologs beyond closely related species. In this study, we performed a broad genomics- and transcriptomics-based survey of *Nix* distribution and investigated *Nix* evolution and function. Our results provide insights into the evolution of sex-determination and the enigmatic homomorphic sex chromosomes in mosquitoes, and will inform broad applications of mosquito-control strategies based on manipulating sex ratios towards the non-biting males.

## Results

### Identification and characterization of the *Nix* gene in divergent mosquito species

To characterize *Nix* in divergent mosquito species beyond *Ae. aegypti* and *Ae. albopictus*, we performed high coverage whole-genome sequencing (WGS) of 9 other species in the Culicinae (Table 1; Supplemental Table S1 for details) and developed a bioinformatic method to rapidly identify and characterize the *Nix* gene from unassembled WGS or RNAseq datasets (Figure 1). This method was also used to identify *Nix* from three additional culicine species in male RNAseq databases previously deposited in NCBI (Table 1). All newly characterized *Nix* sequences are provided in Supplemental data 1. Two isoforms were found in *Ae. atropalpus*, one of which retains a 75 nt intron. Four paralogous copies of *Nix* were identified in *Ae. triseriatus*. In total, *Nix* was identified and characterized from 12 additional species in 4 divergent genera *Aedes*, *Psorophora, Toxorhynchites*, and *Wyeomia*. In contrast, despite exhaustive searches in high coverage datasets including raw reads, *Nix* was not found in *Culex quinquefasciatus*, a representative of one of the early-diverged lineages of the Culicinae.

**Figure 1.**
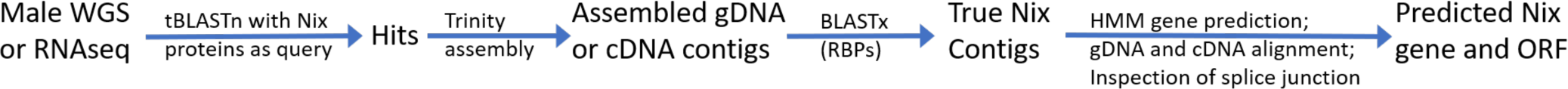
A flow chart showing the process to identify and characterize *Nix*. Details are provided in methods and supplemental methods sections. Note that WGS and RNAseq datasets are not available for all species. To be inclusive of all possible *Nix*-related sequences, tBLASTn was performed under very low stringency (evalue=10) and the query sequences include all known Nix peptides at the time of search. The subsequent BLASTx against a dataset of diverse RNA-binding proteins (RBPs) was used to remove non-*Nix* sequences that better match other related proteins. HMM stands for hidden markov model; specifically, Fgenesh+ (softberry.com) was used for gene prediction using similar protein support.

**Table 1.** Genomic and RNAseq resources generated or used for *Nix* discovery.

| <u>Species</u> | <u>Description</u> | <u>Database Reference</u> |
| --- | --- | --- |
| *Aedes (Stegomyia)* <sup>1</sup><br>*mascarensis* | WGS (male); WGS (female) | This work; PRJNA885905 |
| *Aedes (Stegomyia)*<br>*polynesiensis* | WGS (male); WGS (female) | This work; PRJNA885905 |
| *Aedes (Stegomyia) riversi* | WGS (male); WGS (female) | This work; PRJNA885905 |
| *Aedes (Aedimorphus) vexans* | WGS (male); WGS (female) | This work; PRJNA885905 |
| *Aedes (Hulecoeteomyia)*<br>*japonicus* | WGS (male) | This work; PRJNA885905 |
| *Aedes (Protomacleaya)*<br>*triseriatus* | WGS (male); WGS (female); RNAseq (male) | This work; PRJNA885905 |
| *Aedes (Georgecraigius)*<br>*atropalpus* | WGS (male); WGS (female); RNAseq (embryo) | This work; PRJNA885905 |
| *Aedes (Ochlerotatus)*<br>*detritus* | Published RNAseq (mixed larvae) | Edmunds et al., 2021; PRJNA431084 |
| *Psorophora (Grabhamia)*<br>*columbiae* | WGS (male); WGS (female); RNAseq (male) | This work; PRJNA885905 |
| *Toxorhynchites* sp. <sup>2</sup> | Published RNAseq: Identification to genus only | Kutty et al., 2018; SRR6155935.1 |
| *Toxorhynchites amboinensis* | WGS (male); WGS (female); RNAseq (L3 instar) | This work; PRJNA885905 |
| *Wyeomyia smithii* | Published RNAseq (454; male and female brain Illumina) | Tormey et al., 2015; Basrur et al., 2020<br>SRR11292923-5 (PRJNA612100) |
| *Culex (Culex)*<br>*quinquefasciatus* | *Nix* not found | PRJNA661545 |

Notes: Detailed information on the sample source is provided in Table S1. In addition to morphological identification, species authentication has also been performed by comparing these datasets with sequences of known species origin.
1. Subgenus names are in parentheses.
2. Identification was determined to the genus level.

### *Nix* is male-specific in divergent mosquito species

Male-specificity of *Nix* sequences was first determined by Chromosome Quotient (CQ, Table 2) using previously published methods (Hall, et al. 2013; Matthews, et al. 2018). WGS data were available from both males and females in eight of the culicine species we sequenced (Table 1). *Nix* in all these species is specific to males (CQ<0.01) with the exception of *Ae. triseriatus* which has three male-specific (Ae.triNix1, 2, and 4) and one presumably autosomal paralog (Ae.triNix3, Table 2). No female WGS data were available for *Ae. japonicus,* and no male or female WGS data were available for *Ae. detritus*, one *Toxorhynchites* species, and *W. smithii* (Table 1). Therefore, CQ analysis of *Nix* genes in these species was not possible. However, *Nix* showed no hits in female *W. smithii* RNAseq datasets, consistent with male-specificity. In addition, genomic DNA PCR was performed to confirm the male-specificity of the *Ae. atropalpus Nix* (Supplemental Figure S1). These results are consistent with *Nix* acting as the M factor in divergent species throughout the Culicinae.

**Table 2.** *Nix* is male-specific in divergent mosquito species.

| seq_name | seq_length | female_hits | male_hits | CQ |
| --- | --- | --- | --- | --- |
| Ae.vex.NixORF | 882 | 0 | 175 | 0 |
| Ae.atr.Nix1ORF | 816 | 0 | 121 | 0 |
| Ae.pol.NixORF | 849 | 0 | 270 | 0 |
| Ae.tri.Nix1ORF | 837 | 0 | 157 | 0 |
| Ae.tri.Nix2ORF | 831 | 0 | 101 | 0 |
| Ae.tri.Nix3ORF<sup>1</sup> | 618 | 260 | 276 | 0.94 |
| Ae.triNix4ORF | 321 | 0 | 23 | 0 |
| Tox.amb.NixORF | 858 | 0 | 125 | 0 |
| Ae.riv.NixORF | 849 | 0 | 125 | 0 |
| Ae.mas.NixORF | 867 | 1 | 116 | <0.01 |
| Ps.col.NixORF | 831 | 1 | 139 | <0.01 |

1. This copy is likely autosomal.

### Evolutionary conservation of *Nix* is indicated by three RNA-recognition motifs (RRMs), a conserved intron position, and similar three-dimensional structures

Several characteristics of divergent Nix protein sequences indicate evolutionary conservation. All full-length Nix proteins from the 14 mosquito species we analyzed share three conserved RNA-Binding Domains or RRMs as previously reported for *Nix* from *Ae. aegypti* (Coronado, et al. 2020)(Figure 2). Each RRM contains the characteristic ββαββαβ secondary structure motifs and two short conserved ribonucleotprotein (RNP) motifs (Maris, et al. 2005; Clery, et al. 2008; Muto and Yokoyama 2012), RNP1 and RNP2, having 8 and 6-amino acid consensus sequences, respectively (Coronado, et al. 2020). Only cDNA sequences are available for three species and some *Nix* sequences are from apparently degenerate paralogs. Therefore, these sequences were not included in the intron analysis. Of the 12 remaining sequences from 11 species, all except *Toxorhynchites amboinensis* had an intron in the same position in RRM3, consistent with a common origin. Lastly, AlphaFold structure predictions suggested that divergent Nix proteins share similar three-dimensional structures (Figure 3A-D).

**Figure 2.**
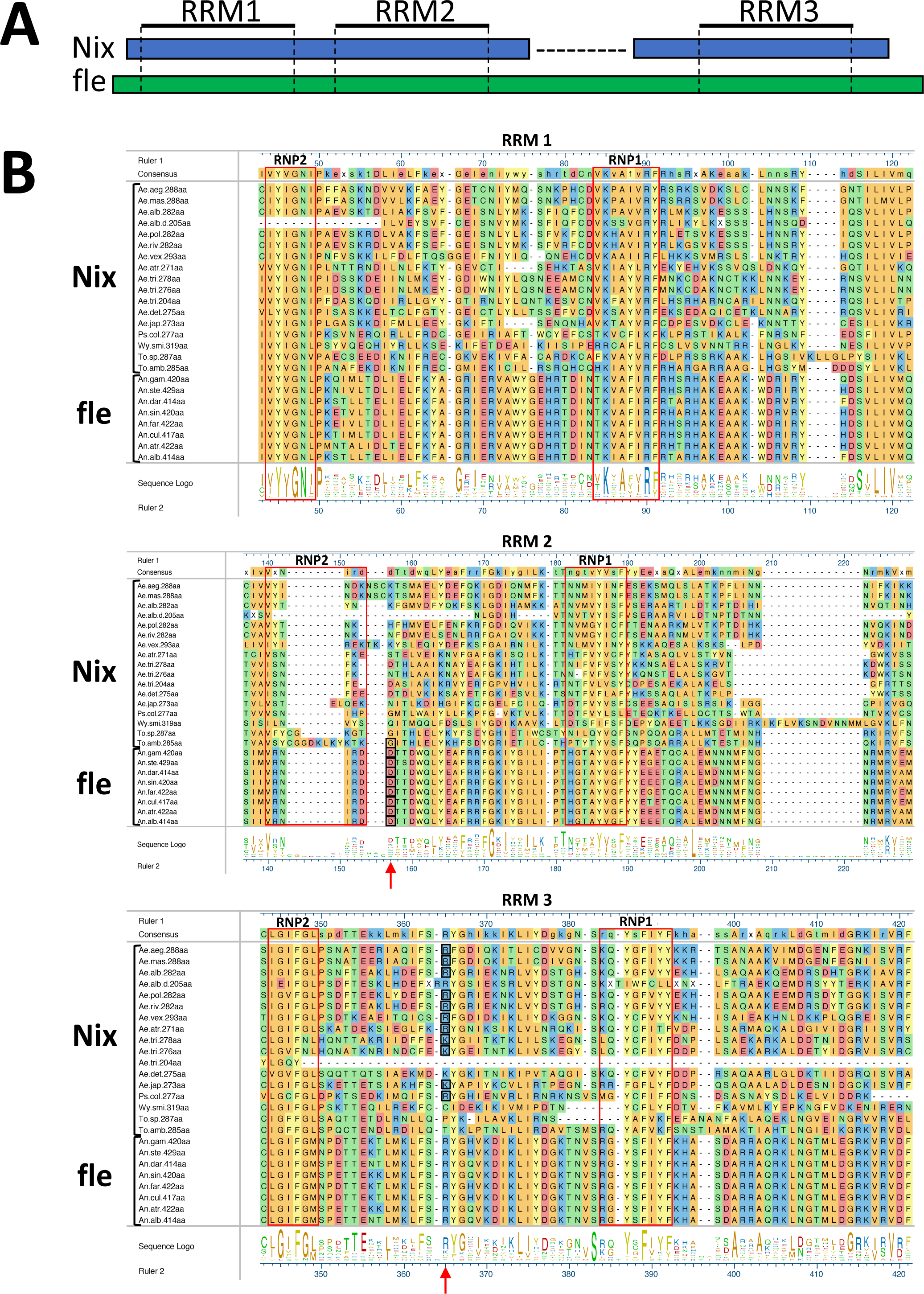
Multiple sequence alignment of *Nix* and *fle* sequences and AlphaFold structure predictions of selected *Nix* and *fle* proteins. A) Schematic shows structure of *Nix* compared to *fle*. B) Multiple sequence alignment of *Nix* and *fle*. Only RRM1, RRM2, and RRM3 are shown. Full alignment is provided as Supplemental Figure S2. Red boxes surround conserved motifs RNP2 and RNP1. Red arrows indicate conserved intron positions with black-boxed residues representing the codon that is split by the intron. Ae.alb.d.205aa is a degenerate copy of *Nix*, and Ae.det.275aa, Wy.smi.319aa, and To.sp.287aa are from RNAseq data only, therefore intron determination was not possible for these sequences. See Table 1 for full species’ names.

**Figure 3.**
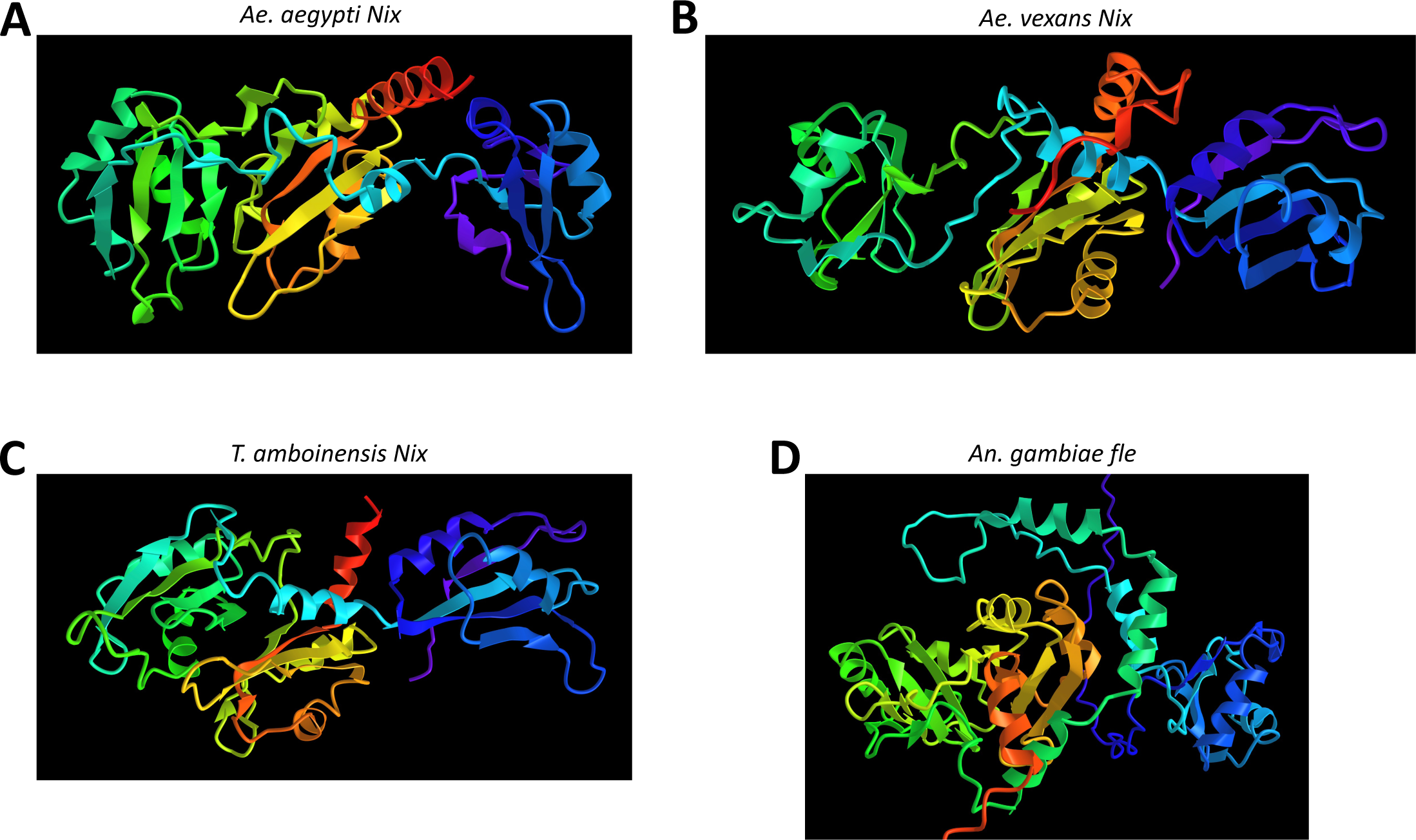
A-D) AlphaFold2 structure predictions (images from NCBI iCn3D viewer) are rainbow-colored (N-terminal red, C-terminal purple) with RRM1 (orange/yellow), RRM2 (green/turquoise) and RRM3 (blue/purple) located in center, left, and right, respectively. Some N-terminal and C-terminal disordered sequence is not show in D.

### *Nix* from the 14 mosquito species form an ancient clade

*Nix* is known to be a distant homolog of a highly conserved splicing factor *tra2* (Hall, et al. 2015). However, the similarity between *Nix* and *tra2* is limited to RRM3 of *Nix* and the single RRM present in *tra2* (Coronado, et al. 2020; Krzywinska, et al. 2021). The closest relative of *Nix*, which also has three RRMs (Figure 2A-B), is a group of *Anopheles*-specific proteins represented by AGAP013051. This gene family has recently been named *femaleless (fle)* as its knockdown confers female-specific lethality due to mis-regulation of X chromosome dosage compensation (Krzywinska, et al. 2021). In contrast to *Nix* which shifts *dsx* and *fru* splicing towards the male isoforms in *Ae. aegypti* (Hall, et al. 2015; Aryan, et al. 2020), *fle* is required for female-specific splicing of *dsx* and *fru* in an *An. gambiae* cell line (Krzywinska, et al. 2021). We thus aligned *fle* from divergent *Anopheles* species with the *Nix* sequences from culicine mosquitoes for comparison (Figure 2B) and to serve as an outgroup to investigate *Nix* evolution.

The entire *Nix* and *fle* alignment (Figure 2B, Supplemental Figure S2) was used to determine the relationship among all Nix protein sequences with *fle* serving as the outgroup for rooting the tree. We performed phylogenetic analysis using RAxML, MrBayes, and BIONJ (Figure 4A-C) (Huelsenbeck and Ronquist 2001; Dereeper, et al. 2008; Stamatakis 2014). *Nix* and *fle* formed two clades with high support, and sequence phylogenies were overall consistent with species phylogeny (Reidenbach, et al. 2009; da Silva, et al. 2020; Zadra, et al. 2021). However, the precise phylogenetic groupings of *Nix* from *Psorophora, Toxorhynchites*, and *Wyeomyia* spp. were not well supported by bootstrap or other statistical replications, perhaps due to sparse sampling and extended evolutionary divergence time among these taxa. In the Baysian analysis (Figure 4B), *Ae. vexans Nix* was the sister taxon of the *Nix* sequences from the Stegomyia subgenus which includes the *Ae. aegypti* and *Ae. albopictus* clades, consistent with the species phylogeny. However, according to the maximum likelihood (Figure 4A) and distance analysis (Figure 4C), *Ae. vexans Nix* was the sister taxon of the *Nix* sequences from the *Ae. aegypti* and *Ae. mascarenesis* clade, albeit with low statistical support. The *Nix* paralogs in *Ae. albopictus* and *Ae. triseriatus* are largely grouped within species, indicating duplication events occurred recently. The branch lengths of sequences in the *Nix* clade are much longer than sequences in the *fle* clade, despite evidence indicating the Anophelinae is basal to the Culicinae (Reidenbach, et al. 2009; Zadra, et al. 2021). Thus, *Nix* evolves at a much faster rate compared to *fle*, which is consistent with the emerging trend of rapid evolution of the primary signals involved in sex-determination (Bopp, et al. 2014) where *Nix* is the primary sex-determining signal in culicine mosquitoes while *fle* is further downstream in the sex-determination pathway of anopheline mosquitoes. Given the distribution of *Nix* and published mosquito species divergence times (Reidenbach, et al. 2009; Zadra, et al. 2021), we estimate *Nix* originated ∼133-165 MYA (Figure 5).

**Figure 4.**
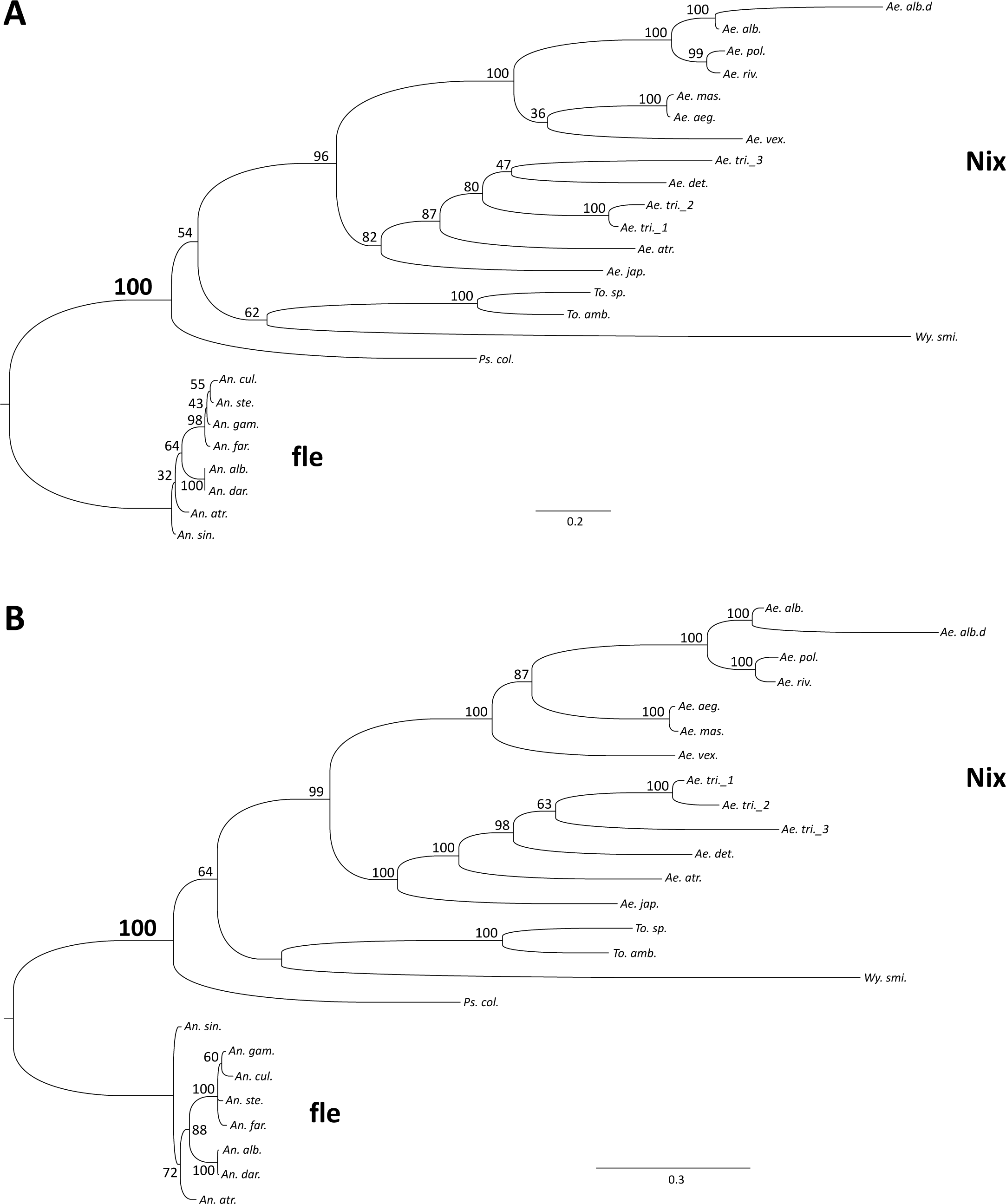
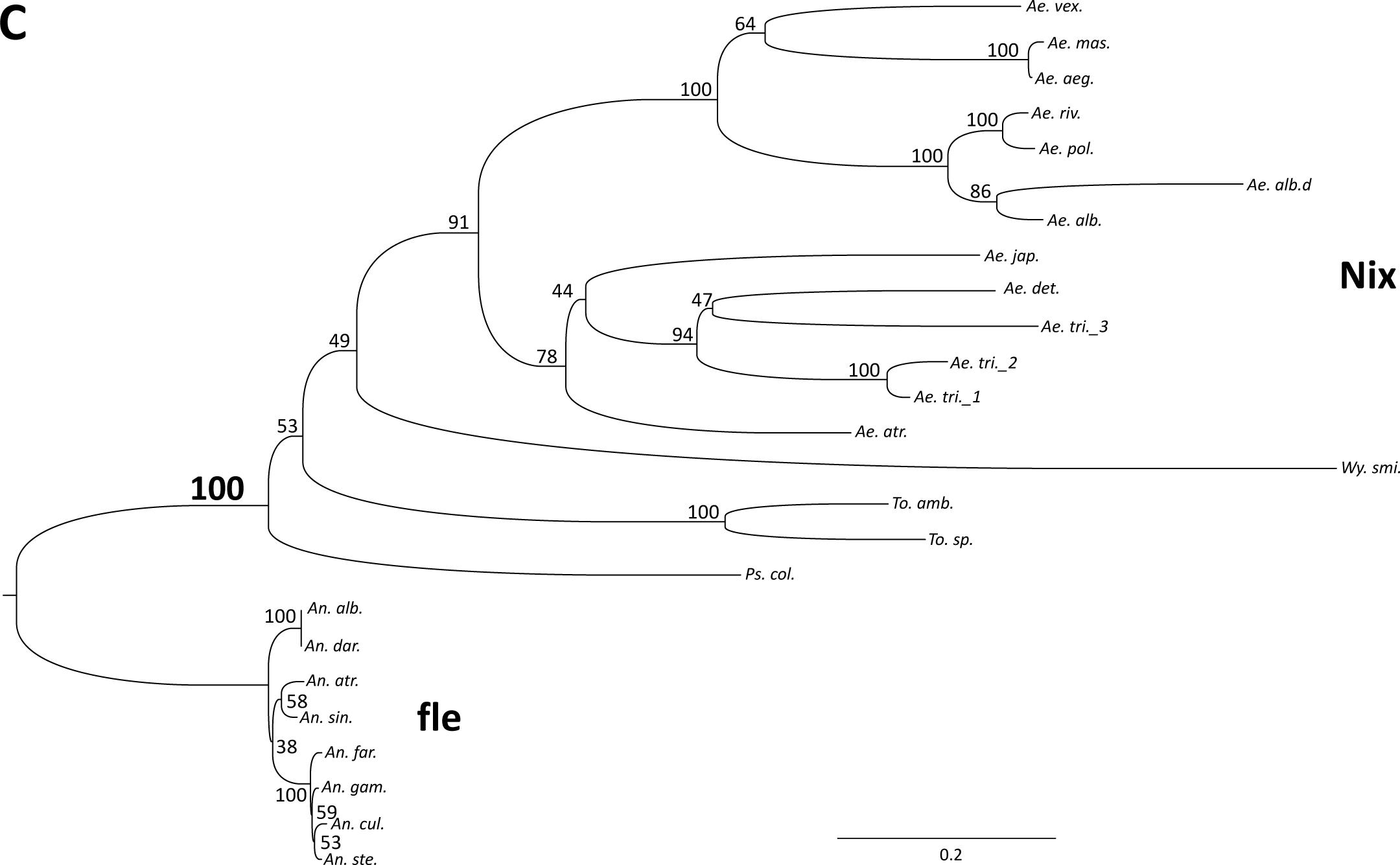
Culicinae *Nix* forms a monophyletic but highly divergent clade when rooted using *fle* as the outgroup. Phylogenetic relation of *Nix* and femaleless (*fle*) was inferred using A) Maximum Likelihood, B) MrBayes, and C) BOINJ. Clade credibility values are indicated. Scale bar shows substitutions per site. See Supplemental Figure S2 for alignment used to infer phylogeny. *fle*, exclusive to Anopheline mosquitoes, was used to root the tree. *Nix* has not been found in *Culex quinquefasciatus*, a species with extensive genomic data. Ae.alb.d represents a degenerate copy of *Nix*. A fourth but truncated *Ae. triseriatus* copy (see Supplemental Data 1) was not included in this analysis. See Table 1 for full species’ names.

**Figure 5.**
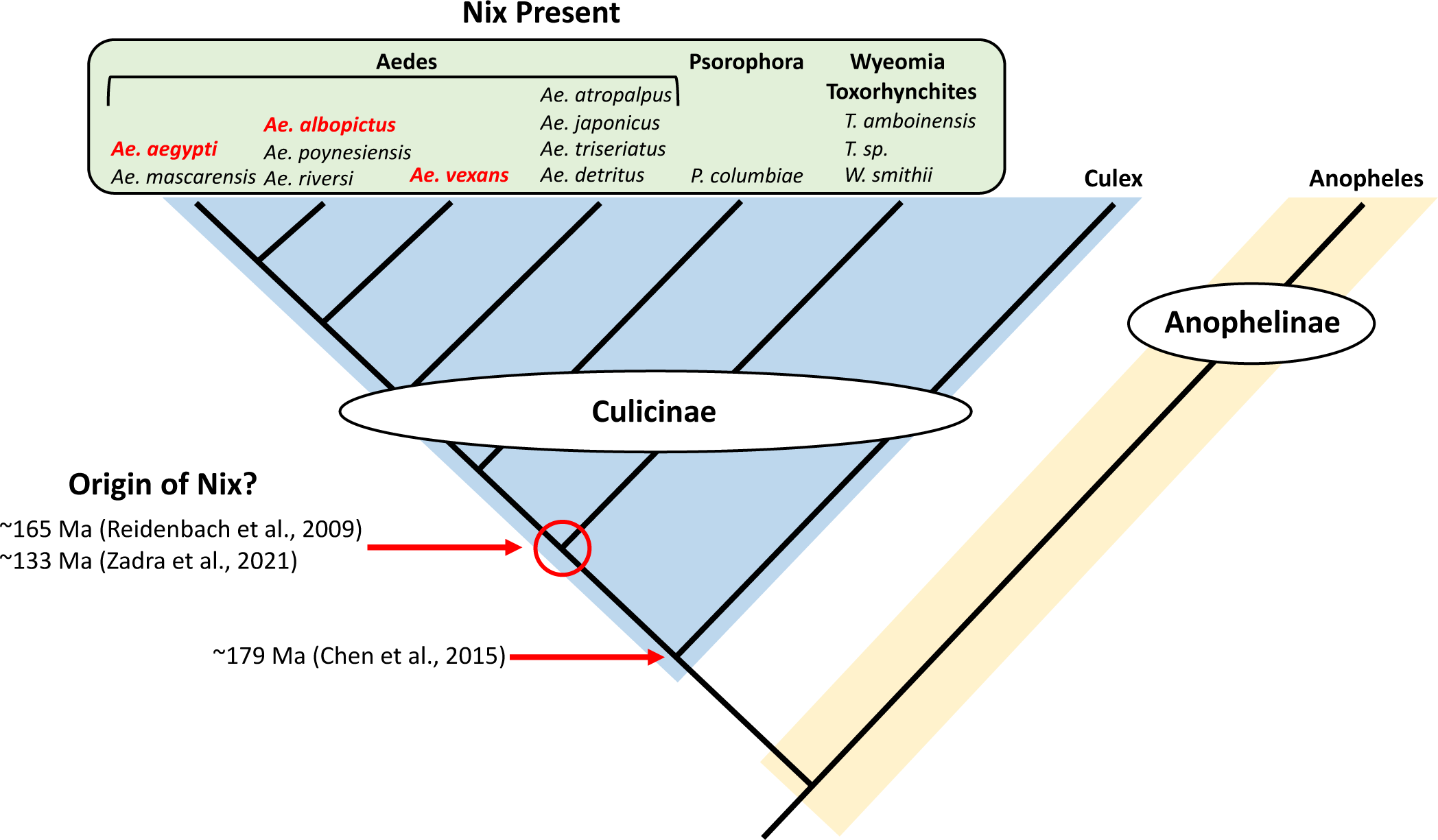
*Nix* is widely distributed and restricted to Culicinae mosquitoes. Cladogram shows relationships between mosquito species based on known mosquito phylogeny (Reidenbach, Cook et al. 2009, da Silva, Machado et al. 2020, Zadra, Rizzoli et al. 2021). Genus names are given at top. The Culicinae and Anophelinae are shaded in light blue and light yellow, respectively. Species for which *Nix* is present are in the green-shaded box and species for which male-determining function of *Nix* has been demonstrated are in red.

### Ae. vexans Nix shows functional conservation in Ae. aegypti while Ae. polynesiensis and Ae. japonicus Nix do not

Our phylogenetic analyses consistently indicated *Nix* from *Ae. vexans* and *Ae. polynesiensis* are more closely related to *Nix* from *Ae. aegypti* than *Nix* from *Ae. japonicus* (Figure 4, 5). We thus tested the conservation of *Nix* function by introducing and expressing each in *Ae. aegypti* via *piggyBac*-mediated transformation (Handler, et al. 1998). The expression cassette (Supplemental Figure S3) utilized the same *Ae. aegypti Nix* promoter/2kb upstream sequence, 5’ and 3’ UTRs that were previously demonstrated to effectively express *Ae. aegypti Nix* and convert *Ae. aegypti* females to flightless males (Aryan, et al. 2020). Transformants were selected using the pUb-EGFP marker in the donor cassette. In lines expressing the *Ae. vexans Nix*, various phenotypes were observed in some genetic females such as partial masculinization, flightless intersex individuals, and females that could never sustain flight (Table 3, Table S2 for detailed phenotypes). We focused on line *Ae.vex.p11* which showed partial masculinization of genetic females (Figure 6). Observed morphological features indicative of masculinization included plumose antennae and presence of gonostyli/gonocoxites, which are male-associated genitalia structures (Figure 6A, Table 3).

**Figure 6.**
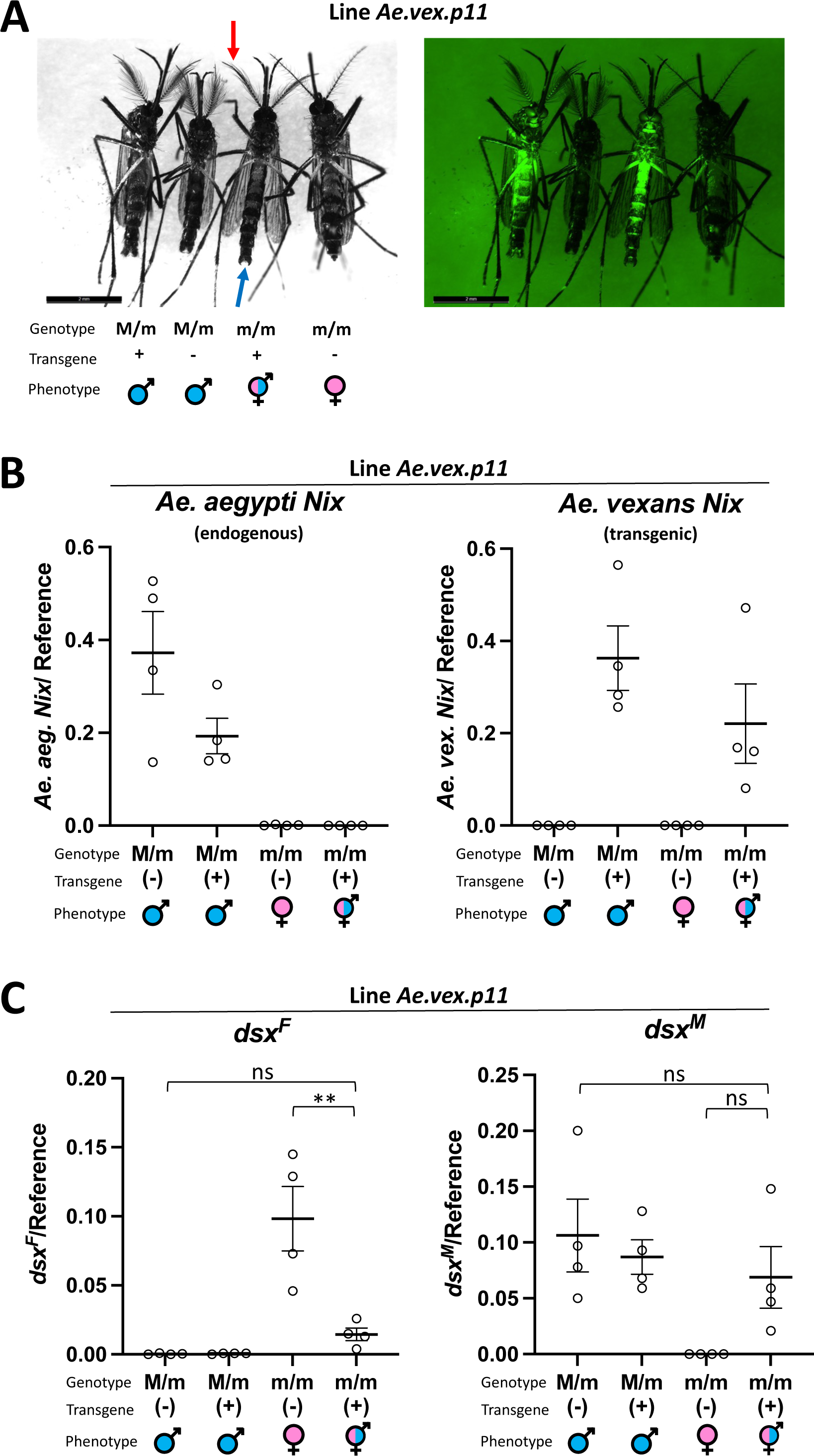
Heterologous expression of transgenic *Ae. vexans Nix* in *Ae.aegypti* masculinizes genetic females by altering the sex determining pathway. A) Images show each of 4 genotype progeny resulting from a cross between transgenic *Ae. aegypti* males having an *Ae. vexans Nix* expression construct, and wild-type females. Image at left was captured with white light. Arrows point to typical male morphological features, plumose antennae (red arrow) and gonocoxites/gonostyli (blue arrow), that are also present in the transgenic intersex individual. Image at right was captured with fluorescent microscope using an EGFP filter, showing transgenic individuals evidenced by EGFP expression in whole body. Left panel and right panel show same individuals in same order. B) Expression of *Ae. aegypti* endogenous *Nix* (left panel) and transgenic *Ae. vexans Nix* (right panel) in *Ae. aegypti* line *Ae.vex.p11* was determined by Droplet Digital PCR relative to gene AAEL002401 used as a control. Genotype, presence of transgene, and phenotype are indicated at bottom. The expression of the transgene in genetic females and males was not significantly different than the endogenous *Nix* expression in WT males (Tukey’s Multiple Comparison Test). C) Expression of female and male *dsx* isoforms (*dsx^F^* and *dsx^M^*) was determined by Droplet Digital PCR relative to an endogenous gene AAEL002401 used as a control. Adult progeny with 4 resulting genotypes from a cross of transgenic males and wild-type females were assayed. X-axis labels: genotypic sex (male, M/m; female; m/m); (+)/(-) indicates presence/absence of the *Ae. vexans Nix* transgenic cassette determined by a fluorescent marker; symbols indicate phenotypic sex (male, female, and intersex). In B) and C) Individual values are shown with the mean and +/- SEM. Statistically significant differences between transgenic genetic females and WT males/WT females are indicated from One-way ANOVA followed by the Tukey’s Multiple Comparison Test.

**Table 3.** Transgenic lines and phenotypes.

| Transgenic Nix Donor Plasmid | #G0 families (# families with G1 positives) | Phenotype of G1 females |
| --- | --- | --- |
| 187_*Ae. japonicus* | 13 (4) | normal |
| 188_*Ae. polynesiensis* | 18 (7) | normal |
| 189_*Ae. vexans* | 12 (7) | flightless, flightless intersex, normal |

Notes: All transgenic males appeared morphologically normal. The phenotypes for line 189\_*Ae. vexans*.Nix were consistent for 10 generations. See Supplementary Table S2 for detailed record of phenotypes.

The lack of full conversion of genetic females to males may be explained by molecular divergence of the *Ae. vexans Nix* from *Ae. aegypti Nix* and/or insufficient expression of the *Ae. vexans Nix* transgene. However, expression of *Ae. vexans Nix* in transgenic genetic females was not significantly different than *Ae. aegypti* WT *Nix* according to Droplet Digital PCR (ddPCR) (Figure 6B). RT-PCR additionally demonstrated expression of the transgenic *Ae. vexans Nix* in only transgenic individuals (Supplemental Figure S4). *Doublesex* is a highly conserved gene at the bottom of the sex determination pathway and is differentially spliced in males vs. females (Shukla and Nagaraju 2010; Verhulst and van de Zande 2015), allowing evaluation of sex-determination signaling in individuals. We performed ddPCR to detect perturbation of sex-determination signaling in transgenic genetic females by assaying for *dsx^F^* and *dsx^M^* isoforms. As expected we detected a shift of *dsx* isforms to male-like patterns (Figure 6C). These results show the function of *Ae. vexans Nix* has been sufficiently conserved to affect sex determination in *Ae. aegypti*.

In contrast, multiple independent transgenic lines expressing each *Nix* ORF from *Ae. polynesiensis* and *Ae. japonicus* produced no detectable phenotypes in *Ae. aegypti* (Table 3). Having multiple independent lines tends to mitigate the potential problem of position effect on transgene expression from a single insertion line. However, when we measured transcript level of the *Ae. polynesiensis* and *Ae. japonicus Nix*, transgenic *Nix* expression was significantly lower than the endogenous *Nix* as determined by ddPCR (Figure 7). It has been suggested that an enhancer may exist in the *Nix* coding region as the *Nix* promoter, when used to drive the expression of *tetracycline transactivator* (tTA), produced significantly less tTA transcripts than the endogenous *Nix* transcripts (Kojin, et al. 2022). It is possible that the *Ae. polynesiensis* and *Ae. japonicus Nix* ORFs lack such an enhancer. Consistent with low expression of the *Ae. polynesiensis* and *Ae. japonicus Nix*, *dsx* splicing was not altered in transgenic females compared to WT (Figure S5). However, we cannot rule out *Nix* sequence divergence being the cause of the lack of phenotype. It is interesting to note *Ae. polynesiensis* is in the subgenus *Stegomyia* with *Ae. aegypti*, whereas *Ae. vexans* is in the subgenus *Aedimorphus* (Table 1, Figure 5).

**Figure 7.**
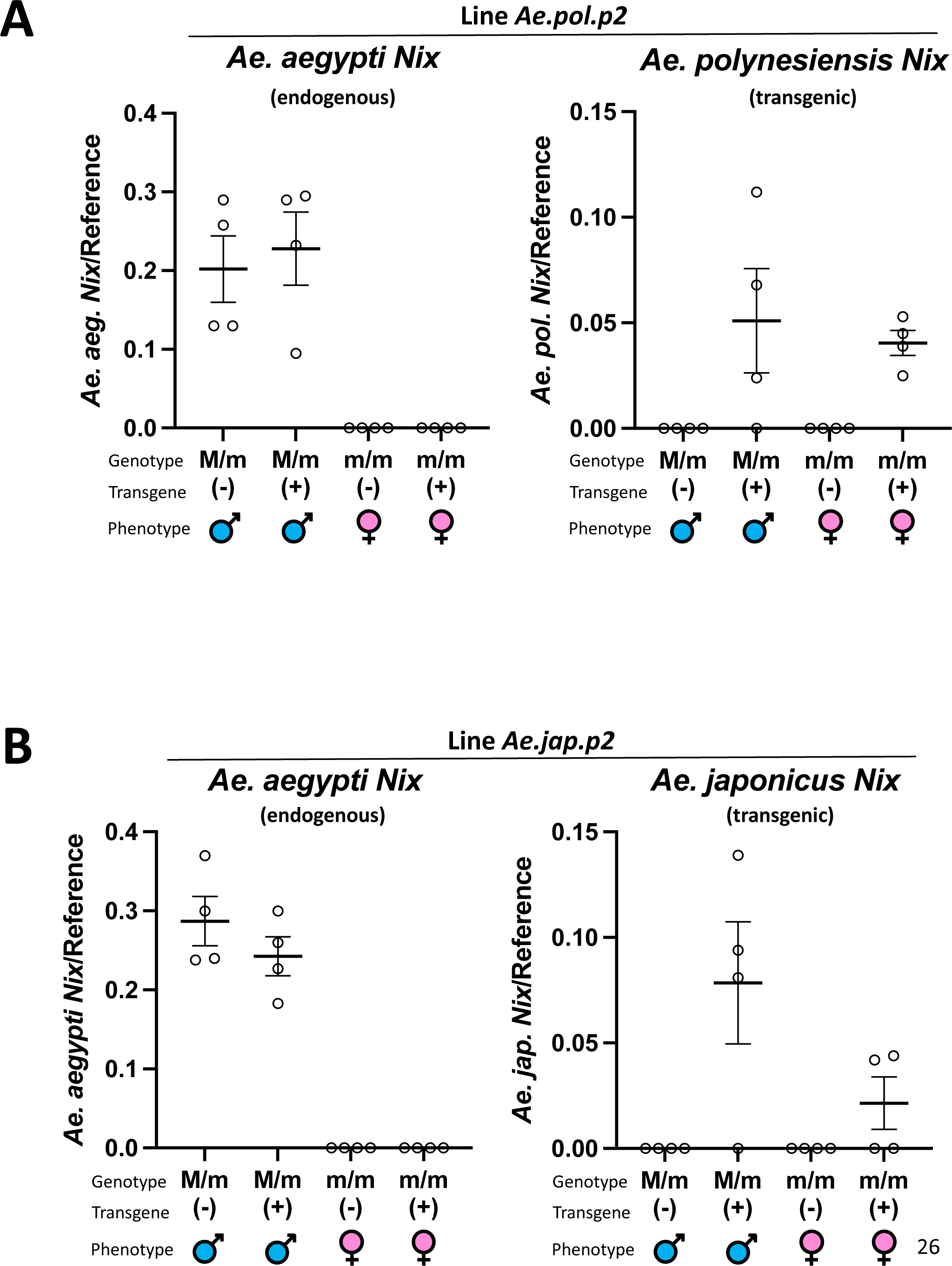
Transcription of transgenic *Ae. polynesiensis* and *Ae. japonicus Nix* compared to native *Nix*, in *Ae. aegypti* transgenic lines. A) Expression of *Ae. aegypti* endogenous *Nix* (WT) and transgenic *Ae. polynesiensis Nix* was determined by ddPCR in the *Ae. aegypti* transgenic line Ae.pol.p2, relative to gene AAEL002401 used as a control. B) Expression of *Ae. aegypti* endogenous *Nix* (WT) and transgenic *Ae. japonicus Nix* was determined by ddPCR in the *Ae. aegypti* transgenic line *Ae.jap.p2*, relative to gene AAEL002401 used as a control. In *Ae. polynesiensis* and *Ae. japonicus Nix*-expressing transgenic lines, the expression of the transgene in genetic females and males was significantly lower than the endogenous *Nix* expression in WT males according to One-way ANOVA followed by the Tukey’s Multiple Comparison Test (*Ae. polynesiensis*, p = 0.0016; *Ae. japonicus*, p < 0.0001). Individual values are shown with the mean and +/- SEM.

### *Nix* and *fle* share a common origin that may have evolved from *tra2* in a common ancestor of mosquitoes

As noted above, the closest relative of *Nix* is *fle* with both sharing three RRMs, while *tra2* is thought to be a distant homolog on the basis of *Nix* RRM3 sharing similarity with the single RRM present in *tra2* (Coronado, et al. 2020; Krzywinska, et al. 2021). *An. ga*mbiae *fle* is 420 aa compared to *Ae. aegypti Nix* being 288 aa but alignment revealed conservation throughout the majority of each, including all 3 RRMs, which strongly suggests homology between the two families. A notable difference between them is the approximately 80 aa sequence that is found in *fle* between RRM2 and RRM3, that is not found in *Nix* sequences (Figure 2A). In search of more support for homology between *Nix* and *fle* gene families, we looked for conservation of intron position in these sequences (Figure 2B, black-boxed residues). All *fle* genes analyzed have a conserved intron at the beginning of the coding sequence for RRM2, ranging in size from 89 to 249 nucleotides. The *Nix* sequence from *T. amboinensis* also has a predicted 152 nt intron interrupting a codon in the same position in the alignment as the intron for the *fle* sequences, supporting a common origin of *T. amboinensis Nix* and *fle* family sequences.

To further explore potential ancestral relationships between *Nix*, *fle*, and *tra2*, we performed phylogenetic inference with RAxML, MrBayes, and BIONJ, using a multiple sequence alignment of the RRM3 domain of *Nix* (14 species) and *fle* (8 anopheline species), and the RRM domain of *tra2* sequences from mosquitoes and several other insect species (Figure 8, alignment provided in Supplemental Figure S6). A *tra2* clade and *Nix*/*fle* clade is evident and well-supported. In contrast, our results provided only weak support for *Nix* and *fle* clades presumably due to high degree of divergence in RRM3. Yet as earlier noted, using full-length sequence with all 3 RRMs yielded very high support for *Nix* and *fle* clades (Figure 4). Thus, our results overall support the hypothesis that *Nix* and *fle* evolved from an ancestral gene in the common ancestor of the Culicidae that subsequently diverged into their unique respective roles in sex-determination in the Culicinae and Anophelinae, which have different types of sex chromosomes. The phylogeny of *tra2* homologs is also overall consistent with known insect species phylogeny, supporting multiple *tra2* duplications (*tra2-*alpha and *tra2-*beta) in the Culicidae (Li, et al. 2019). Assuming *Nix*/*fle* evolved from a *tra2* ancestral sequence, these results suggest yet another duplication of *tra2* in the ancestor of mosquitoes, which subsequently evolved independently in the Culicinae and Anophelinae as *Nix* and *fle*, respectively.

**Figure 8.**
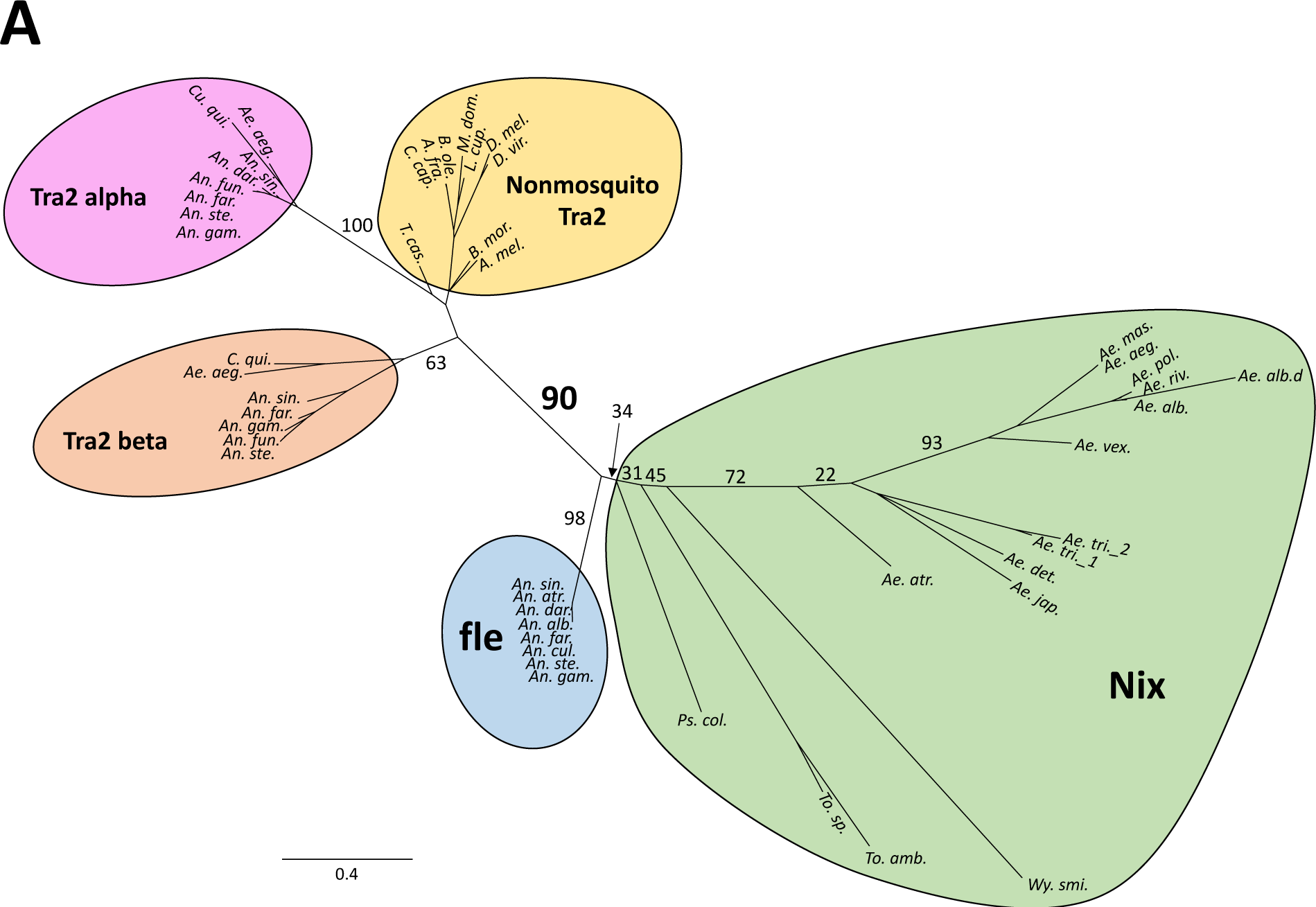
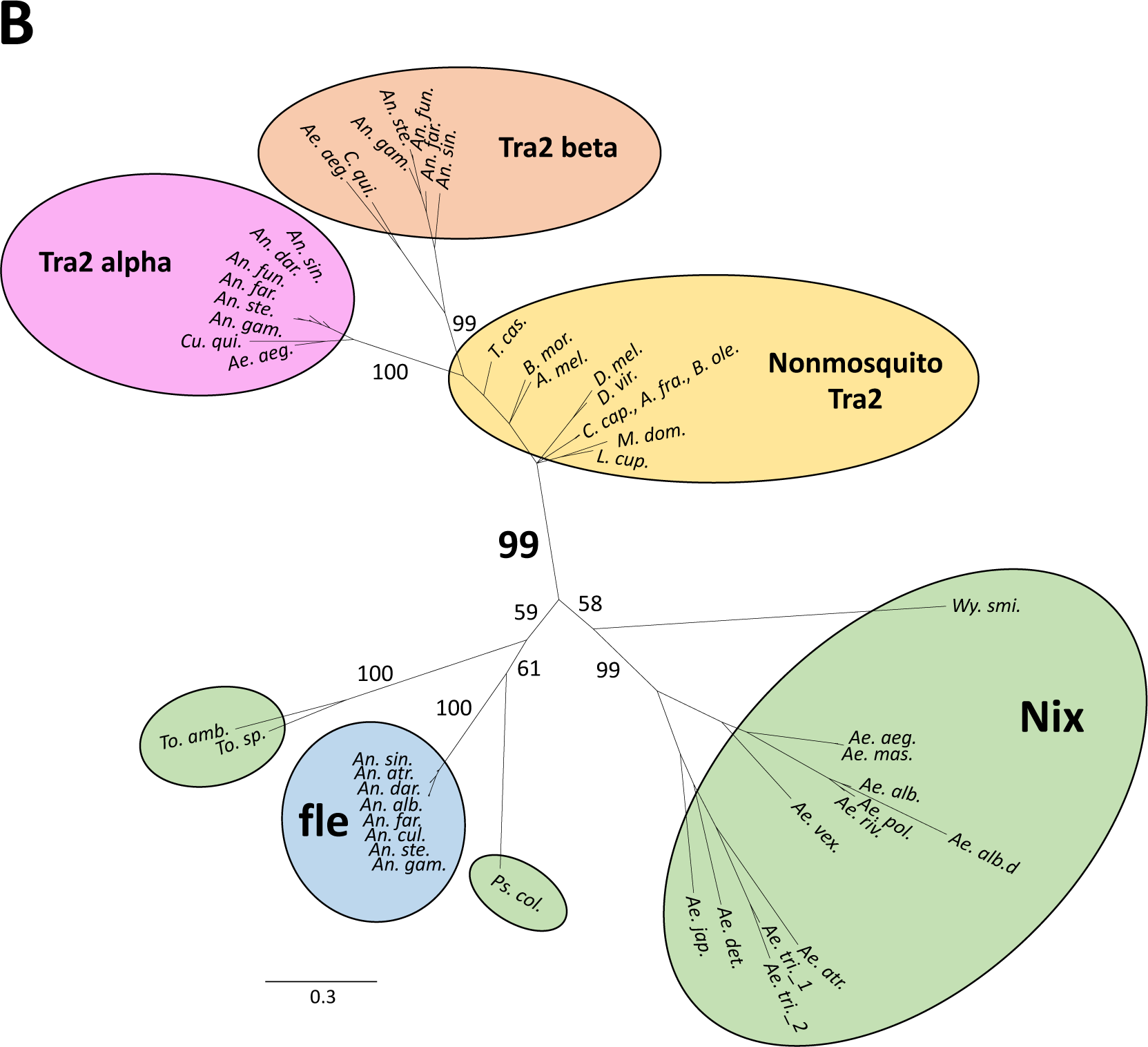
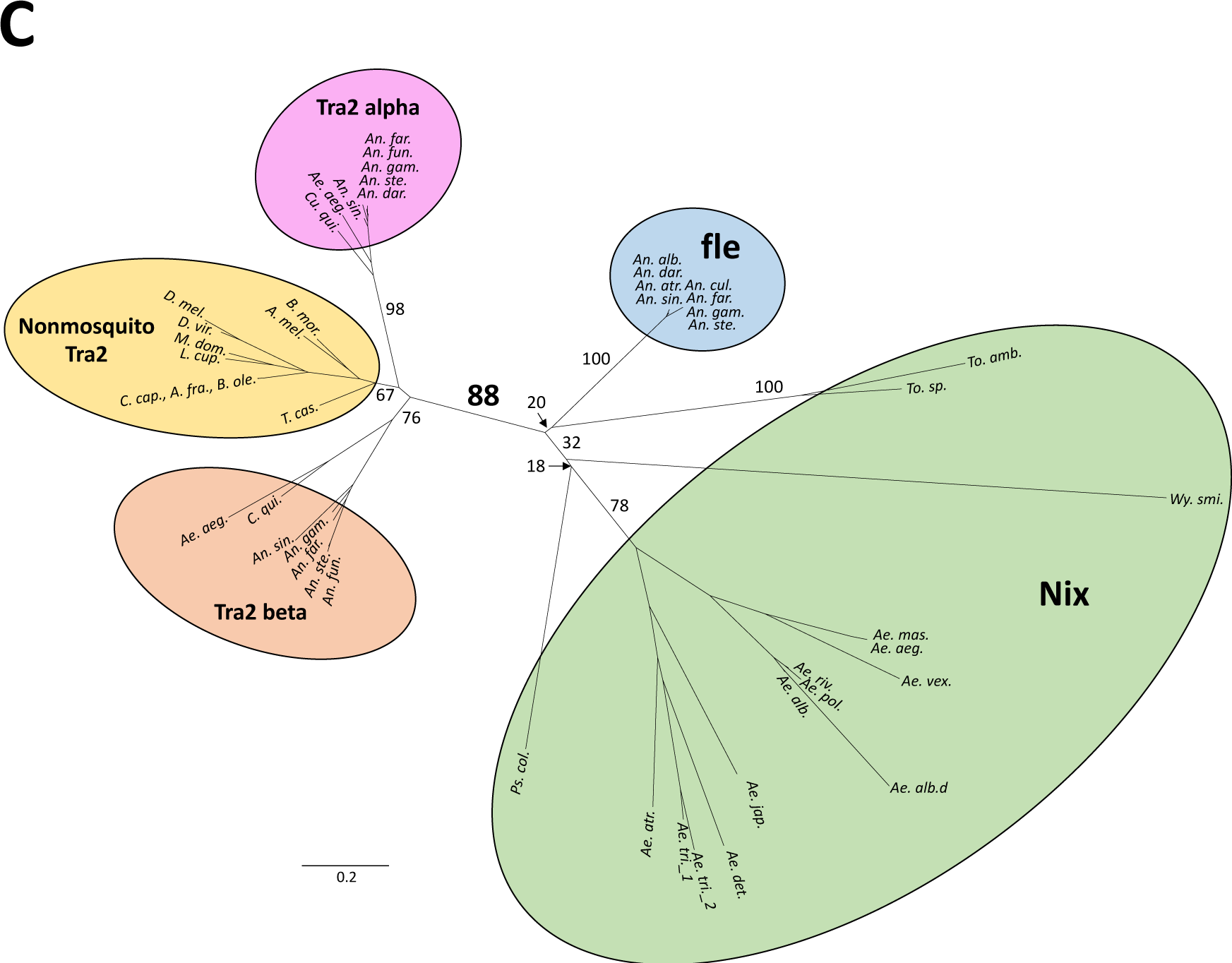
*Nix* is widespread in culicine mosquitoes and forms a sister clade to *fle* which is exclusive to anopheline mosquitoes. Shown are the unrooted phylogenies of *Nix*, femaleless (*fle*), and transformer 2 (Tra2) using amino acid sequences from RNA recognition motif 3 (RRM3) from *Nix* and *fle*, and the single RRM from Tra2. All sequences are from mosquitoes except the sequences in the “Nonmosquito Tra2” clade/group. Clade credibility values are shown for major clades. The labels for clade credibility values for the branch separating Tra2 and *Nix*/*fle* clades are larger and in bold for emphasis. Scale bar shows substitutions per site. See supplemental figure S6 for the RRM 3 alignment used for phylogenetic inference and supplemental figure S7 for phylogenies showing species names and all clade credibility values. *Ae.tri._3* is not included in these trees because it does not have an RRM3. A) Maximum Likelihood Tree generated by RAxML using 1000 bootstrap replicates. B) Tree generated by MrBayes with 1 million generations. *Nix* sequences from three species (*Ps. col, To. amb., To. sp.*) were not grouped in the *Nix* clade but are labeled in a green bubble like all other *Nix* sequences. C) Distance tree generated by BIONJ using 1000 bootstrap replicates. See Table 1 for full species’ names. Nonmosquito species are *Bombyx mori*, *Apis mellifera*, *Drosophila melanogaster*, *Drosophila virilis*, *Musca domestica*, *Lucilia cuprina*, *Ceratitis capitata*, *Bactrocera oleae*, *Anastrepha fraterculus*, *Triboleum castaneum*.

## Discussion

This study identifies *Nix* family members in 14 species among 3 of 4 tribes in the subfamily Culicinae with the only exception being *C. quinquefasciatus*, which is a basal member of the subfamily. Our results suggest *Nix* originated at least ∼133-165 MYA according to published mosquito species divergence times (Reidenbach, et al. 2009; Zadra, et al. 2021) (Figure 5). Phylogenetic inference, sequence comparison, and conserved intron position all indicate that *Nix* and *fle* present in *Anopheles* spp. share a common origin (Figures 2, 4, 8). Neither *Nix* nor *fle* has been found in *C. quinquefasciatus*. A broader survey of other *Culex* spp. and Culicinae tribes will help ascertain the precise evolutionary origin of *Nix*. It is possible *Culex* either lost *Nix* early in its evolution or this gene has diverged beyond recognition relative to the other culicine genera we analyzed. In this context, we note that *Nix* has evolved at a much faster rate than *fle* (Figure 4).

Our transgenic assays indicate that the *Ae. vexans Nix* can function at least partially as the M factor in *Ae. aegypti* (Figure 6), extending functional evidence for *Nix* beyond the *Stegomyia* subgenus of *Aedes*. However, *Nix* ORFs from *Ae. polynesiensis* and *Ae. japonicus* failed to produce detectable phenotypes in *Ae. aegypti* (Table 3, Figure 7, Figure S5). As the lack of phenotype could result from insufficient transgene expression or sequence divergence in coding or other *cis*-required elements, it is important to determine the function of divergent *Nix* genes in their native species, which would likely require germline transformation (e.g.,(Aryan, et al. 2020)), transient embryonic injections (e.g., (Hall, et al. 2015)), or manipulation of sex-specific cell lines (e.g.,(Krzywinska, et al. 2021)) in these species. There is at least one *Nix* copy in all 14 species that is predicted to encode a full Nix protein, although duplicated copies showed signs of degeneration in *Ae. albopictus* and *Ae. triseriatus* (Supplemental Data 1). In addition, all *Nix* copies except a duplication in *Ae. triseriatus* have been shown to be male-specific whenever sex-specific comparisons are possible (Table 2). Thus, all current evidence is consistent with sustained *Nix* function in male mosquitoes as a male-determining factor throughout its ∼133-165 MY of evolution.

We further explored the relationship between *Nix*/*fle* and their distant homolog *tra2* (Hall, et al. 2015; Krzywinska, et al. 2021). In divergent species from flies to humans, *tra2* is a single copy gene that produces various splice isoforms (Dauwalder, et al. 1996; Beil, et al. 1997; Sarno, et al. 2010; Li, et al. 2019). Mosquitoes are the only insect family in which *tra2* duplications have been reported (Li, et al. 2019). All mosquitoes have at least two copies of the *tra2* gene, each forming a monophyletic clade indicating that the duplication occurred in the common ancestor of the mosquito family. One of the *tra2* clades, *tra2-*beta, is evolving at a faster rate than the other *tra2-*alpha clade (Figure 8), indicating possibly relaxed evolutionary constraints or neofunctionalization. *tra2* homology to *Nix* and *fle* is restricted to the RRM3. RRM3-based phylogenetic analysis is consistent with the hypothesis that *Nix* and *fle* evolved from another duplication of *tra2* in the ancestor of mosquitoes. This hypothesis also requires the addition of 2 more RRMs in the ancestor of *Nix*/*fle*.

It is not clear how long the homomorphic sex chromosomes have been maintained in culicine mosquitoes. Although the M locus is only ∼1.3 Mb (Matthews et al., 2018) in *Ae. aegypti*, a >60 Mb M-linked region showed significant differentiation between the M- and m-bearing chromosomes (Fontaine, et al. 2017; Matthews, et al. 2018; Chen, et al. 2022). It is currently unknown whether such a differentiation results from sex-related adaptation or simply reflects that the sex locus resides in a vast recombination desert, which is found near the centromere of all three chromosomes (Fontaine, et al. 2017; Chen, et al. 2022). Investigation into the relationship between *Nix*-linked male-determining regions across divergent species by synteny of gene blocks should provide further insights into the origin, age, and evolution of the homomorphic sex chromosomes in culicine mosquitoes. If the culicine homomorphic sex chromosomes are as old as the *Nix* gene, they will be highly attractive systems for shedding new light on how homomorphic sex chromosomes maintain homomorphy over an extended evolutionary time scale.

Genetic manipulation of the sex-determination pathway is being explored as a new way to control mosquito-borne infectious diseases as the female sex is responsible for pathogen transmission and population expansion. The identification and characterization of *Nix* in diverse mosquito species opens the door to future control applications beyond *Ae. aegypti* and *Ae. albopictus*.

## Materials and Methods

### Samples, library preparation, and Illumina sequencing

Mosquito species used for high coverage whole-genome sequencing (WGS) and RNAseq are shown in Table 1 and detailed information including sex, number of individuals are provided in Supplemental Table S1. Genomic DNA was isolated using Quick-DNA Miniprep Kit (Zymo Research, Irvine, CA) or QIAamp DNA Micro Kit (Qiagen, Hilden, Germany) for pooled or single mosquito samples, respectively, according to the manufacturer’s instructions. Library preparation was performed using NEBNext Ultra II FS DNA Library Prep Kit for Illumina (New England BioLabs Inc., Ipswich, MA, USA). The quantity of the prepared DNA libraries was determined using a Qubit© 2.0 Fluorometer (Thermo Fisher Scientific Inc., Waltham, MA, USA). Quality control was done by the Agilent 2100 Bioanalyzer prior to sequencing. RNA was isolated using either the Quick-RNA Microprep (embryonic samples) or the Quick-RNA Miniprep kit (all other samples) from Zymo Research (Irvine, CA). Libraries were prepared using the NEBNext Ultra RNA Library Prep Kit for Illumina with the Poly(A) mRNA Magnetic Isolation Module. Illumina sequencing was done either by the Virginia Tech Genomics Sequencing Center or Novogene (en.novogene.com) and the specific sequencing platform for each sample varied as indicated in submissions to the Sequence Read Archives (SRA, PRJNA885905).

### Identifying and assembling Nix sequences from WGS and RNAseq datasets

The workflow for the identification, assembly, and annotation of *Nix* in various species is summarized in Figure 1. Further elaborating, tBLASTn was first performed using all known Nix peptides under very low stringency (evalue 10). For example, to retrieve *Nix*-related cDNA sequences for *T. amboinensis*, larvae RNAseq data were used as the database. All *Nix*-related reads and their pairs were retrieved and assembled using Trinity 2.8.5 with default parameters (Grabherr, et al. 2011). The resulting trinity assemblies were used as queries to perform a BLASTx against a dataset of diverse RNA-binding proteins (RBPs) to remove non-*Nix* sequences that better match other related proteins. Similarly, to retrieve *Nix*-related genomic DNA sequences, male *T. amboinensis* WGS data were used as the database in a tBLASTn search with known Nix peptides as the query. All *Nix*-related gDNA reads and their pairs were retrieved and assembled using Trinity. The resulting gDNA trinity assemblies were used as queries to perform a BLASTx against diverse RBPs to remove non-*Nix* sequences. The two resulting *Nix* genomic DNA fragments were separated by a large intron. Therefore, a string of Ns was added between the two sequences to indicate the incomplete intronic sequence. Comparison between the *T. amboinensis Nix* cDNA and the gDNA sequences confirmed/defined the exon-intron boundaries. In addition to *T. amboinensis*, we generated RNAseq or cDNA data for *Ae. atropalpus, Ae. triseriatus,* and *Psorophora columbiae* (Table 1). In the case of *Ae. atropalpus,* two splice isoforms exist and they were subsequently confirmed by sequencing of RT-PCR products (described below). We also downloaded transcriptomic data for either male or mixed-sex samples of *Ae. detritus*, *Toxorhynchites sp*. (identification only at the genus level), and *W. smithii* for *Nix* identification (Table 1). For all remaining species, *Nix* was characterized using male WGS. Protein sequences for *Nix* genes lacking cDNA data were predicted using FGENESH+ (Solovyev 2007) using existing Nix protein sequences as the guide. All *Nix* nucleotide and protein sequences are provided in the Supplemental Data 1.

### Assessing male-specificty of Nix by chromosomal quotient (CQ)

For the eight species in which both male and female WGS data were available, CQ was performed using the *Nix* ORF (Table 2) as previously described (Hall, et al. 2013; Matthews, et al. 2018). Briefly, male and female Illumina WGS data were aligned to *Nix* ORFs using bowtie2 (https://bowtie-bio.sourceforge.net/bowtie2/index.shtml) under default conditions and CQ was calculated as the number of female hits divided by the number of male hits. In some species, only one end of the paired reads was used for each sex.

### Phylogenetic Inference

Multiple sequence alignments (Figures S2 and S6) using amino acid sequences (Supplemental Data 1) were generated with MegAlign Pro®. Version 17.2.1. DNASTAR. Madison, WI, using the MAAFT algorithm. Prior to phylogenetic inference, nonhomologous sequence in the N-terminal region of Transformer-2 sequences was removed from the alignment. Phylogenetic inference using Maximum Likelihood was performed with 1000 bootstrap replicates using RAxML (Stamatakis 2014) as a feature of MegAlign Pro®(Version 17.2.1. DNASTAR. Madison, WI). Phylogenetic inference was performed using MrBayes 3.2.7a x86_64 (Huelsenbeck and Ronquist 2001). Default parameters and the “mixed amino acid” model were implemented for 1M generations, where the WAG amino acid substitution model (Whelan and Goldman 2001) was determined to have a posterior probability of 1.0, and minimum and maximum probabilities of 1.0 from 2 independent runs. BIONJ was performed using Phylogeny.fr (http://www.phylogeny.fr/index.cgi) with 1000 bootstrap replicates and default parameters (Dereeper, et al. 2008). See Supplemental Data 1 for sequences and alignments.

### Plasmids Constructs

Plasmids containing the *Nix* ORF from 3 species (*Ae. polynesiensis*, *Ae. vexans*, and *Ae. japonicus*) (Table 3, see Supplementary Data S2 for plasmid sequences) were designed for *piggyBac*-mediated transformation and generation of mRNA for embryonic injection in *Ae. aegypti*. Each *Nix* ORF was expressed using the UTR’s and promoter/upstream sequence from *Ae. aegypti* (Aryan 2020). ORFs were synthesized and cloned into *piggyBac* donor backbone plasmids (Horn and Wimmer 2000) by Epoch Life Sciences (Missouri City, TX).

### Mosquito rearing

*Ae. aegypti* (Liverpool strain) was maintained at 28°C and 60-70% humidity, with a 14/10 hour day/night light cycle. Adult mosquitoes were maintained on 10% sucrose and blood-fed using artificial membrane feeders and defibrinated sheep’s blood (Colorado Serum Company; Denver, CO).

### PiggyBac*-*mediated transformation

Donor plasmids were co-injected at 0.5 μg/μl with the *piggyBac* mRNA helper at 0.3 μg/μl into 1h old embryos (Handler, et al. 1998). Surviving G_0_ females were mated to Liverpool males in pools of 20-25. G_0_ males were mated individually to 5 Liverpool females and mosquitoes from 15-20 of these cages were merged into one large pool. G_1_ larvae were screened for GFP fluorescence using a Leica M165 FC fluorescence microscope. Positive G_1_ individuals were out-crossed to Liverpool females to ensure all transgene cassettes were stably inherited to the G_2_ generation. Three *piggyBac* based donor constructs were used: the 187_Ae. japonicus, 188_Ae. polynesiensis, and 189_Ae. vexans were injected into *Ae. aegypti* embryos (Liverpool) with *piggyBac* mRNA helper. Transgenic individuals were identified in four 187_Ae. japonicus, seven 188_ Ae. polynesiensis, and seven 189_Ae. vexans pools. In the 189_Ae. vexans experiment, a range of phenotypes were observed including flightless intersex and flightless females.

### PCR on Ae. atropalpus gDNA

Genomic DNA was extracted using Zymo Quick-gDNA Miniprep Plus kit. PCR was performed with the following reaction in total volume of 20 ul: 1X Phire reaction buffer, 200uM dNTPs, 0.5uM of primers, 0.4 ul of Phire Hot Start II DNA Polymerase (Thermo Scientific), and 10 ng of gDNA. The PCR cycling condition is as following: 98C 30 sec, followed by 30 cycles of 98C 5 sec, 63C 5 sec, and 72C 15 sec, finally 72C 10 min.

### RT-PCR

*Ae. aegypti* adult male and females were collected and flash frozen. RNA was extracted following the total RNA extraction protocol for the Quick-RNA Miniprep kit (Zymo Research, Irvine, CA) and RNA samples were treated by DNase I. cDNA synthesis was performed using SuperScript III RT first strand cDNA synthesis kit (Invitrogen). The amount of starting RNA in each reverse transcription reaction was 2 ug. The final cDNA product was diluted 1:3 with nuclease-free water and stored at −20°C. For RT-PCR of *Ae.vex.Nix* samples*, AevexNix* and RPS7 (reference gene) was performed using Phire II DNA polymerase (Thermo Fisher Scientific, Waltham Massachusetts). The cycling conditions for vexNix and RPS7 reactions were as follows: 98°C for 30 seconds; 33 cycles of 98°C for 5 seconds, 55°C for 5 seconds, and 72°C for 15 seconds; and final extension at 72°C for 1 minutes. For RT-PCR of *Ae. atropalpus Nix samples,* approximately 7 male and 7 female pupae were used to isolate total RNA and cDNA was made as described above. PCR was performed with the following reaction in total volume of 20 ul: 1X Phire reaction buffer, 200uM dNTPs, 0.5uM of primers, 0.4 ul of Phire Hot Start II DNA Polymerase (Thermo Scientific), and 10 ng of cDNA. The PCR cycling condition is as following: 98C 30 sec, followed by 35 cycles of 98°C 5 sec, 55°C 10 sec, and 72°C 20 sec, finally 72°C 10 min. PCR products amplified with primers Aeatro-Nix-F1/R1 were purified with GFX PCR DNA and Gel Band Purification Kit (GE Healthcare) and cloned into pJET1.2/blunt cloning vector. One positive clone from each colony was selected and sequenced. PCR products amplified with primers Aeatro-Nix-F2/R2 were purified with GFX PCR DNA and Gel Band Purification Kit (GE Healthcare) and directly sequenced.

### RT-qPCR

Target gene and AAEL002401(reference gene) expression was quantified by RT-qPCR using FAM- and HEX -labelled GoTaq® Probe qPCR and RT-qPCR assay (Promega), TaqMan universal PCR master mix (Promega) and the CFX96 Touch Real-Time PCR Detection System (BioRad Laboratories, Hercules, CA, USA). Samples were assayed in triplicates and Ct values were automatically generated by the BioRad CFX Manager 3.1 software. A Ct = 40 corresponding to the final RT-qPCR cycle was assigned to samples where no Ct value was obtained. Relative gene expression was calculated as ΔCt (*vexNix* Ct – AAEL002401 Ct). The cycling conditions reactions were as follows: 95°C for 3 minutes; 39 cycles of 95°C for 10 seconds, 60°C for 30 seconds.

### Droplet Digital PCR (ddPCR)

ddPCR for target gene and AAEL002401 (reference gene) were performed with two probe and primer mixes were used to carry out TaqMan assays respectively. The two probes are labeled with different fluorescent dyes (Target-FAM, and AAEL002401-HEX) to allow detection of both in one reaction droplet ddPCR was performed with a BioRad QX100 ddPCR machine using the recommended protocols and reagents (BioRad Labatories, Hercules, CA). ddPCR was performed on biological quadruplicates of males and females. The cycling conditions reactions were as follows: 95°C for 10 minutes; 40 cycles of 94°C for 30 seconds, 59°C for 1 minute, and final at 98°C for 10 minutes.

### Statistics

Data were analyzed and associated graphs generated using GraphPad Prism (v 9.3.1). Statistically significant differences between expression data were analyzed using One-way ANOVA followed by the Tukey’s Multiple Comparison Test.

### Nix protein structure predictions

To predict *Nix* and *fle* protein structures, a modified AlphaFold2 (Jumper, et al. 2021), called AlphaFold2 using MMSeqs2 (Mirdita, et al. 2022), was used via a Jupyter notebook inside Google Collaboratory (https://colab.research.google.com/github/sokrypton/ColabFold/blob/main/AlphaFold2.ipynb).

Structure models with the highest average predicted Local Distance Difference Test (pLDDT) scores (Model 1) were selected for structure images by exporting PDB files. 3-D structure images were generated using iCn3D (Wang, et al. 2020; Wang, et al. 2022) (https://www.ncbi.nlm.nih.gov/Structure/icn3d/full.html).

### Primers and probes list

Supplemental Table S3

## Data Availability Statement

Strains and plasmids are available upon request. Sequencing data are deposited and available at https://www.ncbi.nlm.nih.gov/sra/PRJNA885905.

## Supporting information

Combined Supplemental Info and Supplemental Data

## Acknowledgements

Acknowledgements and funding

This work is supported by NIH grants R01AI123338 and 1R01AI157491 and the Virginia Agriculture Experimental Station. We thank Jeff Powell at Yale University and Daniel Dixon at the Anastasia Mosquito Control District (St. Augustine, FL, USA) for kindly providing the *Aedes mascarensis* and *Psorophora columbiae* samples.

