## Supplementary material for "On the origin and evolution of the mosquito male-determining factor *Nix*": Combined Supplemental Info and Supplemental Data

### Supplemental Information

**Supplemental data 1. Nucleotide and protein sequences of all newly discovered *Nix*, other sequences used in this study, alignments used for phylogenetic analysis**

**Supplemental data 2. Plasmid sequences**

**Table S1. Detailed information for sequencing samples attached at the end of Supplemental Information**

**Table S2. Detailed phenotype description/record for 189_*Ae. vexans Nix* lines**

| | **G1 positives from pools** | | | | | | |
|---|---|---|---|---|---|---|---|
| **Phenotypes** | **P4** | **P8** | **P10** | **P11** | **P1** | **P2** | **P6** |
| (+) male | 5 | 2 | 4 | 44 | these pools had only flightless females which cannot mate so these lines were terminated | | |
| (+) intersex flightless | 2 | 4 | 1 | 36 | | | |
| (+) female flightless | 6 | 0 | 0 | 0 | | | |
| (+) female | 2 | 0 | 1 | 0 | | | |

Negative males and females were not counted

**Table S3. Primers and probes**

ddPCR/RT-qPCR

Gene AAEL002401 (internal reference)

| | |
|---|---|
| Probe HEX | CGTATTGGTTGGAGGCTATGACGA |
| Forward Primer | TACAAGATGCGCAATGGATA |
| Reverse Primer | TGGCCAGATAGTCGATGTAAT |

Ae. aeg. Nix native

| | |
|---|---|
| Probe FAM | CGTGCAAATGTGTAAAAAAAGAAATGC |
| Forward Primer | GATGTGATCTTTTTCAAAGAAAAT |
| Reverse Primer | GATGCAAAGAATGGAATATTTC |

Ae. pol. Nix

| | |
|---|---|
| Probe FAM | AGCACATCTCTGCTCAAGCTGCC |
| Forward Primer | TACGACTCTACCGGACACTCT |
| Reverse Primer | ATCACTGCGGTCCATTTCCT |

Ae. jpn. Nix

| | |
|---|---|
| Probe FAM | TGCCAAACATTTTTCCAAGTATGCACCC |
| Forward Primer | GGGCTCTCCAAAGAGACAACA |
| Reverse Primer | GCGGATTGTGGATCGTCGAA |

Ae. vex. Nix

| | |
|---|---|
| Probe FAM | CCATCTGATACCAAGGAAGCA |

| | |
|---|---|
| Forward Primer | CAGTACCATCGGCATATTTG |
| Reverse Primer | TCGAGTTTCCACCTTTGTC |

Ae. aegypti DsxF

| | |
|---|---|
| Probe FAM | TGACGAAGGTCAAGCCGTG |
| Forward Primer | GAACTTGTCAAACGATCTCAATG |
| Reverse Primer | ATGTTCAGATTGTGCAATCG |

Ae. aegypti DsxM

| | |
|---|---|
| Probe FAM | CGGATTGACGAAGGATACGACATT |
| Forward Primer | GATACCCCTGGGAGATGATG |
| Reverse Primer | TGGAACGCTTCGGAAAGTAG |

Ae. aegypti RPS7

| | |
|---|---|
| Forward Primer | ATGGTTTTCGGATCAAAGG |
| Reverse Primer | CTTGTGTTCAATGGTGGTCTG |

*Ae. atropalpus* RT-PCR

| Primer name | Sequence | Location |
|---|---|---|
| Aeatro-Nix-F1 | AGTTCGGTTCAGTCGCTTGAT | Exon 1 |
| Aeatro-Nix-R1 | CTGTGCGCTTGCTTTTGTGT | Exon 2 |
| Aeatro-Nix-F2 | TTACACAAAAGCAAGCGCACA | Exon 2 |
| Aeatro-Nix-R2 | CTTTTTGAGCCATCCGAGCTG | Exon 3 |

*Ae. vexans* transgenic RT-PCR

| | |
|---|---|
| Forward Primer | GAAATGTGATTTTTGTAATATAC |
| Reverse Primer | CGATATCATCAGTCAATACTAATAGT |

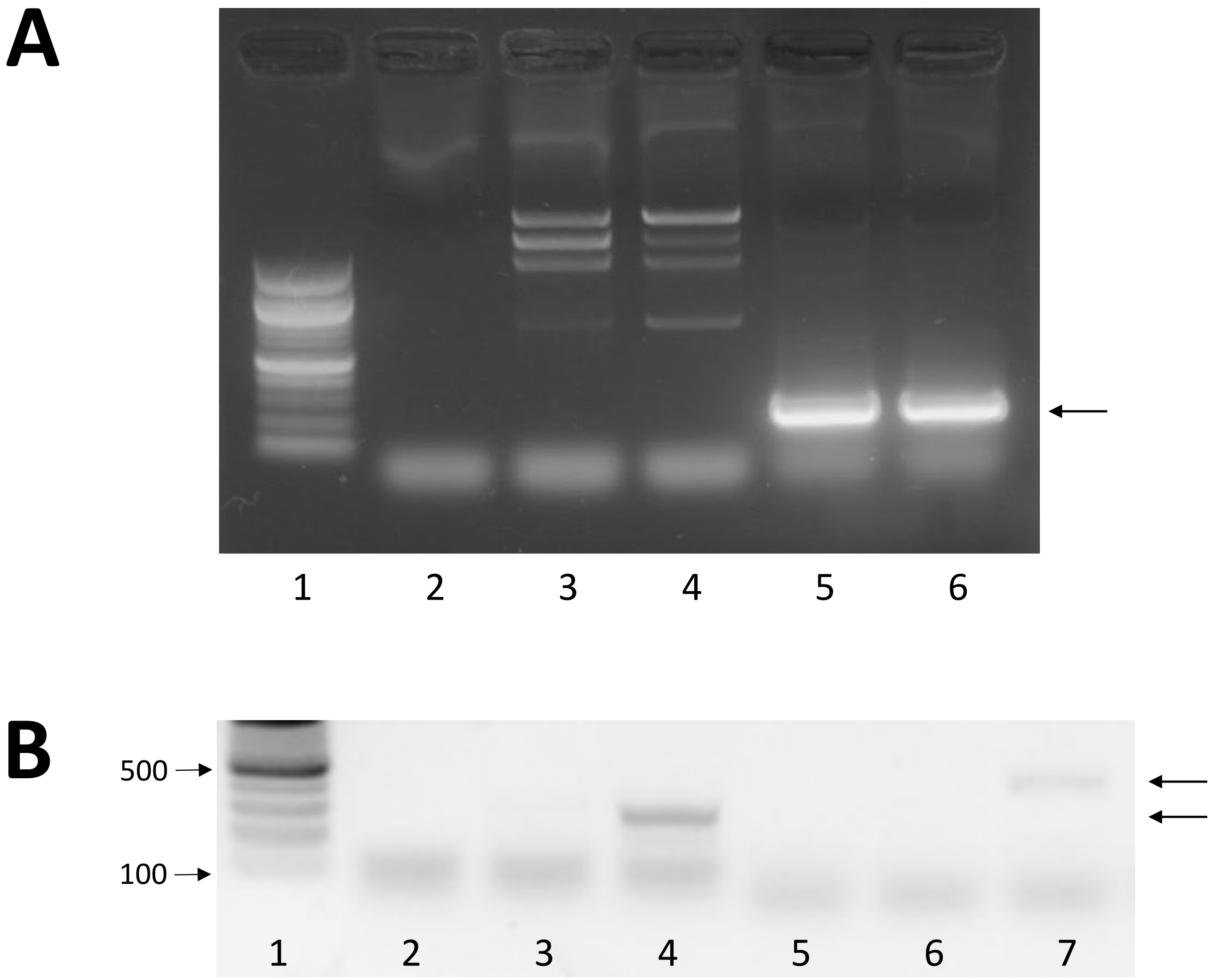

**Figure S1. Male-specific inheritance and expression of *Ae. atropalpus Nix*.** A) PCR performed on *Ae. atropalpus* genomic DNA confirms male-specificity of the *Nix* gene. The expected amplicon size of 246 bp was obtained from primers Aeatro-Nix-F1 and Aeatro-Nix-R1 that span exons 1 and 2. Genomic DNA from one adult was used for each sample. Samples: 1, 100 bp DNA marker; 2, negative control (H2O); 3, female; 4, female; 5, male; 6, male. B) RT-PCR was performed using cDNA made from RNA isolated from *Ae. atropalpus* pupae. Samples: 1, 100 bp DNA marker; 2, Negative control; 3, Female cDNA; 4, Male cDNA; 5, Negative control; 6, Female cDNA; 7, Male cDNA. Lanes 2, 3, 4: PCR amplicon (246 bp expected) from primers Aeatro-Nix-F1 and Aeatro-Nix-R1 that span exons 1 and 2. Lanes 5, 6, 7: PCR amplicon (335 bp expected) from primers Aeatro-Nix-F2 and Aeatro-Nix-R2 that span exons 2 and 3. Each sample cDNA was generated from RNA isolated from approximately 7 pooled pupae. Arrows indicate bands of expected size.

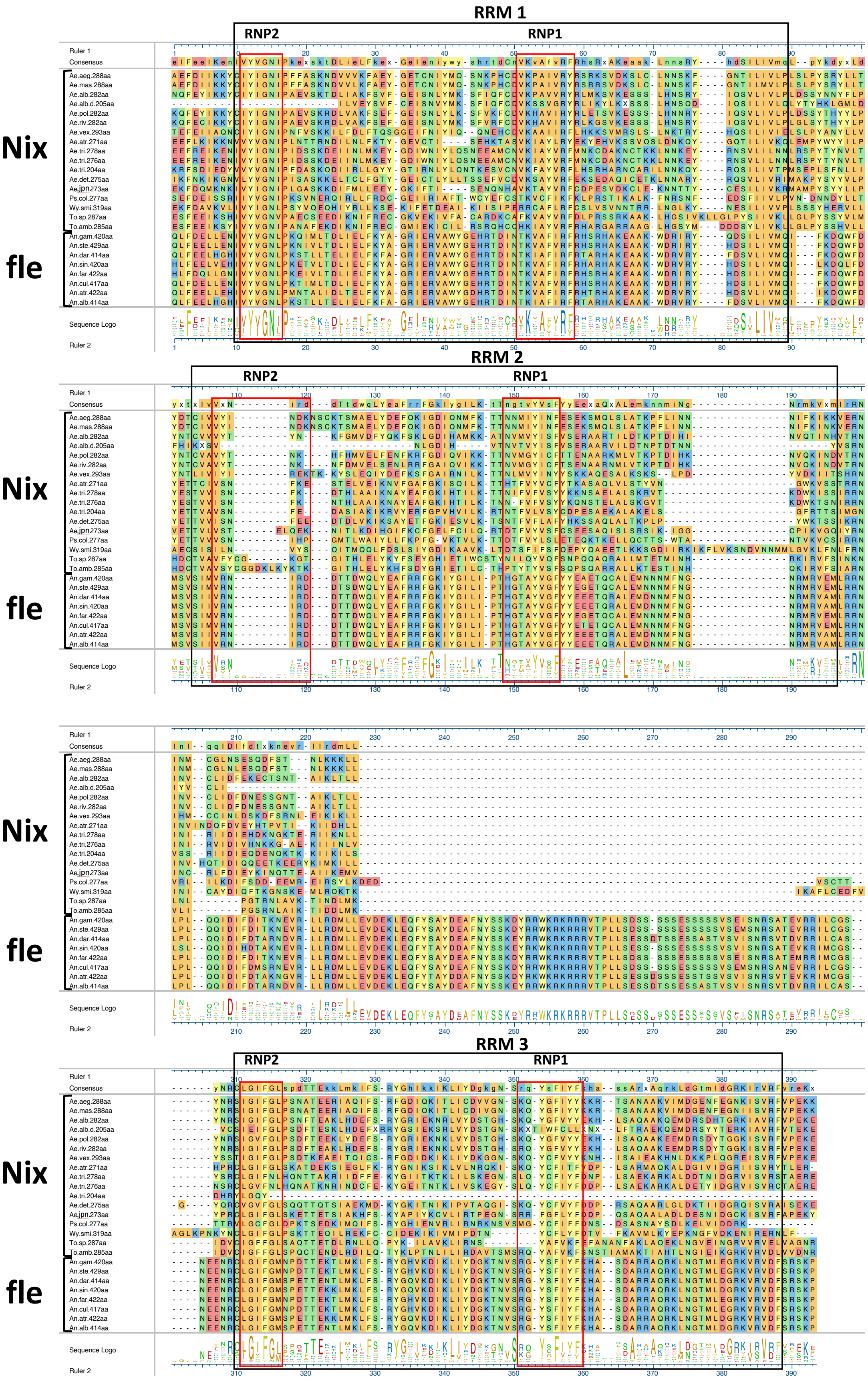
RRM 1
RNP2
RNP1
RRM 2
RNP2
RNP1
RRM 3
RNP2
RNP1
Nix
fle
Ruler 1
Consensus
Sequence Logo
Ruler 2
Ae.aeg.288aa
Ae.mas.288aa
Ae.alb.282aa
Ae.alb.d.205aa
Ae.pol.282aa
Ae.riv.282aa
Ae.vex.293aa
Ae.atr.271aa
Ae.tri.278aa
Ae.tri.276aa
Ae.tri.204aa
Ae.det.275aa
Ae.jpn.273aa
Ps.col.277aa
Wy.smi.319aa
To.sp.287aa
To.amb.285aa
An.gam.420aa
An.ste.429aa
An.dar.414aa
An.sin.420aa
An.far.422aa
An.cul.417aa
An.atr.422aa
An.alb.414aa

**Figure S2. Multiple sequence alignment of full-length *Nix* and *fle* sequences used for phylogenetic inference in Figure 4.** Alignment was produced by MegAlignPro (see Methods). Non-aligning N-terminal and C-terminal ends were trimmed. See Supplemental Data 1 for full-length sequences. Black boxes surround conserved RNA Recognition Motifs (RRMs). Red boxes surround conserved motifs RNP2 and RNP1.

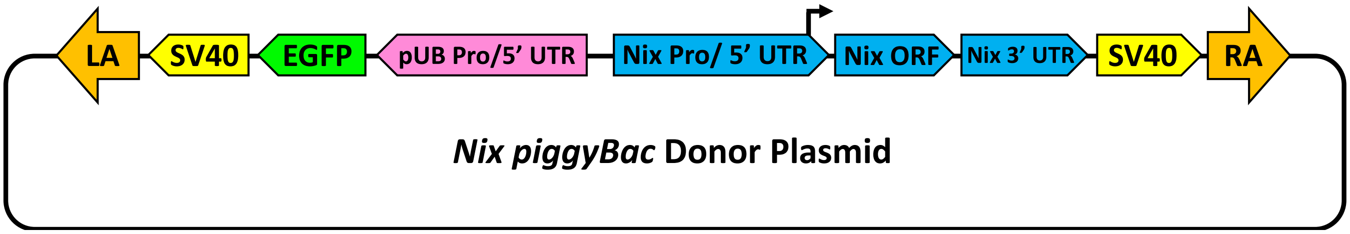

**Figure S3. *piggyBac* Donor Plasmid for heterologous *Nix* expression.** Schematic shows *piggyBac* donor plasmid used for transformation and expression of *Nix* ORFs. "LA" and "RA" indicate *piggyBac* Right and Left Arms, respectively; SV40, SV40 polyadenylation signal; pUB Pro/5' UTR, *Ae. aegypti* polyubiquitin promoter and 5' UTR; Nix Pro/ 5' UTR, *Ae. aegypti Nix* promoter and 5' UTR.

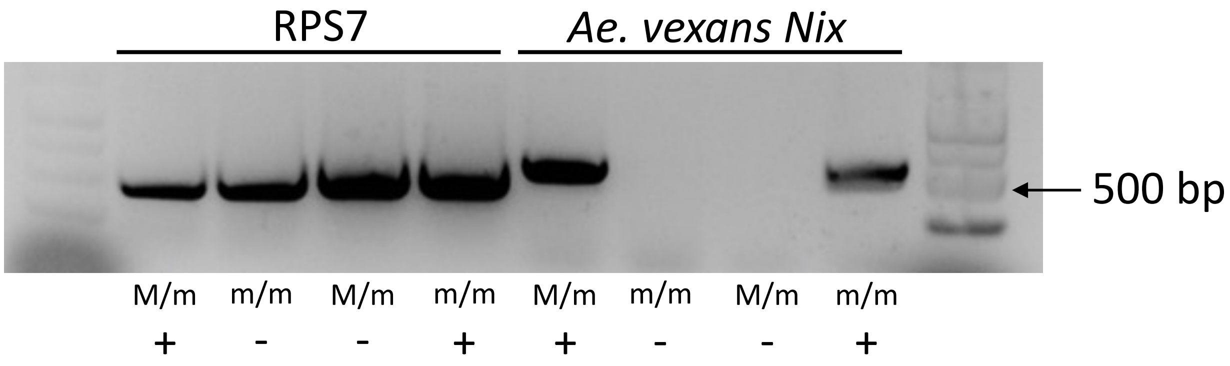

**Figure S4. RT-PCR verifies heterologous expression of *Ae. vexans Nix* in transgenic *Ae. aegypti* (Line *Ae.vex.p11*) individuals.** RT-PCR was performed using the same cDNA as in A) but using a single individual for each genotype. The expected 524 bp amplicon from the transgenic *Ae. vexans Nix* ORF was detected only in transgenic individuals. RPS7 was used as a control. Genotypic sex and presence/absence (+/-) of the *Ae. vexans Nix* transgenic cassette are indicated at bottom.

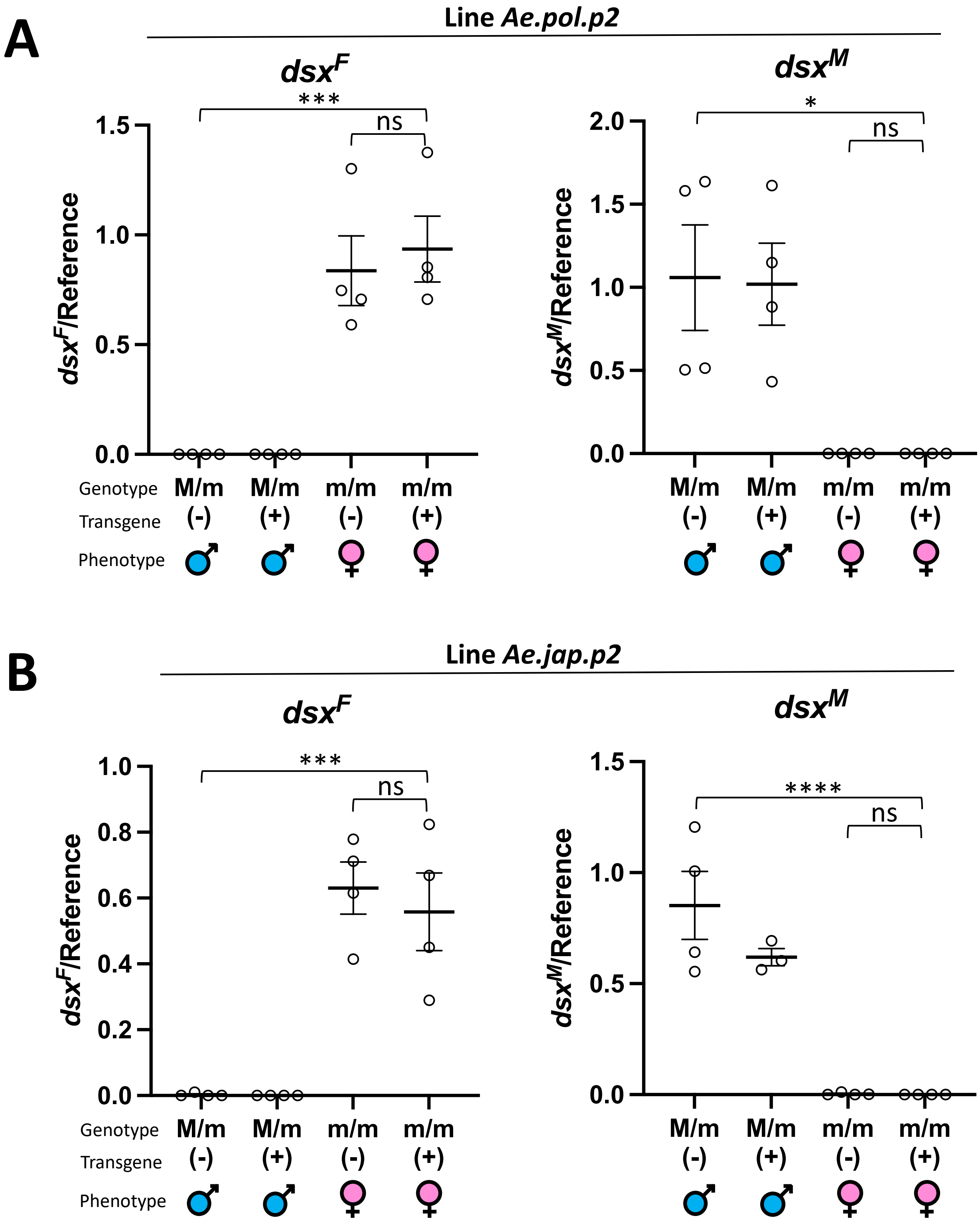

**Figure S5. Expression of female and male *dsx* isoforms (*dsx*$^{F}$ and *dsx*$^{M}$) was determined in lines *Ae.pol.p2* (A) and *Ae.jpn.p2* (B) by RT-qPCR relative to an endogenous gene AAEL002401 used as a control.** Adult progeny with 4 resulting genotypes from a cross of transgenic males and wild-type females were assayed. X-axis labels: genotypic sex (male, M/m; female; m/m); (+)/(-) indicates presence/absence of the *Ae. vexans Nix* transgenic cassette determined by a fluorescent marker; symbols indicate phenotypic sex (male, female). Individual values are shown with the mean and +/- SEM. Statistically significant differences were determined according to One-way ANOVA followed by the Tukey's Multiple Comparison Test.

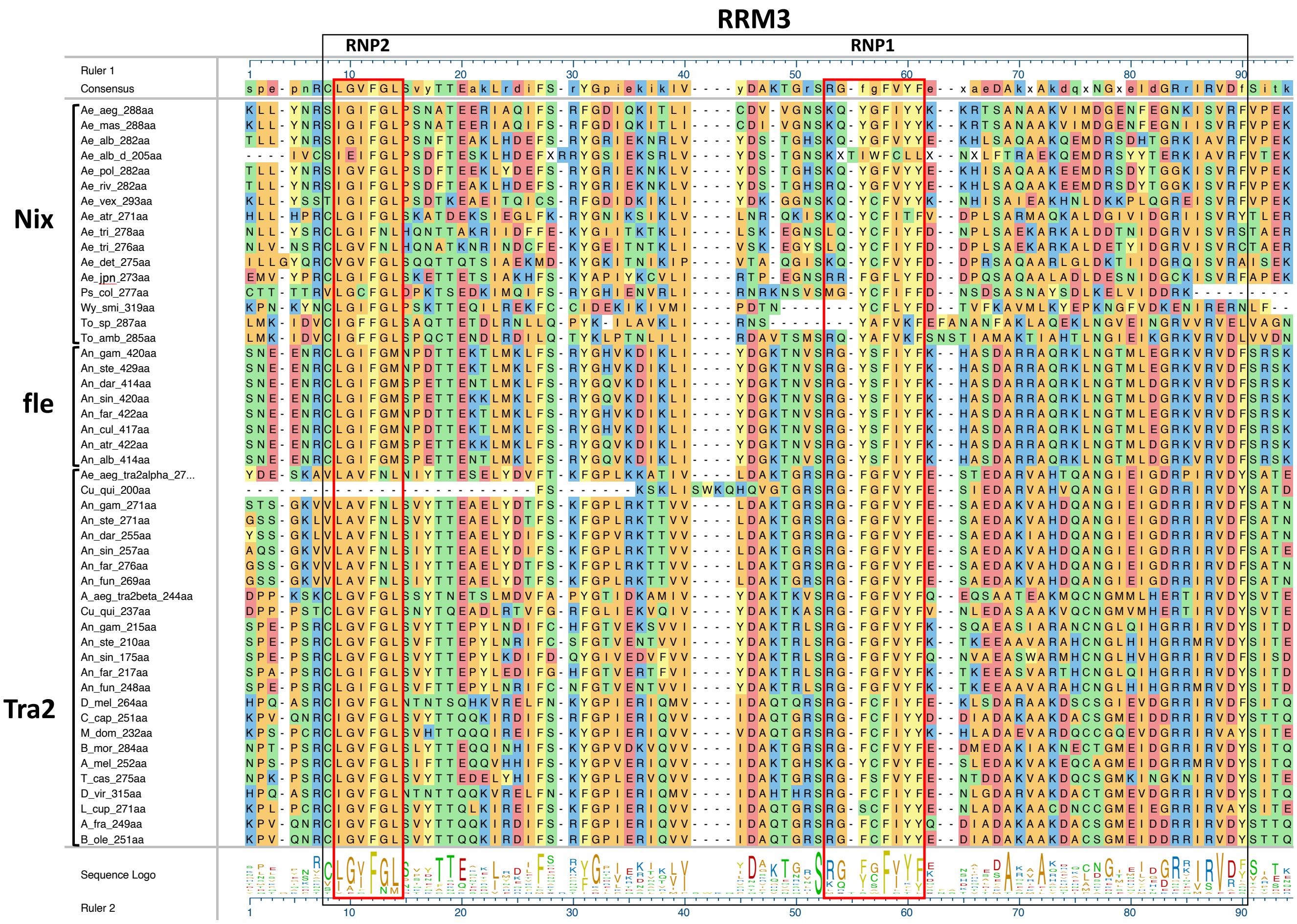

**Figure S6. RRM3 of *Nix* and *fle* sequences aligned with the single RRM of *tra2* sequences.** This multiple sequence alignment was used for phylogenetic inference in Figure 6. Screenshots are from MegAlign Pro®(Version 17.2.1. DNASTAR. Madison, WI), and are annotated to show RNA recognition motifs (RRM), and motifs RNP1 and RNP2 are (boxed in red).

**A**

RRM3 Maximum likelihood Reference Tree for Taxa

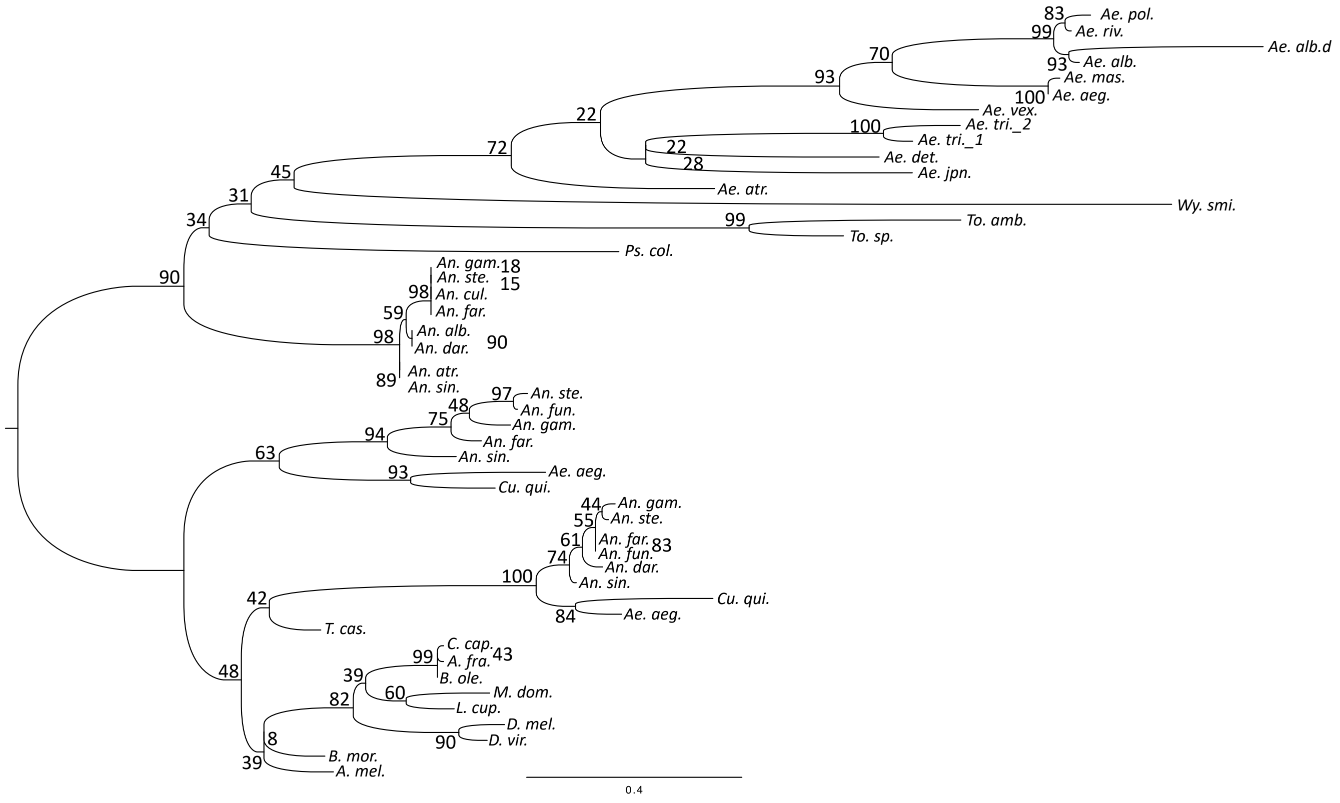

# B

RRM3 MrBayes Reference Tree for Taxa

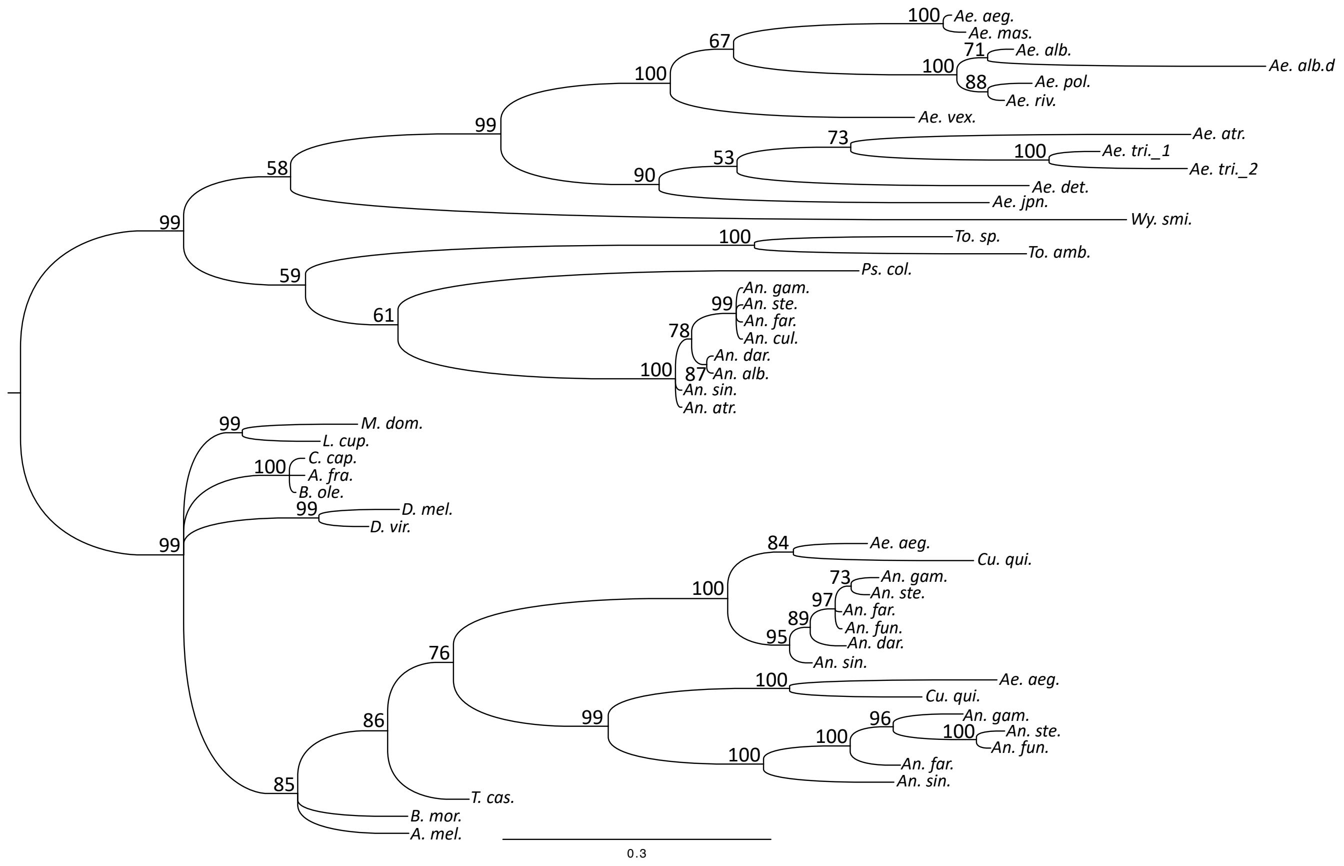

# C

RRM3 BIONJ Reference Tree for Taxa

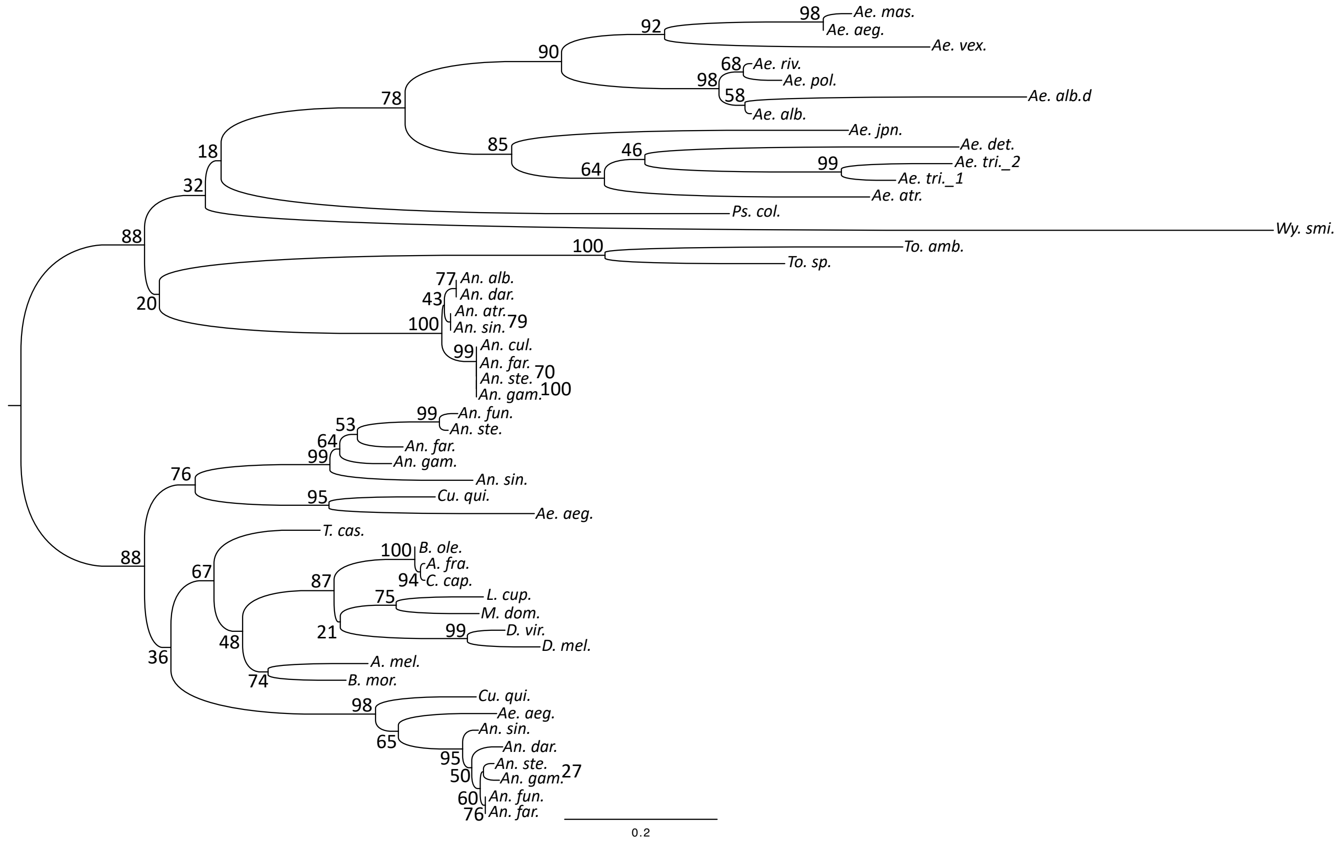

**Figure S7. Reference trees for phylogenies in Figure 8 showing all taxa and clade credibility values.** A) Maximum likelihood B) MrBayes C) BIONJ.

### Table S1. Detailed information for sequencing samples

| | | | | | |
|---|---|---|---|---|---|
| ***The following are genomic DNA sequence samples*** | | | | | |
| Species | Source | Strain or Other Metadata | Sex | Stage | Numbers |
| *Aedes mascarensis* | Jeff Powell, Yale | Colonied by Powell | Male | adult | >20 |
| *Aedes mascarensis* | Jeff Powell, Yale | Colonied by Powell | Female | adult | >20 |
| *Aedes polynesiensis* | Stephen Dobson, Univ. Kentucky | | Male | adult | >20 |
| *Aedes polynesiensis* | Stephen Dobson, Univ. Kentucky | | Female | adult | >20 |
| *Aedes riversi* | Stephen Dobson, Univ. Kentucky | | Male | adult | >20 |
| *Aedes riversi* | Stephen Dobson, Univ. Kentucky | | Female | adult | >20 |
| *Aedes vexans* | Daniel A. Hartman, Rebekah Kading and Greg Ebel, Colorado State University | Collected 8/23/2016 by CDC light trap, LaSalle, Colorado | Male | adult | 5 |
| *Aedes vexans* | Daniel A. Hartman, Rebekah Kading and Greg Ebel, Colorado State University | Collected 7/20/2017 by CDC light trap, LaSalle, Colorado | Female | adult | >20 |
| *Aedes japonicus* | Sally Paulson and Fan Yang, Virginia Tech | Eggs were from female mosquitoes collected using gravid traps in forested areas around Blacksburg, VA. All adult mosquitoes were tested for virus to ensure the absence of virus in the F1 progeny. They were then maintained in the lab using standard rearing conditions | Male | adult | 5 |
| *Aedes triseriatus* | Sally Paulson and Fan Yang, Virginia Tech | same as above | Male | adult | >20 |
| *Aedes triseriatus* | Sally Paulson and Fan Yang, Virginia Tech | same as above | Female | adult | >20 |
| *Aedes atropalpus* | Mike Strand, Univ. Georgia | | Male | adult | >20 |
| *Aedes atropalpus* | Mike Strand, Univ. Georgia | | Female | adult | >20 |
| *Psorophora columbiae* | Daniel Dixon; Rudy Xue, Anastasia Mosquito Control District, Florida | | male | adult | >20 |
| *Psorophora columbiae* | Daniel Dixon; Rudy Xue, Anastasia Mosquito Control District, Florida | | female | adult | >20 |
| *Toxorhynchites amboinensis* | Mike Strand, Univ. Georgia | | Male | adult | 1 |
| *Toxorhynchites amboinensis* | Mike Strand, Univ. Georgia | | Female | adult | 1 |
| ***The following are RNAseq datasets*** | | | | | |
| *Aedes triseriatus* | Sally Paulson, Virginia Tech | See above | Male | adult | 15 |
| *Aedes atropalpus* | Mike Strand, Univ. Georgia | See above | mixed | embryo, 2-12 hr | >100 |
| *Aedes atropalpus* | Mike Strand, Univ. Georgia | See above | mixed | embryo, 2-12 hr | >100 |
| *Aedes atropalpus* | Mike Strand, Univ. Georgia | See above | mixed | embryo, 12-24 hr | >100 |
| *Aedes atropalpus* | Mike Strand, Univ. Georgia | See above | mixed | embryo, 24-36 hr | >100 |
| *Psorophora columbiae* | Daniel Dixon; Rudy Xue, Anastasia Mosquito Control District, Florida | See above | male | adult | 6 |
| *Toxorhynchites amboinensis* | Mike Strand, Univ. Georgia | See above | mixed | L3 instar | 3x8 |

**Supplemental Data 1. Sequences used in this study**

**This file contains:**
A) Predicted *Nix* sequences
B) Other sequences used in this study
C) Alignments used for phylogenetic analysis

---

#### A) Predicted *Nix* sequences

See Supplemental Table S1 for description of the genomic and RNAseq datasets.

Predicted exons and introns of Nix for several species are shown below
Introns are underlined
nnn... indicates undetermined genomic sequence between two Trinity assemblies
ATG indicates start codons
TAA and UAG indicate stop codons

```
>Ae.mascarensis
ATAGTAAAAATAAGCACGGCTGTGTAGTTAAACATACTACTTCTTTTTTATCAGTGTATGAGTATAAAGAACTTATTTTTTGG
GTAGTCTGGTTTCTCCGTGTAGTCGCGAATGCTTATAGTAAATAGTGACGTAACCATTTGCATTGCATAATCGAGTAAAAAAT
GAGAGCAGTGGCACGCAACATGAGTTTTTCACCTGAGCAAGTTAAGAACATTGACTACGGAGCTCAGTCCAGCAATAATGTTA
ATCATAAAAAGATGTGATCTTTTTTAAAGAAAATGTAATTAGTGTTTTCTAAGTTTTATATTGATTTTTTTTTGTCGTCGTGC
AAATGTGTAAAAAAAGAAATGCTGAGATCAATGCAGAATTTGACATCATCAAAAAATATTGTATATATATAGGAAATATTCCA
TTCTTTGCATCAAAGAATGATGTGGTTCTCAAATTTGCTGAATACGGCGAAACATGTAACATATATATGCAATCAAATAAACC
ACATTGCGACGTTAAACCAGCAATTGTTCGATATCGTTCGAGAAGAAGTGTAGACAAGTCTTTGTGTTTAAACAATTCAAAAT
TTGGCAACACAATATTAATGGTTCTACCACTAAGTTTGCCTTATTCAAGATATCTGCTAACCTATGATACTTGCATTGTGGTA
TATATTAATGATAAGAATTCTTGTAAGACATCAATGGCTGAGCTATATGATGAATTTCAAAAAATTGGTGATATTCAAAACAT
GTTTAAAACGACAAATAATATGATTTATATAAACTTTGAGTCTGAAAAGTCTATGCAATTATCTTTAGCTACTAAGCCCTTTT
TAATAAATAATAATATTTTCAAAATCAACAAAGTTGAACGAAACATTAATATGTGCGGTCTAAATTTAGAAAGCCAAGATTTC
TCCACAAATCTAAAAAAAAAACTTTTGTATAATCGCTCCATTGGAATATTTGGATTGCCATCAAATGCCACGGAAGAGCGAAT
TGCACAAATATTTTCAAGGTAAGTATGTGCAAATTAACGCGTGCTTCAGAATAATTACGGCGCTCATATTAAATAATAGAAAT
ATTAGTAGTAGAACGCTAGTGGCGCTGTTGTCACAAAACTGATTTGACTAATTATTACACTTAATTGCTATTTT

nnnnnnnnnn

CATTATGATCATCATTATCTAAACTTCTAAAGTAGTTTTTTCTGTTCAAAACTTAAAATTATTCGATGAAAACGTGTTTACTT
AGTTTCGGTTGGAGAATAGAATACAGTGAACACATTTTCATGTGGAGTTTCTTGATGACGGAAACTTTTTTTCTTGTACGAGC
CATACATTCTTTTGTTCTTTTGTCTCTTACTGTGCGCCGTGTGTAGATGCTGTCTTGCTCTCTCACGGGAAATTCAAAAACAG
CGTGCACAGTAAAAGTGAAACGTTTTATAGCATGCTTAGGCTCATAGCCTCATTTAATCTTAGGAGTATTTGATTTAAATTTA
TTTATAAAAAGACAATAATTTATTTACAAAAATCATTTGTTCGTTCCAGTGAAGCTTTAATGCTTAATTTATGCTCTTCCAGG
TTTGGCGACATTCAAAAAATTACATTGATATGCGACATAGTCGGCAACTCGAAGCAATATGGTTTTATATATTACAAGAAACG
TACTTCTGCTAATGCTGCTAAAGTGATAATGGATGGAGAAAATTTTGAAGGAAATATAATTTCCGTTCGTTTTGTCCCAGAAA
AAAAAGTGTTTAAAAATTAATAAAGTGATATATGACACACATTTGTTTTTCTTTTGTCTAAGACTGAGTACATACGCCCTAAA
AATATAAAATATAATATTTAGAATGTAGAATGTAATGAAAAAAGATAACACCTGGCACCGTAAAATGTCAAAAAAGTGGACTT
CTGATTTCATTTACTATCGTTCTTGCTTGACCCAATCTCACCCCAATTTTTAATGTGTTAGACACCGTACCGAAAA

>Ae.polynesiensis
TGTTTTGAGCCGTTTTCAAGGTTGGTATTCACCCAACCAAAATACATGGTTAATTTTCCCCAACAATTTTGCTGATTCTACCT
AAGATTTAGTTTGAAATTGTTAATATGATGGACTATTCTTCAACCTCTTTTTTGTTAAAGGCAAACGTTCCAATGGTAAGAAA
TAGCTGTAATCTTAGTTTGAATTTCATAAATTTAGATTTGGAAGAACCAAACTATTTAGGTAGAAAATACCTAAATATAGCGT
CTGTTTTAACTGCAGCCGCAATGCAGCATTTAACAATTTTTGACTTTGGAGTAAACAAAAATTGACCTTGATTGCCAACCATA
ATTTTAGTTGATACAAACTAGTAACATTTCTCTCCGTGTATATTTGCGTAATGCGTATCCAAATGTGAGTAACATATACCCAT
ATATTTAGAAAAATAGGTAAACTGATCTTAGCGTGTAGGTCGTACACATTTTTGTGCTCGGAATATTAAGTGTGCTCATCACA
```

```
CTGCTTTTATTTTTGGCACGATTTATTATTTTTTCGTAATGTACAGCAAAAATGAGATCAATCTCATTAATAAGCAATTTGAA
TATATTAAAAAATATTGCATATATATTGGAAACATTCCTGCCGAAGTGTCAAAAAGAGATTTAGTTGCAAAATTTTCCGAATT
CGGGGAAATATCCAACTTATATTTGAAGTCATTCGTAAAGTTCTGTGATGTGAAACATGCTGTAATTCGTTACAGATTGGAAA
CAAGTGTAAAGGAATCTTCAAGTTTACACAATAATCGATATATTCAATCGGTTTTAATAGTTCTGCCACTAGACTCGTCGTAC
ACCCATTACTATCTTCCTTACAACACTTGTGTTGCGGTGTACACTAACAAACATTTTCATATGGTAGAACTTTTTGAAAATTT
TAAAAGATTTGGAGACATTCAAGTCATAAAGAAAACTACAAACGTCATGGGATACATTTGTTTTACAACAGAAAATGCTGCAA
GAAAAATGTTGGTTACTAAGCCCACAGATATCCATAAAAATGTACAAAAAATTAATGATGTTACGCGAAACATTAACGTTTGC
TTAATAGATTTCGATAACGAATCTTCTGGAAATACGGCGATAAAACTAACACTTTTATATAATCGCTCGATAGGAGTATTCGG
GCTGCCCTCAGATTTCACAGAAGAAAAACTGTACGATGAATTTTCAAGGTTAGTGTGTTAATGTGATAATATTTAACTATCTA
ATCTTATAATCATATCAAGTCTAACAATCTCAATTATATATTATAACAGTATGTCAACATTAGACATCTTTTTATATGCTTTC
AGGTATGGCAGAATAGAAAAAAATAAACTAGTGTACGACTCTACCGGACACTCTAAACAATACGGTTTTGTTTATTATGAAAA
GCACATCTCTGCTCAAGCTGCCAAAGAGGAAATGGACCGCAGTGATTATACAGGAGGTAAAATTTCCGTCCGTTTTGTTCCAG
AAAAAGAGTAGATCATCTTCATCAGTGTAAGGGTATGCCTAAGCAAAGCAACTGGAACTACAACTATAATGAAGATGGCGTAG
TTTTCTCAAATATTCTGAGATTACTCATTATCTATATATAATTAATTCGTCGGACCCGCAAATCTACTAATCGCGGTTTGATT
GGCCGATATTCTCTAAATTTACTAAGCAACTTGCACGCAATTAAGGCTGCTTAGTGCATCATCATCCTCGGCAGTAATTGCTC
AAAATTTCAAAGTGATAAAATTCAGCCCTCGAAGCAAATATTGGTGTGAAAAAGTATCATCCTATGTGCTCACTTGTAGTGAA
TGTGATGATACTTTTTCAGATGAGTAGTTGCAAAACTATTCGTTTCTTATTACTTCAAAAACGCGAGTTAAATAATCATTGAA
AAGCAAAACATCAATGATTCGAATGAAAGTTCAATGATGCACTATATCACCTTAAATTTACCGCAAGTTTCTCAACAAGCATT
TTTGTGCCAGCACAACGTTGGTTTGATCTGGTATGACAATAAAAATCAAATCTTT

>Ae.riversi
GGCGTATCTGCAATGATCGATTTATCTTCATCGACTTTCTCTTTGGTTAATAACTTGGAA
AGCAAATCGTTTTCGACTGTTTTTGTTGTTGAAGACGCTAGTCCTTGTCATCTTCTAACG
GGATACACCGAAAGATCACCCGAAAACAGATCGTGTTTTCCGGAATAATCCAAAAAGAAA
ATCGATGAAGAGAAATCGATCATTGCAGATACGCCCGTATGATACTAAACGCGAGAATAT
ATTTAGAAAATGGTAAACTGATCTTAGCGTGTAGGTCGTACACATTGAGCTCGGAATTTT
AAGTGTGCTCAACACACTGCTTTTTTTTCTTTTCATTTTTGGCACGAATTATTATTTTGC
GTAATGTACAATAAAAGTGAGATCAATCTCATTAATAAGCAATTTGAATGTATTAAAAAA
TATTGCATATATATTGGAAACATTCCTGCCGAAGTGTCGAAACGAGATTTAGTTGCTAAA
TTTTCCGAATTCGGTGAAATTTCCAACTTATATATGAAGTCATTCGTAAGGTTCTGTGAT
GTGAAACATGCTGTAATTCGTTACAGATTAAAAGGAAGTGTAAAGGAATCTTCAAGTTTA
CACAATAGTCGATATATTCAATCGGTTTTAATAGTTCTGCCACTAGGTTTGTCGTACACC
CATTACTATCTTCCTTACAACACTTGTGTTGCGGTGTACACTAACAAAAATTTTGATATG
GTAGAACTTTCTGAAAATTTAAGGAGATTTGGAGCCATTCAAGTCATAAAGAAAACTACA
AACGTCATGGGATACATTTGTTTTACATCAGAAAATGCTGCAAGAAACATGTTGGTTACT
AAGCCCACAGATATCCATAAAAATGTACAAAAAATTAATGATGTTACGCGAAACATTAAC
GTATGCTTAATAGATTTCGATAACGAATCTTCTGGAAATACGGCGATAAAATTAACACTT
TTATATAATCGCTCAATAGGAATATTCGGACTACCATCAGATTTTACAGAAGCAAAACTG
CACGATGAATTTTCAAGGTTAGTGTGTTAATGTGAACCATATCTTATCAATCTATTATCA
ATCTTATCAATCTATTATTATATTATATTAAAACGTATGTCAACATTAAATGTTGACATC
TTTTTATATGCTTTCAGGTATGGCAGAATAGAAAAAAATAAGCTAGTGTACGACTCAACC
GGACACTCTAGACAATACGGTTTTGTTTATTATGAAAAGCACTTGTCTGCTCAAGCTGCC
AAAGAGGAAATGGACCGCAGTGATTATACAGGAGGTAAAATTTCCGTCCGTTTTGTTCCA
GAAAAAGAGTAGACCATCTTCATCCGTGTAAAGTAACTGGAACTACAACTATAATGGAGA
TGGCGTAGGTTTCTCAAATATTCTGAGATCTACTCATTATTATCCATATACAATTAAATC
GTCGGACCCGCAAATCTACTAATCGCAGTTTGATTGGCCGATAATCTCTAAATTTACTAA
GCGACTTGCACGCAATCAAATTTACCGAAAGCTTCTCAACAAGCATTTTTGTGCCTGCAC
AATGTTGGTTTGATCTGGTATAACAATAATATTCAAATCTTTCCAACAAAAAAGCTAACC
ATCTGCAGAGCAATACCGTATTTCCTTGTGTAACAAGAGCCTTTTTATGCGATTTTGCGA
GCCTAGGGCTCTAGTAGTTATTAAAGATATTCAGCAATTGGCATTTCGTACTAACTAATT
ATATAAGTATGATAGAACCGTATGACGGTATTCTCCACATTGATTATTGTTTCATTTGGC
TAAAAGCATTTTATGTAAGTGATACTGAGATTAATGCGTGAGAGTATAATTTT

>Ae.vexans
TTTGTTTTTTTTTTGCCATCCACTAATACAGGTAGTCGTGAGAATCTCAGTTCAAGAAGATCCGCTCTGTTACGATGGAGCTG
AATATTTCAAGCGATCCGCTTTTGGATCTTTTGAATGCCCGTGCCGATAAGGAAGCT
```

CGTACCGAGGATAGGATGTTTACTCCTTTACATTATGCCGCAAGTCATCGCCACCTAACGAGTGAAGTAATCAACACAGAATT
TGAAATTATTGCTCAGAATTGTATTTACATTGGAAACATTCCAAATTTTGTTTCAAA
AAAAATATTGTTTGATTTGTTTACTCAATCTGGCGGTGAAATATTTAACATTTATATACAGCAAAATGAACACTGCGATGTAA
AAGCAGCAATCATTAGATTTCTACATAAAAAAAGTGTAATGCGATCCTTAAGTTTGA
ACAAAACTCGATATCATCAGTCAATACTAATAGTAATAGAACTAAGTTTGCCATATGCAAACTATCTACTAGTATATAATACA
TTGATAGTGATTTATATCCGAGAGAAAACAAAAAAATATTCACTGGAGCAAATTTAC
GATGAGTTTAAAAGTTTTGGAGCAATTCGAAATATTTTGAAAACAACGAATTTAATGGTTTATATTAATTATTACTCAAAAAA
AGCCCAAGAATCAGCCCTTAAATCAAAGTCACTTCCTGATTATGTTGATAAAATAAT
TACATCTCATAGAAATATACACATGTGTTGCATAAATTTAGACAGTAAGGATTTCTCGAGAAATTTAGAAATCAAAATTAAAC
TTTTATATAGCAGTACCATCGGCATATTTGGATTGCCATCTGATACCAAGGAAGCAG
AAATTACACAAATATGTTCAAGGTAAGTTTGTAGATTAACTATTACTAAAACTTGCATAATTTATAATTGGGCCGAAAAGTCA
TAAGTGACCATTTTACAATGTAGCTGATGTTACGTTTTGTTAAAAATAAACATTTAT
TAGTCA

nnnnnnnnnnnnnnnnnnnnnnn

CCTAGTAATTATCATTTTCAATTTCTGAAACTTTACTTATATTCGCCTATAAATCATTCTCTAATCTTTTCAGACTTTAACTA
TTTTTTTTTATTTTTTTTAGGTTTGGCGACATTGATAAAATTAAGCTTATCTATGACA
AAGGTGGAAACTCGAAGCAATACTGCTTCGTATATTACAAAAATCACATTTCTGCAATTGAGGCAAAACACAATCTGGACAAA
AAACCACTTCAAGGACGCGAAATTTCAGTGCGTTTTGTTCCAGAAAAGGAGTAAATG
GTTTGTTGAATAAGAATGAAATTTTAATGATATAAATCATAGCGAACTTCAAAATGAAACATTCATTTTTAGAGCATACGGAA
CCGTTCAAATGGGGTCATGCACAAATTAC

>Ae.atropalpus
ATTTTTGAGTGTCCTAACAGAACAACCATTTCGTGTTTGAATTTCGTCTTTCATATTAATTCATAGCAGAATTAAAAGGAAGA
AATTTGTATTAGAACACATAACAACAAACTAGGTTATATTGAACACGTTCAGACAAACCAATAAAAAGGAAAGTTACTGAACG
TAATTTTTTATCAGCAGGGTGTGCGTTACATTCATGTAACGCTTATCTTCGGCGAGCTGCTCTTCGGCGAGTTGCCTCTTTCC
CTGTAACGCATATGTTCGGCGAGCTGCGCATCGACGAGTTGCCTCTTTTCCTCAACCGGTTATTCAAAAAGTAAATTGTAGTG
GTTGGTTTTCTCTGTCGAAAAAATGTCAATGCTACGTGAAGAATTCTTGAAAATCAAGAAAAACGTTGTCTATGTAGGAAATA
TTCCCTTGAATACCACAAGAAACGATATTTTGAACCTTTTTAAGACATATGGTGAAGTATGTACCATAAGTGAGCATAAAACG
GCTTCAGTTAAAATTGCCTATTTGAGGTACGAGAAATATGAACACGTTAAAAGTTCGGTTCAGTCGCTTGATAACAAACAATA
TGGTCAAACTATTCTAATTGTAAAGCAGTTAGAAATGCCTTATTGGTATTATATACTACCGTATGAAACAACATGTATAGTAA
GTAATTTCAAGGAAAGTACAGAATTAGTTGAAATAAAAAACGTCTTTGGTGCATTTGGAAAAATATCACAGATTTTGAAGACT
ACGCATACATTTGTGTATGTTTGCTTTTACACAAAAGCAAGCGCACAGCTTGTACTGTCAACATATGTTAATGGATGGAAAGT
ATCTTCTACCAGACGCAATATAAACGTTATAAACGACCAATTTGATGTTGAATACCACACGCCAGTAACTATCAAAATTATAG
ATCATTTATTGCATCCAAGATGTTTAGGCATCTTTGGATTGAGTAAAGCAACAGACGAAAAAAGCATAGAAGGATTATTTAAG
AGGTAAGTTTTGTTTGCAACTAATTATAGCATGTTTAAAGAGCTTCTGTTTTTTTTAAATAGCTTGGATGGTAATATAAAACT
TAATAGAACAAGGTATTTCCAAACTAAATTCCAGCATTTCTTTAATAGTCCTATATACATAAGCCTGAGAATCAGACAGTTGC
AATTTAGATTTAATCTAAACCTGTTTATGTCTTCTGGTATTTAGTTCTAAATGGAACTGACATAGTTATGGTTATTTATCAGA
ATAAAATGTGATCGTGTCCATGTGAGCACTCCACAATAGATTCTTCACATAAAGGTACACA

nnnnnnnnnn

AAACTTAACACAACACGAGTAGAGTGTAATGACCGTAAGACCGTGATAGCAATAAATAAAGAACGACGGTACGTCTGTATCGT
TCCGTTATCAATTTTGTACTGTTTATTTACATTTTCATTTGATGATTGAAGAATTATTTAAAACTTCTTAGATTTGAAAGCAG
GTACTAGATGTTTTCATTATTTTATAATTAGAAATAAATAACCTGATAGTATAGGAATTGGTTTTTTGTAATATTTAAGTATG
ATTTAAAACAATGTTTCTTACTCAATAATATTATTTTATTTTCAGGTATGGAAATATAAAATCTATCAAGCTAGTGCTCAACA
GACAAAAAATATCAAAACAATACTGCTTCATTACCTTTGTGGATCCTCTATCAGCTCGGATGGCTCAAAAAGCTCTAGATGGG
ATTGTAATTGATGGACGTATCATATCTGTAAGATATACTCTGGAAAGATAATAAAATATATTTTTCAACTATTAAATAGAGAT
TTATATGTGAATGTCTAAAGTGTGTGCGCGATTCTGTGTGTGTGTGTTTTTTTCTTCTTTGAAAGGTCAAAATGATTACAGCC
TCTCGTGATGCTGTACGCTATGCCCGGGCTACTCTCATGCATATTATTCTATAAGCACTTTTGTAAAAATCACCCACTCAGCC
GATGTGCAGGCAGTGATTTTGACCTCACCTATAACGGAGAAGGACCAATGTAAAAATCACCGACTCAGCCGATGTGCAGGCAG
TGATTTTGACCTCACCTATAACGGAGAAGGACCAATGTAAAAATCACCGACTCAGCCGATGTGCAGGCAGTG

>Ae.japonicus
GACTCGAACCAAAAACAATAAATCCCGTGATATAGGTTGATTTTAATAAAGACTGTAGTTTTAAACCTGATATTCTTACTCTC
GAAGTGAATTTTACTATACAGAATGAGTCTGAAACTCAAAAATTCATTTGGTAAATTTTACCCATTTGTATTTAAAATATTAT

ATCCAAAACTAGGTATATTTTACCCATTGCTATTTCAAAAGCTATTACCTAAAATTAGGTAAATTTTACCCAATGTTAATTTG
A
ATGGCATTACCCAATATTAGGTATATATTACCCATTCGTATTTGAAATGTTAAAACCTAAAATTGAGTTTTCAGCTAATATCT
AATTTCACTGGAAATAACTTGCATGGCATTTGATATCAAGATACTCTATATTACTTAATATAGTTTGTAGTGCGAACGGGGTA
TTATCATGTTTCAACGCTGTAGTCTGAACAAAGAATTATATGGATTTTTCCTTACCCATTAATGGGGGTTTGCAGACGACCGT
G
CAAGACCAAAACAACCAAATAAAACAACAACAATCAACATCGGATCGGCAGCCATTTGTTCTGAAAACCGCTAGTTAAGTTCG
CACTGTCTTGAAGTTCGGTTTTGCAATTGGTTGGAACCAGTTCTAACCCCGGTAATTAAACGGAGTCCTTAGAGGACAAACGG
TTAGACGTGATTTTTTTTATTGTTTCTCAAAGTATGAGAAATTGCATGGAAAAATTCGATCAAATGAAAAATAAAATAGTTTA
T
ATTGGAAATATTCCCCTAGGAGCTTCGAAAAAAGATATCTTTATGTTATTGGAGGAATATGGTAAAATATTTACTATAAGTGA
GAATCAAAACCATGCAGTGAAAACAGCCTATGTGCGGTTTTGTGATCCAGAATCTGTAGATAAGTGTTTAGAAAAAAATAACA
CTACGTATTGTGAATCTATATTGATAGTAAAGAGAATGGCTATGCCATATTCATACTATTTATTACCCGTTGAAACAACGGTG
T
TAGTATCTACAGAACTACAGGAGAAAAACATAACGCTAAAAGACATTCATGGCATATTTAAGTGTTTTGGCGAGTTATTTTGC
ATTTTACAAAGAACAGATACGTTTGTATACGTATCCTTTTGCTCTGAGGAAAGTGCACAGATTTCTTTATCACGATCTATTAA
AATAGGTGGATGTCCAATTAAGGTTGGTCAAATCTATAGAAACATCAATTGCCGGCTATTCGACATCGAATATAAAATCAACC
A
AACGACTGAAGCAATTATTAAAGAAATGGTGTATCCCCGATGCTTAGGGATTTTTGGGCTCTCCAAAGAGACAACAGAAACGA
GTATTGCCAAACATTTTTCCAAGTAAGTAATTTGTTATTAGGTGAACTGAATTTTTAATTGAAAAAAATCGATCTTTTTTTCG
TCTACATCGATGACGAGTGTTCCTACTAGCACAATCTTTTTTTTTTAGATCTCAACTATTTTTTCTCAATTTTTAAACTAAAA
A
TTAATAAAAAAATTCTAACAAACCGGTATGCAAAGTGACATTTATTCAAAGTTTACAACAAGTTGACCTACACGCTCAAAACT
GCCTCTCTTTCATTTGTATTGCTGTCCCTTTTTTTCTCACGGAAAAGTTACTCAATTCTCGTAAAAAGTGGAACAACTCAA

nnnnnnnnnnnnnnnnn

CTGCCAATAATCAAATCCCCCGCTATAAGGCGACTCTGTATCGAACTTCAAGACTGCTAATCACTCGCGACAAAGCGGAAAAA
CATTCGAACGCCATTATTTGCGTCCGGATGTCAAAGTACTACCACTAATAAACAAAACTAAGCTGATGTAGCTCTTTTTCCGG
GTGCTGGGGCGAAGCACAATAAATCAAAAAAACGTGAACATACTACCGACACTCACACATTTCGGAAGGAAATCGATTGTATT
TCGTCCTTCCCAAAGCAAGATTTTCTGATTTCAATGTTAGATTAAGTTAAATTACGAGTTATTGTTGATCGTACTTGCCACAT
ATTGATATTAGTTCAAAAACTGTACCTGTTATCGTAATTAATTGCTGTGGAATTGCACACAAAGTTGAGATACACTATTCGAT
AAACTGATGATAAATGCAACATGCTTAAAATTGTAAACAAGAAAGAAGCATGAATAATAGGAGTCGTCTCACCTCTTCAACAG
CTATGAAGAAATGCTTAAATTATTATAATTCTAGAGAAATAATGTTTTACTAAGATTGGCCAATTGTTTCATTGCAG

GTATGCACCCATATACAAGTGTGTTTTGATAAGAACACCAGAAGGAAATTCAAGAAGATTCGGTTTTTTGTACTTCGACGATC
CACAATCCGCACAAGCTGCCCTTGCTGACCTTGATGAAAGTAATATTGATGGATGCAAAATTTCGGTACGTTTTGCTCCCGAA
AAATATGATTAATACCTGCTAAACTGCAAATAAATACAAACTGACATATTTGTTTAACGTTCGATAATAAAACTAAAGCTAGG
CAGGATTCAACGGCCATGTCAAACAATCCAAAATCAGAATCACGCATTTTCCAATTTCATGGAAATTGATATTTTTACACGA
TTAGTGTCTTCCAAAAGCTTCATCAAAATAAATTTGTTTACCTTTTATTAATATTTAGGGACAATGTATCCGCATAATAATGC
GAACCTAGCTTCGCAGTAGCAATACTTCAGTGACGTACTCACTTTTGAAATTTCGAGTGTACCGAAGATAATCTCTCGGCATT
CGAGCGTACTAGTAGAGCAGACCAAAAGCTCCATAAATTTTGTTTCATAGAACCTTTTATGGATCTATTCATCAATTTTAATG
AAATTAGTAACTTTTTCATCTAACTTCGTTTGGAATATGTGATATGGCGAAGGTCTCAGGAATACAAAAATGGACGCTATCAT
AGCTAATTCACATTGACAATTCGGCATGAGTGCCCCTGCAATCCTTATGAGCACTCATGCTGAATTTGGACGCTAACGGACCA
TTGTGGCTCCAGTTATAGCGTAAACCATGGCTGAGTAAACCTTAGCAATGTTGGTCTCTGCTTTTAAACAATTTGAGGCTGAT
TTAT

>Ae.triseriatus.1
GCACTGGAATGCTTAAAGTTGGTTTATGCATCCAATGCATAAACCCGATTTATTCATTCC
AATGAGTGGATAGACCGAATATGGTTTTAGCGTAAAAAATATGCATTATAATTTAATTTT
TCTTGTCCGTGCAGAATATGGTTTTGGCGTAAAAAATATGCATTATAGTTTATTTTTTTC
TGTCCGTGTAGCGTGTGTTTGGGGTTAATTTTACTCAAATTTGGATAAATTAGAATGAGC
GTGTAGTTTTGAGCTGTCATTGTACAGACTCATGATTAACATAAAGCCGGGCGCCCACTG
ATGTGGCGGAGGTGTGGCATCCGCTAGCATGTCGAAACATCATGCCGACGTTCGCCACAC
GCCACACGACTTTGACGCGATGTTTTGACATGCAAGCAGCTGTCACACAGCCGCCACATC
AGTGGGCTCCCGGCCTAAGAACCTTTAATTTAAGTGCAATAAATATGACAGATATCACCA
AAATTCCTGAAGAATTTCGTGAAATCAAAGAGAATATTGTTTACATTGGGAATATTCCCA

TTGATTCTTCGAAAGATGAGATAATCAATTTAATGAAGGAATACGGTGATATCTGGAATA
TATATTTACAAAGCAATGAAGAAGCAATGTGCAATGTGAAAATTGCCTATGTGCGGTTTA
TGAATAAGTGCGATGCTAAAAACTGCACTAAAAAATTAAACAATAAAGAATACCGGAATT
CGGTATTGATTCTTAATAATCTAAGAAGTCCGTACACGTATAACGTGTTAACGTACGAAA
GCACGGTTGTGATAAGCAATTTTAAAGATACGCACCTAGCAGCAATAAAAAACGCCTATG
AGGCTTTTGGTAAAATTCACACCATTTTAAAAACAACAAATATTTTTGTATTTGTTTCGT
ACTATAAAAAAAAATAGCGCAGAATTAGCGCTATCAAAACGCGTTACGAAAGATTGGAAAA
TTTCAAGTATACGAAGAAATATAAATATTCGTATAATTGATATTGAGCATGATAAAAATG
GGAAGACTGAAAGAATTATAAAAAATCTGTTGTATTCCCGATGTTTAGGAATCTTTAATT
TACATCAAAACACAACAGCTAAACGAATAATCGATTTTTTTGAAAAGTATAAAGATTAGT
CATTGTAAGTATAAATAATCTAATTTTTTTTATTGCTTTGCAGGTATGGAATAATTACAA
AAACGAAATTAATACTATCAAAGGAAGGAAACTCGTTGCAATATTGTTTCATATACTTCG
ATAATCCACTTTCCGCAGAAAAGGCCCGTAAAGCTCTCGATGACACTAATATTGATGGAC
GAGTCATATCCGTGCGGTCTACTGCAGAAAGAGAA**TAA**TCCAAGGATGTCCCTTCAATAG
GTCGTAAGCTGGGTCCTAGGGTACACACTCAGCTTTCATCTTCCAGCTCGGTTATTTTTT
TTACCGAGATCTCAACAGCAGGTACCCTCAGTAATCTAATTAGCTGTATTCCGGTAAGTT
ATTTCGACAGTTTCATTACCCGAGCAAAGTGTGAGTACAAAATGATAGCAGACTGAAAAC
AAATTTTGATATCATCATCAAACTGAATACTGAATTTGATATCGCTTTATGATCGAGACC
AGAGCTGGTAAATGTCATTTGGTAATTTCATCCTGTCATTGCGTCTAACTTTTTTAATAA
GCAATATTCCATAAAATGGCAAGCGAAGCAATTGTAGGTCTTGATCTAAGCTACAACTTT
TGCGAATCGACAAAATCTGTATCTCTGATACATAACGAACTAGCGCGATAGAACGGGTTA
ATGTTTATCAATGACGGTCATTGATAAACATTAACCTGTTTTATCG

>Ae.triseriatus.2
GCACTGGAATGCTTAAAGTTGGTTTATGCATCCAATGCATAAACCCGATTTATTCATTCC
AATGAGTGGATAGACCGAATATGGTTTTAGCGTAAAAAATATGCATTATAATTTAATTTT
TCTTGTCCGTGCAGAATATGGTTTTGGCGTAAAAAATATGCATTATAGTTTATTTTTTTC
TGTCCGTGTAGCGTGTGTTTGGGGTTAATTTTACTCAAATTTGGATAAATTAGAATGAGC
GTGTAGTTTTGAGCTGTCATTGTACAGACTCATGATTAACATAAAGCCGGGCGCCCACTG
ATGTGGCGGAGGTGTGGCATCCGCTAGCATGTCGAAACATCATGCCGACGTTCGCCACAC
GCCACACGACTTTGACGCGATGTTTTGACATGCAAGCAGCTGTCACACAGCCGCCACATC
AGTGGGCTCCCGGCCTAAGAACCTTTAATTTAAGTGCAATAAATATGACAGATATCACCA
AAATTCCTGAAGAATTTCGTGAAATCAAAGAGAATATTGTTTACATTGGGAATATTCCCA
TTGATTCTTCGAAAGATGAGATAATCAATTTAATGAAGGAATACGGTGATATCTGGAATA
TATATTTACAAAGCAATGAAGAAGCAATGTGCAATGTGAAAATTGCCTATGTGCGGTTTA
TGAATAAGTGCGATGCTAAAAACTGCACTAAAAAATTAAACAATAAAGAATACCGGAATT
CGGTATTGATTCTTAATAATCTAAGAAGTCCGTACACGTATAACGTGTTAACGTACGAAA
GTACTGTTGTGATAAGCAATTTTAAAAATACGCACCTAGCAGCAATAAAAAACGCCTATG
AGGCTTTTGGTAAAATTCATACCATTTTAAAAACTAATAATTTTGTGTTTGTTTCGTACT
ATAAACAAAATAGCACAGAACTGGCGCTATCAAAAGGCGTCACGAAAGATTGGAAAACTT
CAAACATACGAAGAAATATAAATATTCGTGTAATTGATATTGTGCATAATAAAAAAGGAG
CCGAAAAAATAATAATAAATTTGGTTAATTCAAGATGTTTAGGAGTTTTTAACTTACATC
AAAACGCAACAAAAAACCGAATAAACGATTGTTTTGAAAAGTAAAAAAAAAACATCAATG
TGAATATAAGATAATCTAATTTTTTTTATTGTTTTGCAGGTATGGGGAAATAACGAATAC
GAAATTAATAGTATCAAAGGAAGGATATTCGTTACAATATTGTTTCATATACTTCGACGA
TCCACTTTCCGCAGAAAAGGCCCGTAAAGCTCTCGATGAAACTTATATTGATGGAAGAGT
TATATCCGTACGGTGTACTGCAGAAAGAGAATAATCCAAGGATGTCCCTTCAATAGGTCG
TAAGCTGGGTCCTAGGGTACACACTCAGCTTTCATCTTCCAGCTCGGTTATTTTTTTTAC
CGAGATCTCAACAGCAGGTACCCTCAGTAATCTAATTAGCTGTATTCCGGTAAGTTATTT
CGACAGTTTCATTACCCGAGCAAAGTGTGAGTACAAAATGATAGCAGACTGAAAACAAAT
TTTGATATCATCATCAAACTGAATACTGAATTTGATATCGCTTTATGATCGAGACCAGAG
CTGGTAAATGTCATTTGGTAATTTCATCCTGTCATTGCGTCTAACTTTTTTAATAAGCAA
TATTCCATAAAATGGCAAGCGAAGCAATTGTAGGTCTTGATCTAAGCTACAACTTTTGCG
AATCGACAAAATCTGTATCTCTGATACATAACGAACTAGCGCGATAGAACGGGTTAATGT
TTATCAATGACGGTCATTGATAAACATTAACCTGTTTTATCG

>Ae.triseriatus.3

TAACATAAACATGATTAGTAATGAAACGAAGTTATTTTTTCATCCATTAATGGGTAAAAA
TATACCCATTATTGATCCCTGAACAACACAAACCAATGATTAGCTCCTCAACGTCGTTGG
GACAATTAAATGGCGGCAAGTTGAAAATGTGCCCATAATTTCTGAAACAGTCTATAAGCA
TGCATATTTGCATTTCTCCGTGTATGGGAATGACATTTGTGTCAAACGAGCCTAACACCA
ACACGTGAAACTAATATCAAAATTGAAGTATCAGCATTATTGTCGTTTGGTAAAATGGCT
CATAGTACAGATCCTATTTTTAAAAGATTTAGTGATATCGAAGATTACGTTGTTTACGTG
GGGAATATTCCCTTTGATGCTTCTAAACAAGACATAATTAGGTTGTTGGGGTATTACGGT
ACAATACGTAATTTGTATTTACAGAACACAAAAGAATCAGTGTGCAACGTGAAATTCGCG
TATGTGAGGTTTCTGCATAGTAGGCACGCTCGAAACTGCGCGAGAATTTTAAATAATAAA
CAATATCGACAGTCGATATTAATAGTTACACCAATGTCAGAACCTTTATCGTTTAACCTT
TTAATTTACGAAACAACTGTTGCAGTGAGCAATTTTGAAGATGCGTCCATAGCAAAAATA
AAACGCGTCTATGAGCGTTTTGGACCTGTTCACGTCATTTTAAAAAGAACAAACACTTTC
GTGTTAGTTTCGTATTGCGATCCAGAGAGTGCAGAAAAGGCACTAGCAAAAGAATTGTCG
GGTTTTAGAACAACAAGCATAATGGGCAATGTAAGTAGTCGCATAATTGATATTGAGCAA
GATGAAAATCAAAAGACTAAAAAAATAATTAAAATTCTGAGTGATCACCGATATTTGGGA
CAATATCGTTAGTCCGCGGACATGGGGATCAGGGGGCGGCGAGGGGGTGAAAATGACGGC
TGGCAAACTGGCGAGTGTTTATGGCCTTTTTCCTCCAAGGAGGTGTTAGAGAAGACGAAC
AAACAATCTGATTGTCGCGTTGCCGATTGTACAACTGGAAAGTTAAGGTAAGCAGCTTGC
GTGTTTAAACCATAATGTATTGATTAAAAAAAACTTTTCAGCCTTTCTAGATGCTGTCCT
TTGTATTTGCGAGTTCTGCCATAGTGCAGTGGTTGGTACCGCTGGGAGAGAAACGCA

Stop codon not determined
>Ae.triseriatus.4
ATCAAAGCAGCTCCAGAGATTCTAGATCGGGATTTAATCCGAGGTTAACACTAGATGGAG
CTTACCACCATTAGAGTAACTATAATATGCATAATAAGAGTGACGTCATGTGTCAGTTTT
GAAAGGTCTTCTGCTTGTTAAGCCATTTCATGTGACAGGTCCGGCTGTGCAACTGTTTAC
AATCTCCAAACAACATTTACATACTCCAGTCTGTCATTTCCATAGAAGAACTTACAAACA
GCTGGAAAATCAGCTGTTCTGAAACGTCTCCCGAAATGCCTTATCGTTACTCCACGTGCA
CTAATGTTCCTCGTGTGAGGCAAGGAAGAGAATGGCAATAAAATGTATGTTATCATATTT
AATATGATAACTACTAAAATTTCTAAAGGAATTCTTGATATAAAAAAAAGCATTGTTTAC
GTAGGGAATATTCCCATTGATTCTACGAAAGATGATATATTCAGTTTAATGAAGCAATAC
GGTGATATCTGTAATATATATATTTACAGAACAACGAAGAAGCAATATGCAATGTGAAAATT
GCCTATGTATGGTTTATGACTAAGTACGATGCTAAAAACTGTGAAAAAGCATTACATAAT
AAAGAATATCGGAATTCCATATTAATTATTAATAATCTAGGAATTCCGTACACGTATAAC
GTGTTAACGTACGAAAGTACTGTTGTGATAAGCAATTTTAAAAATACGCACCTAGCAGCA
ATAAAAAACG

>Tox.amboinensis
CTAATCTAGCGTTGCTCCTCAGGATCAGCAGATGGCAATTCGTGAATCCTTTGAAGAAATCAAATCATACA
TAGTCTATGTAGGAAATATTCCTGCGAATGCTTTTGAAAAAGATATCAAAAACATTTTTCGTGAGTGCGGAATGATCGAAAAG
ATTTGTATTCTGCGTTCGCGCCAGCATTGTCAACACAAA
ATAGCATACGTTCGCTTCAGGCATGCGCGTGGCGCGCGTAGGGCAGCAGGATTACATGGATCTTACATGGACGATGACAGTTA
TCTTATTGTCAAGTTACTCGGATTACCATATTCATCGCA
TGTTTTATTGCATGATTGCACTGTTGCGGTATCTTACTGCGGTGGAGATAAGCTTAAATATAAAACCAAAGGTACAGGTAAAG
GCTATACTATACAGGGTGATTGGTAAATACATTATAACC
TTAAAATGGCAGATTGAAAAAGATGATAAACGAAAAGTTTTTCAAGTAACAAAGCTATACCACGGCGATGATATAGTGGGATA
AAGTTTTTCCATAAGCGTCGCTATAGCTTTTGTCGTAGA
CCACCTACAGACCTTTTTCAAATATTCTTAACAAATTCTTGAAAAGTGTGTTGAATCTAAAATTTTTGACTATATTTTTTGTT
TGCTCAATGGGCAGGTGAATGGACAATAGTGTGATGCGT
TAGCTCATACCCTAAAGATACAGTGGCTCTCAAAATTGATCGT

nnnnnnnnnnnnnnnnnnnn

AATAGCATGAAATGAAATTAAATGCAATTTAAATGTGCATTACTGAAGTGAGATACCCCTAGGTTGGGCATCAATGAGTTAAA
TACTGATAAGTGAGCATGTTATACTCACATGTTTCATATTTATTTCTCTCTTATACCTTTAACGTTCAATATTCTGGAAATCT
TCAAGTGACATTTTGAATCTAGGTATATGTAAATGAAGCCGATTACCCTTAATGTATTTATCAACCTGTGTATTTATCTATTT
TCATGTTGTCTAATACATGTTTTTGATTTTATTTAGGTATAACTCACCTTGAATTATATAAACACTTCAGCGATTACGGTCGT

ATAGAAACAATTTTGTGCACACATCCAACATATACATATGTGAGTTTTAGCCAACCAAGCCAAGCAAGGCGTGCGCTGCTAAA
GACTGAGAGTACCATAAACCATCAAAAAATTAGAGTTTTCTCGATTGCACGCAACGTCCTGATACCTGGTTCACGAAATCTGG
CAATAAAAACCATAGACGACTTAATGAAAATTGATGTGTGCATAGGCTTTTTTGGATTAAGTCCTCAGTGCACTGAAAACGAC
CTGCGAGATATCTTGCAAACCTACAAACTACCAACGAATCTTATTTTAATTCGAGACGCGGTAACTAGTATGTCGCGACAGTA
TGCTTTTGTAAAATTTTCAAATAGCACAATAGCAATGGCGAAAACAATAGCACACACACTGAATGGAATTGAGATAAAAGGAC
GAAAAGTTCGTGTGGACTTAGTAGTAGACAACCGTGAATAAATATAAAACTTTGTTTGGTATTTGTTTGTAAATCTTCTGGTT
TACGGAAC

>Tox.sp.Nix
CAAGTTCAAAAAGTGCTAGTGGATTGTGTGTGATAAAGATAGTTATAATTTGTCTAGGTTGTAGATTCCAAATAGCAATTTTA
CAGTCACAATATTTGTTTAACCGATATTCGTAAAGCTAATAAGTTGGTAGCCGAGAAATCAGAATGCAAACACTTTAACTTAT
ATCACTAAGATAAATCCGAGAACAATTTTCAAATACAAAAGAAATAGTTTGATACACTCCACGCCGAAAAAATAAGCTTTCAG
CTAATACTGCGAGACTACTCTGGATATGCAGATGGCAATTCGTGAATCCTTTGAAGAAATCAAGTCACACATCGTCTACGTAG
GAAACGTTCCGGCAGAATGCTCTGAAGAAGATATCAAAAATATCTTTCGCGAGTGTGGAAAAGTCGAGAAGATTGTTTTCGCG
TGCGCGCGAGATAAATGTGCCTTCAAGGTAGCTTACGTGAGATTTGATCTTCCACGATCTTCTCGTAAAGCAGCCAAATTACA
TGGATCTATTGTCAAGCTACTCGGACTACCGTATTCAATTATTGTCAAGTTACTCGGACTACCGTATTCATCGTATGTTTTAA
TGCATGATTGCACTGTTGCGGTATTTTACTGCGGTAAAGGTACAGGTATCACACACCTTGAATTATATAAATATTTCAGCGAG
TATGGTCATATAGAAACAATTTGGTGCAGTACGTACAACATTTTGCAGTACGTGCAATTTAGCAACCCACAACAAGCGCAGCG
CGCGTTATTGATGACGGAAACTATGATCAACCATCGAAAAATTAGAGTTTTTTCAATAAATAAGAATTTAAATTTACCTGGTA
CACGAAATCTGGCCGTAAAAACCATAAATGACTTAATGAAAATTGACGTATGCATAGGCTTTTTTGGATTAAGTGCTCAAACA
ACAGAAACCGACCTGCGAAATCTACTGCAACCCTATAAAATATTAGCTGTTAAGTTGATTCGTAATTCGTATGCTTTTGTGAA
ATTTGAATTTGCAAATGCAAATTTTGCAAAATTAGCACAAGAAAAACTTAATGGAGTTGAGATAAATGGACGGGTAGTTCGAG
TGGAACTTGTAGCAGGCAACCGCTGGTACACGAAATCTGGCCGTAAAAACCATAAATGACTGAATGAAAATTGACATATGCAT
AGGCTTTTTTGGAGTAAGTGCTCAAACAACAGAAACCGACCTGCGAAATCTACTGCAACCCTATAAAATATTAGCTGTTAAGT
TGATTCGTAATTCGTATGCTTTTGTGAAATTTGAATTTGCAAATGCAAATTTTGCAAAATTAGCACAAGAAAAACTTAATGGA
GTTGAGATAAATGGACGGGTAGTTCGAGTGGAACTTGTAGCAGGCAACCGCTGGTACACGAAATCTGGCCGTA

>Ae.detritrus.Nix
GCCCAATCCTAATCTGTGGGGCAGAAATATGACAGAGGATATTTCTATAAAATTTAATAAAATAAAAGGAAATGTACTGTACG
TTGGGAATATTCCCATTAGTGCGTCAAAAAAAGAACTTACGTGTTTATTTGGAACGTACGGTGAAATCTGTACATTATACTTA
CTGACCAGCAGTGAATTTGTGTGCGACGTTAAAAGCGCTTACGTACGTTTTGAAAAAAGTGAAGACGCCCAAATTTGCGAAAC
TAAATTAAACAACGCACGATACCGTCAGTCTGTATTGATCGTAAGAATAATGGCCAAGCCATATTCTTACTACGTGTTGCCAT
ACGAAACAACTGTCGTGATCAGCAATTTTGAAGAAGATACTGATTTAGTGAAAATAAAAAGCGCGTATGAGACTTTTGGAAAG
ATAGAATCAGTTTTAAAGACATCTAACACCTTTGTGTTCCTTGCATTTTATCATAAATCAAGCGCCCAACTGGCACTAACAAA
ACCTTTGCCATATTGGAAGACTTCTTCAATAAAAAGAAATATAAATGTACATCAAACCATCGATATTCAGCAAGAAGAAACGA
AAGAAGAACGATATAAAATTATGAAAATTTTACTTGGATATCAGCGCTGTGTAGGGGTTTTTGGATTGTCACAACAGACAACC
CAAACATCAATAGCGGAAAAAAATGGATAAGTACGGTAAAATCACCAACATTAAAATACCAGTGACGGCGCAAGGTATTTCGAA
ACAATACTGCTTCGTGTACTTTGACGATCCACGATCCGCACAAGCAGCCCGTCTTGGACTCGATAAGACTATTATTGATGGAC
GTCAAATATCGGTACGAGCCATATCGGAAAAAGAATAAAATAAAGAATACTGAAG

>Wy.michelli.Nix
AAATCGGACATCACTAGTTGTCTAATTTATTGTTTACCATTTTATTTAAAATTTCTTTGCGTCAGTAGTTAATATTTATAGTT
CAACAAGTGAAAATTTTCAATTACTCATATTTATCAATAAGTAATAAGGAATTTTTATTTCCACAGTAAAAAGCGGTGCTTAT
ATTTTGTTTCAATTGTGAATATTCACAGAAATAGCTGCTAAAATACCTGATGTTTATAATATTTGCAAATAATCTACAAGTAG
TAAAACAATCAGTATGTCCGGTGTAAACCGAATCTACCAGCAGCATTTAGTTGGAAAAAGGATTGCGTTAAATGGTATCTTCT
GAGACGTCAAGTATACTACTGAATCATACCGGTCAGAACATGGAAATGTGTAAAATTGACGAAAAATTCGATGCTGTTAAAGT
TTTAATTGTTTACATAGGAAATCTTCCAAGTTACGTGCAAGAACAACATATATATAGGCTGCTTAAATCGGAAAAAATTTTCG
AAACAGACGAAGCGATAAAAATTATAAGTATTCCGGAACGGCGTTGTGCTTTTCTTAGATTCTGTTCTTTAGTTTCGGTTAAT
AATACTCGTAGATTGAATGGATTGAAGTACAATGAATCAATATTAATTGTTGTTCCTCTTTCATCAAGCTATCATGAAAGAGT
ATTACTGGCCGAATGTAGCATTTCTATTCTCAATGTATATTCACAAATTACAATGCAACAGCTTTTTGATAGTCTAAGTATTT
ATGGTGATATTAAAGCCGCTGTGAAATTGACTGACACAAGTTTCATTTTTTCTTTTCAAGAACCTTATCAAGCTGAGGAGACT
CTGAAAAAATCTGGTGATATAATTCGAAAAATAAAATTTCTCGTAAAATCAAATGATGTAAACAACATGATGCTGGGTGTGAA
ACTGTTTAATCTTTTTAGGAATATAAATATATGTGCATACGATATTCAATTTACTAAAGGGAATAGTAAAGAAATGTTAAGAC
AAAAAACAAAAATTAAGGCTTTTTTATGTGAAGATTTTGTAGCAGGCTTAAAGCCAAATAAATATAATTGTTTAGGTATTTTT
GGTCTACCCTCTAAAACCACCGAACAAATACTACGTGAAAAATTCTGTTGCATTGACGAGAAAATCAAAATTGTGATGATACC
AGATACAAACTATTGTTTTCTATATTTTGATACAGTTTTTAAAGCAGTGATGCTCAAATATGAGCCTAAAAATGGATTTGTTG
ATAAAGAAAACATTAGAGAAAGAAACTTATTTTAGGAGAAG

Stop codon not determined

>Ps.columbiae.Nix
CGTGTACTGAACAACTGACAAAATTGAATGAATAATATTCAAATTTTGAATACTTTTTGACGAGCGTGTAAAGTCCATCATTC
AAATAGACCATTTTACGTGACAGCTAAAATCAGCTGATCGAAAAATTTATGAATGAAATTTTGACGAGAGAGAGAGGACTTGC
ATACAAAAAGTAAACATTAGTCTTCTCCGTTCTGTCATTTGACGCCCCCCGTAAAATGGCTTATTGTTTTTTTACTGGCGTAG
CAAGCTCAATTTGTTGCGTTTGTTTATGTTTCAACTAACTGTATGTATATTCTTGTTATACTCAGAAGACTAAAATATGGCAT
CTTATCAAAAAACAAGAGATAGCGAGTTTGATGAAATATCATCAAGAATTATATATGTAGGAAATATTCCAAAATCAGTGAAT
GAAAGACAAATACGTCTACTATTTCGTGACTGCGGTGAAATTATTCGAATTGCTTTTACTTGGTGCTATGAGTTTTGTTCAAC
AAAAGTTTGTTTTATAAAGTTCAAACTTCCTCGGAGTACAATTAAAGCACTAAAGTTTAATAGAAGTAATTTCGAAGATTCGT
TCATAATAGTGCTTCCTGTAAATGATAATAACGATTATTTATTAACCTACGAGACAACTGTGGTTGTTTCTAATATTCATCCC
GGAATGACCTTGTGGGCCATTTATCTATTGTTTAAACCGTTTGGTGTAAAAACAGTTCTCAAAACCACCGATACTTTTGTTTA
TCTGTCTTTAGAAACCGAGCAAAAAACTAAAGAATTATTACAATGTACTACTTCATGGACAGCTAATACGGTCAAAGTATGTT
CTATAAGAAGGAATGTTCGTTTAATTTTGAAAGACATTTTCAGTGATGATGAAGAAATGCGGGAAATAAGAAGTTATTTGAAA
GACGAAGATGTTTCTTGTACAACTACAACACGCGTTTTAGGCTGTTTTGGATTGGATCCAAAAACTTCGGAAGACAAAATAAT
GCAGATATTTTCAAGGTATGTATACTGGATTGCGTCGCAATTCGCACTTATGAGTGCGATAAAAAAAGTGCGACTTTGCGAAT
AAAACTGCGCTCCTTTATCGGATCGATTTGGTGTCTTCCGCGCACCTGTTTCTTGGTCAATTTGCGCACTTCTGTAGAAGAAC
CGAACTTGATCCGATGAAGGAGCGCAGTTTTATCCACAAAATCGCACTTTTGAGGGGGTCCAGTTAGCGCAGAAATGCATAAT
ATTTATGGTTAATTAAATATTCACCCATATTCTGATATTCGTTCGACCATATGCTTAAACATGGTTCGAACGAATATGAGAAT
AGTCCATTTTATGGAAGGCGTCAAATGACAGGAGCATAAAGAGTAATGTTTACCTTTTGTATGGAATTTTGACAGGAGCTTAA
AGAGCTCCCATTCATAAAATTATCGA
nnnnn
ACGAAGAAACACCTCGATCGCGCACCTAGAGTTCTTTCGAGCCCCTTGTCATAGACCCAAAATAAATCGTTTTCAAGTTATGT
ATGGATTAAACGCGTGTTTTTGGCATAATCCACCACTGTACGATGCACAAAAAAGAAATCGAAATAAGTGAACATGAACATGT
CTTTATGGTGTCCCGCATCTATAACTTCCTCTATCCTATATATGATTTGGAAACCATCAAACTAATAATTTTTCTCTGATTTT
GCTTTTCAGATACGGCCATATAGAGAATGTTCGATTGATCCGTAACAGAAAAAATTCAGTTTCTATGGGATATTGCTTTATAT
TCTTTGATAATAGTGATTCAGCTAGCAATGCCTATTCAGATTTGAAAGAATTGGTAATTGATGATCGAAAAGTAAGAGTAG

**Nix cDNA sequences derived from either trinity assembly of RNA-seq data or RT-PCR**

>Ae.atropalpusTRINITY_DN23179_c0_g1_i2_rev Trinity of RNAseq of mixed eggs
CAAAAAGTAAATTGTAGTGGTTGGTTTTCTCTGTCGAAAAAATGTCAATGCTACGTGAAGAATTCTTGAAAATCAAGAAAAAC
GTTGTCTATGTAGGAAATATTCCCTTGAATACCACAAGA
AACGATATTTTGAACCTTTTTAAGACATATGGTGAAGTATGTACCATAAGTGAGCATAAAACG
GCTTCAGTTAAAATTGCCTATTTGAGGTACGAGAAATATGAACACGTTAAAAGTTCGGTTCAGTCGCTTGATAACAAACAATA
TGGTCAAACTATTCTAATTGTAAAGCAGTTAGAAATGCC
TTATTGGTATTATATACTACCGTATGAAACAACATGTATA
GTAAGTAATTTCAAGGAAAGTAC
AGAATTAGTTGAAATAAAAAACGTCTTTGGTGCATTTGGAAAAATATCACAGATTTTGAAGACTACGCATACATTTGTGTATG
TTTGCTTTTACACAAAAGCAAGCGCACAGCTTGTACTGT
CAACATATGTTAATGGATGGAAAGTATCTTCTACCAGACGCAATATAAACGTTATAAACGACC
AATTTGATGTTGAATACCACACGCCAGTAACTATCAAAATTATAGATCATTTATTGCATCCAAGATGTTTAGGCATCTTTGGA
TTGAGTAAAGCAACAGACGAAAAAAGCATAGAAGGATTA
TTTAAGAGGTATGGAAATATAAAATCTATCAAGCTAGTGCTCAACAGACAAAAAATATCAAAA
CAATACTGCTTCATTACCTTTGTGGATCCTCTATCAGCTCGGATGGCTCAAAAAGCTCTAGATGGGATTGTAATTGATGGACG
TATCATATCTGTAAGATATACTCTGGAAAGATAATAAAA
TATATTTTTCAACTATTAAAAAAAC

>Ae.atropalpus-Nix-F1_R1 RT-PCR sequence
AGTTCGGTTCAGTCGCTTGATAACAAACAATATGGTCAAACTATTCTAATTGTAAAGCAGTTAGAAATGCCTTATTGGTATTA
TATACT
ACCGTATGAAACAACATGTATAGTAAGTAATTTCAAGGAAAGTACAGAATTAGTTGAAATAAAAAACGTCTTTGGTGCATTTG
GAAAAATATCACAGATTTTGAAGACTACGCATACATTTGTGTATGTTTGCTTTTACACAAAAGCAAGCGCACAG

>Ae.atropalpus-Nix-F2_R2 RT-PCR sequence

TTACACAAAAGCAAGCGCACAGCTTGTACTGTCAACATATGTTAATGGATGGAAAGTATCTTCTACCAGACGCAATATAAACG
TTATAAACGACCAATTTGATGTTGAATACCACACGCCAGTAACTATCAAAATTATAGATCATTTATTGCATCCAAGATGTTTA
GGCATCTTTGGAT
TGAGTAAAGCAACAGACGAAAAAAGCATAGAAGGATTATTTAAGAGGTATGGAAATATAA
AATCTATCAAGCTAGTGCTCAACAGACAAAAAATATCAAAACAATACTGC
TTCATTACCTTTGTGGATCCTCTATCAGCTCGGATGGCTCAAAAAG

>Ae.atropalpus-Nix-F3/R3 RT-PCR sequence
AAGTTCGGTTCAGTCGCTTGATAACAAACAATATGGTCAAACTATTCTAATTGTAAAGCAGTTAGAAATGCCTTATTGGTATT
ATATACTACCGTATGAAACAACATGTATAATTTTGAAGACTACGCATACATTTGTGTATGTTTGCTTTTACACAAAAGCAAGC
GCACAGCTTGTACTGTCAACATATGTTAATGGATGGAAAGTATCTTCTACCAGACGCAATATAAACGTTATAAACGACCAATT
TGATGTTGAATACCACACGCCAGTAACTATCAAAATTATAGATCATTTATTGCATCCAAGATGTTTAGGCATCTTTGGATTGA
GTAAAGCAACAGACGAAAAAAGCATAGAAGGATTATTTAAGAGGTATGGAAATATAAAATCTATCAAGCTAGTGCTCAACAGA
CAAAAAATATCAAAACAATACTGCTTCATTACCTTTGTGGATCCTCTATCAGCTCGGATGGC

>Tox_ambTRINITY_DN8221_c0_g1_i7
AAGTAGAATGTAGATCCGTGTATGGGCTGCTTAAGAAATGTCGCATCTCTATTATAAACAAATCAACTCAAAAAGTACCAGTG
GAATGTTGTAGTGTAGTGAAGCTGAAACGTTTGTTCAAT
CGATATTTGTAAAGCTGATAAATTGGTAGTCGAAAAATCCGAATGCAAACACTTCAGCTTATATCAC
TAAGCTAAATCGGTGACAATTTACAAAGACGTTAGAGATAACACACAAGCTGAAGGATCTTTCAGCTAATCTAGCGTTGCTCC
TCAGGATCAGCAGATGGCAATTCGTGAATCCTTTGAAGA
AATCAAATCATACATAGTCTATGTAGGAAATATTCCTGCGAATGCTTTTGAAAAAGATATCAAAAAC
ATTTTTCGTGAGTGCGGAATGATCGAAAAGATTTGTATTCTGCGTTCGCGCCAGCATTGTCAACACAAAATAGCATACGTTCG
CTTCAGGCATGCGCGTGGCGCGCGTAGGGCAGCAGGATT
ACATGGATCTTACATGGACGATGACAGTTATCTTATTGTCAAGTTACTCGGATTACCATATTCATCG
CATGTTTTATTGCATGATTGCACTGTTGCGGTATCTTACTGCGGTGGAGATAAGCTTAAATATAAAACCAAAGGTATAACTCA
CCTTGAATTATATAAACACTTCAGCGATTACGGTCGTAT
AGAAACAATTTTGTGCACACATCCAACATATACATATGTGAGTTTTAGCCAACCAAGCCAAGCAAGG
CGTGCGCTGCTAAAGACTGAGAGTACCATAAACCATCAAAAAATTAGAGTTTTCTCGATTGCACGCAACGTCCTGATACCTGG
TTCACGAAATCTGGCAATAAAAACCATAGACGACTTAAT
GAAAATTGATGTGTGCATAGGCTTTTTTTGGATTAAGTCCTCAGTGCACTGAAAACGACCTGCGAGA
TATCTTGCAAACCTACAAACTACCAACGAATCTTATTTTAATTCGAGACGCGGTAACTAGTATGTCGCGACAGTATGCTTTTG
TAAAATTTTCAAATAGCACAATAGCAATGGCGAAAACAA
TAGCACACACACTGAATGGAATTGAGATAAAAGGACGAAAAGTTCGTGTGGACTTAGTAGTAGACAA
CCGTGAATAAATATAAAACTTTGTTTGGTATTTGTTTGTAAATCTTCTGGTTTACGGAAC

**Nix protein sequences:**
>Ae.aeg.Nix_AAEL022912_288aa
MCKKRNAEINAEFDIIKKYCIYIGNIPFFASKNDVVVKFAEYGETCNIYMQSNKPHCDVK
PAIVRYRSRKSVDKSLCLNNSKFGNTILIVLPLSLPYSRYLLTYDTCIVVYINDKNSCKT
SMAELYDEFQKIGDIQNMFKTTNNMIYINFESEKSMQLSLATKPFLINNNIFKIKKVERN
INMCGLNSESQDFSTNLKKKLLYNRSIGIFGLPSNATEERIAQIFSRFGDIQKITLICDV
VGNSKQYGFIYYKKRTSANAAKVIMDGENFEGNKISVRFVPEKKVFKN
>Ae.alb_Nix.Full_282aa
MYSKSELNLINNQFEYIKKYCIYIGNIPAEVSKTDLIAKFSVFGEISNLYMKSFIQFCDV
KPAVVRYRLMKSVKESSSLHNSRYIQSVLIVLPLDSSYNNYFLPYNTCVVVYTYNKFGMV
DFYQKFSKLGDIHAMKKATNVMVYISFVSERAARTILDTKPTDIHINVQTINHVTRNINV
CLIDFEKECTSNTAIKLTLLYNRSIGIFGLPSNFTEAKLHDEFSRYGRIEKNRLVYDSTG
HSKQYGFVYYEKHLSAQAAKQEMDRSDHTGRKIAVRFVPEKE
>Ae.alb.Nix_duplicate_205aa
ILVEYSVFCEISNVYMKSFIQFCDVKSSVGRYRLIKYLKxSSSLHNSQDIQSILIVLQLY
TYHKLGMLDFHIKxSVNLGDIHVTNVTVYISFVSERAARVILDTNPTDTNNYVSRNIYVC
LIVCSIEIFGLPSDFTESKLHDEFxRRYGSIEKSRLVYDSTGNSKxTIWFCLLxNxLFTR
AEKQEMDRSYYTERKIAVRFVTEKE
>Ae.atr.Nix1_271aa
MSMLREEFLKIKKNVVYVGNIPLNTTRNDILNLFKTYGEVCTISEHKTASVKIAYLRYEK
YEHVKSSVQSLDNKQYGQTILIVKQLEMPYWYYILPYETTCIVSNFKESTELVEIKNVFG
AFGKISQILKTTHTFVYVCFYTKASAQLVLSTYVNGWKVSSTRRNINVINDQFDVEYHTP

```
VTIKIIDHLLHPRCLGIFGLSKATDEKSIEGLFKRYGNIKSIKLVLNRQKISKQYCFITF
VDPLSARMAQKALDGIVIDGRIISVRYTLER
>Ae.det.Nix_275aa
MTEDISIKFNKIKGNVLYVGNIPISASKKELTCLFGTYGEICTLYLLTSSEFVCDVKSAY
VRFEKSEDAQICETKLNNARYRQSVLIVRIMAKPYSYYVLPYETTVVISNFEEDTDLVKI
KSAYETFGKIESVLKTSNTFVFLAFYHKSSAQLALTKPLPYWKTSSIKRNINVHQTIDIQ
QEETKEERYKIMKILLGYQRCVGVFGLSQQTTQTSIAEKMDKYGKITNIKIPVTAQGISK
QYCFVYFDDPRSAQAARLGLDKTIIDGRQISVRAISEKE
>Ae.jpn.Nix_273aa
MEKFDQMKNKIVYIGNIPLGASKKDIFMLLEEYGKIFTISENQNHAVKTAYVRFCDPESV
DKCLEKNNTTYCESILIVKRMAMPYSYYLLPVETTVLVSTELQEKNITLKDIHGIFKCFG
ELFCILQRTDTFVYVSFCSEESAQISLSRSIKIGGCPIKVGQIYRNINCRLFDIEYKINQ
TTEAIIKEMVYPRCLGIFGLSKETTETSIAKHFSKYAPIYKCVLIRTPEGNSRRFGFLYF
DDPQSAQAALADLDESNIDGCKISVRFAPEKYD
>Ae.mas.Nix_288aa
MCKKRNAEINAEFDIIKKYCIYIGNIPFFASKNDVVLKFAEYGETCNIYMQSNKPHCDVK
PAIVRYRSRRSVDKSLCLNNSKFGNTILMVLPLSLPYSRYLLTYDTCIVVYINDKNSCKT
SMAELYDEFQKIGDIQNMFKTTNNMIYINFESEKSMQLSLATKPFLINNNIFKINKVERN
INMCGLNLESQDFSTNLKKKLLYNRSIGIFGLPSNATEERIAQIFSRFGDIQKITLICDI
VGNSKQYGFIYYKKRTSANAAKVIMDGENFEGNIISVRFVPEKKVFKN
>Ae.pol.Nix_282aa
MYSKNEINLINKQFEYIKKYCIYIGNIPAEVSKRDLVAKFSEFGEISNLYLKSFVKFCDV
KHAVIRYRLETSVKESSSLHNNRYIQSVLIVLPLDSSYTHYYLPYNTCVAVYTNKHFHMV
ELFENFKRFGDIQVIKKTTNVMGYICFTTENAARKMLVTKPTDIHKNVQKINDVTRNINV
CLIDFDNESSGNTAIKLTLLYNRSIGVFGLPSDFTEEKLYDEFSRYGRIEKNKLVYDSTG
HSKQYGFVYYEKHISAQAAKEEMDRSDYTGGKISVRFVPEKE
>Ae.riv.Nix_282aa
MYNKSEINLINKQFECIKKYCIYIGNIPAEVSKRDLVAKFSEFGEISNLYMKSFVRFCDV
KHAVIRYRLKGSVKESSSLHNSRYIQSVLIVLPLGLSYTHYYLPYNTCVAVYTNKNFDMV
ELSENLRRFGAIQVIKKTTNVMGYICFTSENAARNMLVTKPTDIHKNVQKINDVTRNINV
CLIDFDNESSGNTAIKLTLLYNRSIGIFGLPSDFTEAKLHDEFSRYGRIEKNKLVYDSTG
HSRQYGFVYYEKHLSAQAAKEEMDRSDYTGGKISVRFVPEKE
>Ae.tri.Nix1_278aa
MTDITKIPEEFREIKENIVYIGNIPIDSSKDEIINLMKEYGDIWNIYLQSNEEAMCNVKI
AYVRFMNKCDAKNCTKKLNNKEYRNSVLILNNLRSPYTYNVLTYESTVVISNFKDTHLAA
IKNAYEAFGKIHTILKTTNIFVFVSYYKKNSAELALSKRVTKDWKISSIRRNINIRIIDI
EHDKNGKTERIIKNLLYSRCLGIFNLHQNTTAKRIIDFFEKYGIITKTKLILSKEGNSLQ
YCFIYFDNPLSAEKARKALDDTNIDGRVISVRSTAERE
>Ae.tri.Nix2_276aa
MTDITKIPEEFREIKENIVYIGNIPIDSSKDEIINLMKEYGDIWNIYLQSNEEAMCNVKI
AYVRFMNKCDAKNCTKKLNNKEYRNSVLILNNLRSPYTYNVLTYESTVVISNFKNTHLAA
IKNAYEAFGKIHTILKTNNFVFVSYYKQNSTELALSKGVTKDWKTSNIRRNINIRVIDIV
HNKKGAEKIIINLVNSRCLGVFNLHQNATKNRINDCFEKYGEITNTKLIVSKEGYSLQYC
FIYFDDPLSAEKARKALDETYIDGRVISVRCTAERE
>Ae.tri.Nix3_204aa
MAHSTDPIFKRFSDIEDYVVYVGNIPFDASKQDIIRLLGYYGTIRNLYLQNTKESVCNVK
FAYVRFLHSRHARNCARILNNKQYRQSILIVTPMSEPLSFNLLIYETTVAVSNFEDASIA
KIKRVYERFGPVHVILKRTNTFVLVSYCDPESAEKALAKELSGFRTTSIMGNVSSRIIDI
EQDENQKTKKIIKILSDHRYLGQY
>Ae.vex.Nix_293aa
MFTPLHYAASHRHLTSEVINTEFEIIAQNCIYIGNIPNFVSKKILFDLFTQSGGEIFNIY
IQQNEHCDVKAAIIRFLHKKSVMRSLSLNKTRYHQSILIVIELSLPYANYLLVYNTLIVI
YIREKTKKYSLEQIYDEFKSFGAIRNILKTTNLMVYINYYSKKAQESALKSKSLPDYVDK
IITSHRNIHMCCINLDSKDFSRNLEIKIKLLYSSTIGIFGLPSDTKEAEITQICSRFGDI
DKIKLIYDKGGNSKQYCFVYYKNHISAIEAKHNLDKKPLQGREISVRFVPEKE
>Ps.col.Nix_277aa
MASYQKTRDSEFDEISSRIIYVGNIPKSVNERQIRLLFRDCGEIIRIAFTWCYEFCSTKV
CFIKFKLPRSTIKALKFNRSNFEDSFIIVLPVNDNNDYLLTYETTVVVSNIHPGMTLWAI
YLLFKPFGVKTVLKTTDTFVYLSLETEQKTKELLQCTTSWTANTVKVCSIRRNVRLILKD
IFSDDEEMREIRSYLKDEDVSCTTTTRVLGCFGLDPKTSEDKIMQIFSRYGHIENVRLIR
NRKNSVSMGYCFIFFDNSDSASNAYSDLKELVIDDRK
>Tox.sp.Nix_287aa
MQMAIRESFEEIKSHIVYVGNVPAECSEEDIKNIFRECGKVEKIVFACARDKCAFKVAYV
```

```
RFDLPRSSRKAAKLHGSIVKLLGLPYSIIVKLLGLPYSSYVLMHDCTVAVFYCGKGTGIT
HLELYKYFSEYGHIETIWCSTYNILQYVQFSNPQQAQRALLMTETMINHRKIRVFSINKN
LNLPGTRNLAVKTINDLMKIDVCIGFFGLSAQTTETDLRNLLQPYKILAVKLIRNSYAFV
KFEFANANFAKLAQEKLNGVEINGRVVRVELVAGNRWYTKSGRKNHK
>Tox.amb.Nix_285aa
MAIRESFEEIKSYIVYVGNIPANAFEKDIKNIFRECGMIEKICILRSRQHCQHKIAYVRF
RHARGARRAAGLHGSYMDDDSYLIVKLLGLPYSSHVLLHDCTVAVSYCGGDKLKYKTKGI
THLELYKHFSDYGRIETILCTHPTYTYVSFSQPSQARRALLKTESTINHQKIRVFSIARN
VLIPGSRNLAIKTIDDLMKIDVCIGFFGLSPQCTENDLRDILQTYKLPTNLILIRDAVTS
MSRQYAFVKFSNSTIAMAKTIAHTLNGIEIKGRKVRVDLVVDNRE
>Wyeomyia.smithii_Nix_319aa
MVSSETSSILLNHTGQNMEMCKIDEKFDAVKVLIVYIGNLPSYVQEQHIYRLLKSEKIFE
TDEAIKIISIPERRCAFLRFCSLVSVNNTRRLNGLKYNESILIVVPLSSSYHERVLLAEC
SISILNVYSQITMQQLFDSLSIYGDIKAAVKLTDTSFIFSFQEPYQAEETLKKSGDIIRK
IKFLVKSNDVNNMMLGVKLFNLFRNINICAYDIQFTKGNSKEMLRQKTKIKAFLCEDFVA
GLKPNKYNCLGIFGLPSKTTEQILREKFCCIDEKIKIVMIPDTNYCFLYFDTVFKAVMLK
YEPKNGFVDKENIRERNLF
```

#### B) Other sequences used in this study

| Abbreviation | Clade | Genus/species | Accession |
|---|---|---|---|
| An.ste.429aa | fle, femaleless | *Anopheles stephensi* | ASTE008781 |
| An.cul.417aa | fle, femaleless | *Anopheles culicifacies* | ACUA023238 |
| An.gam.420aa | fle, femaleless | *Anopheles gambiae* | AGAP013051 |
| An.far.422aa | fle, femaleless | *Anopheles farauti* | AFAF008202 |
| An.dar.414aa | fle, femaleless | *Anopheles darlingi* | ADAC008134 |
| An.alb.414aa | fle, femaleless | *Anopheles albimanus* | AALB002485 |
| An.atr.420aa | fle, femaleless | *Anopheles atroparvus* | AATE020440 |
| An.sin.420aa | fle, femaleless | *Anopheles sinensis* | ASIS015421 |
| Ae.aeg. | transformer 2 (alpha) | *Aedes aegypti* | AAEL006416 |
| C.qui. | transformer 2 (alpha) | *Culex quinquefasciatus* | CPIJ004538 |
| An.gam. | transformer 2 (alpha) | *Anopheles gambiae* | AGAP006798 |
| An.ste. | transformer 2 (alpha) | *Anopheles stephensi* | ASTE010411 |
| An.dar. | transformer 2 (alpha) | *Anopheles darlingi* | ADAC007816 |
| An.sin. | transformer 2 (alpha) | *Anopheles sinensis* | ASIC019630 |
| An.far. | transformer 2 (alpha) | *Anopheles farauti* | AFAF004041 |
| An.fun. | transformer 2 (alpha) | *Anopheles funestus* | AFUN001838 |
| Ae.aeg. | transformer 2 (beta) | *Aedes aegypti* | AAEL004293 |
| C.qui. | transformer 2 (beta) | *Culex quinquefasciatus* | XP_038111372.1 |
| An.gam. | transformer 2 (beta) | *Anopheles gambiae* | AGAP029421 |
| An.ste. | transformer 2 (beta) | *Anopheles stephensi* | ASTE014045 |
| An.sin. | transformer 2 (beta) | *Anopheles sinensis* | ASIS021243 |
| An.far. | transformer 2 (beta) | *Anopheles farauti* | AFAF014144 |
| An.fun. | transformer 2 (beta) | *Anopheles funestus* | AFUN005410 |
| A.mel. | transformer 2 | *Apis mellifera* | NP_001252514.1 |
| T.cas. | transformer 2 | *Tribolium castaneum* | XP_008197650.1 |

| | | | |
|---|---|---|---|
| B.mor. | transformer 2 | *Bombyx mori* | NP_001119705 |
| D.mel. | transformer 2 | *Drosophila melanogaster* | NP_476764 |
| D.vir. | transformer 2 | *Drosophila virilis* | XP_002049699.1 |
| M.dom. | transformer 2 | *Musca domestica* | XP_011293246 |
| L.cup. | transformer 2 | *Lucilia cuprina* | ACS34688.1 |
| C.cap. | transformer 2 | *Ceratitis capitata* | NP_001266337 |
| B.ole. | transformer 2 | *Bactrocera oleae* | CAD67988.1 |
| A.fra. | transformer 2 | *Anastrepha fraterculus* | CBJ17284.1 |

#### C) Alignments used for phylogenetic analysis

Nexus file of alignment used for phylogeny in Fig 4

```
#NEXUS

BEGIN TAXA;
       DIMENSIONS NTAX=25;
       TAXLABELS
              Ae.aeg.288aa
              Ae.mas.288aa
              Ae.alb.282aa
              Ae.alb.d.205aa
              Ae.pol.282aa
              Ae.riv.282aa
              Ae.vex.293aa
              Ae.atr.271aa
              Ae.tri.278aa
              Ae.tri.276aa
              Ae.tri.204aa
              Ae.det.275aa
              Ae.jap.273aa
              Ps.col.277aa
              Wy.smi.319aa
              To.sp.287aa
              To.amb.285aa
              An.gam.420aa
              An.ste.429aa
              An.dar.414aa
              An.sin.420aa
              An.far.422aa
              An.cul.417aa
              An.atr.422aa
              An.alb.414aa
              ;
END;

BEGIN CHARACTERS;
       DIMENSIONS NCHAR=393;
       FORMAT datatype=protein missing=. gap=- interleave;
MATRIX
Ae.aeg.288aa    AEFDIIKKYCIYIGNIPFFASKNDVVVKFAEY-GETCNIYMQ-SNKPHCD
Ae.mas.288aa    AEFDIIKKYCIYIGNIPFFASKNDVVLKFAEY-GETCNIYMQ-SNKPHCD
Ae.alb.282aa    NQFEYIKKYCIYIGNIPAEVSKTDLIAKFSVF-GEISNLYMK-SFIQFCD
Ae.alb.d.205aa  ------------------------ILVEYSVF-CEISNVYMK-SFIQFCD
Ae.pol.282aa    KQFEYIKKYCIYIGNIPAEVSKRDLVAKFSEF-GEISNLYLK-SFVKFCD
Ae.riv.282aa    KQFECIKKYCIYIGNIPAEVSKRDLVAKFSEF-GEISNLYMK-SFVRFCD
```

```
Ae.vex.293aa     TEFEIIAQNCIYIGNIPNFVSKKILFDLFTQSGGEIFNIYIQ--QNEHCD
Ae.atr.271aa     EEFLKIKKNVVYVGNIPLNTTRNDILNLFKTY-GEVCTI----SEHKTAS
Ae.tri.278aa     EEFREIKENIVYIGNIPIDSSKDEIINLMKEY-GDIWNIYLQSNEEAMCN
Ae.tri.276aa     EEFREIKENIVYIGNIPIDSSKDEIINLMKEY-GDIWNIYLQSNEEAMCN
Ae.tri.204aa     KRFSDIEDYVVYVGNIPFDASKQDIIRLLGYY-GTIRNLYLQNTKESVCN
Ae.det.275aa     IKFNKIKGNVLYVGNIPISASKKELTCLFGTY-GEICTLYLLTSSEFVCD
Ae.jap.273aa     EKFDQMKNKIVYIGNIPLGASKKDIFMLLEEY-GKIFTI----SENQNHA
Ps.col.277aa     SEFDEISSRIIYVGNIPKSVNERQIRLLFRDC-GEIIRIAFT-WCYEFCS
Wy.smi.319aa     EKFDAVKVLIVYIGNLPSYVQEQHIYRLLKSE-KIFETDEAI-KIISIPE
To.sp.287aa      ESFEEIKSHIVYVGNVPAECSEEDIKNIFREC-GKVEKIVFA-CARDKCA
To.amb.285aa     ESFEEIKSYIVYVGNIPANAFEKDIKNIFREC-GMIEKICIL-RSRQHCQ
An.gam.420aa     QLFDELLENIVYVGNLPKQIMLTDLIELFKYA-GRIERVAWYGEHRTDIN
An.ste.429aa     QLFEELLENIVYVGNLPKNIVLTDLIELFKYA-GRIERVAWYGEHRTDIN
An.dar.414aa     QLFEELHGHIVYVGNLPKSTLLTELIELFKFA-GRIERVAWYGEHRTDIN
An.sin.420aa     HLFEELVEHIVYVGNLPKETVLTDLIELFKYA-GRIERVAWYGEHRTDIN
An.far.422aa     HLFDQLLGNIVYVGNLPKEIVLTDLIELFKYA-GRIERVAWYGEHRTDIN
An.cul.417aa     QLFDELLENIVYVGNLPKTIMLTDLIELFKYA-GRIERVAWYGEHRTDIN
An.atr.422aa     QLFEELLEHIVYVGNLPMNTALIDLTELFKYA-GRIERVAWYGDHRTDIN
An.alb.414aa     QLFEELHGHIVYVGNLPKSTLLTELIELFKFA-GRIERVAWYGEHRTDIN

Ae.aeg.288aa     VKPAIVRYRSRKSVDKSLC-LNNSKF----GNTILIVLPLSLPYSRYLLT
Ae.mas.288aa     VKPAIVRYRSRRSVDKSLC-LNNSKF----GNTILMVLPLSLPYSRYLLT
Ae.alb.282aa     VKPAVVRYRLMKSVKESSS-LHNSRY----IQSVLIVLPLDSSYNNYFLP
Ae.alb.d.205aa   VKSSVGRYRLIKYLKXSSS-LHNSQD----IQSILIVLQLYTYHKLGMLD
Ae.pol.282aa     VKHAVIRYRLETSVKESSS-LHNNRY----IQSVLIVLPLDSSYTHYYLP
Ae.riv.282aa     VKHAVIRYRLKGSVKESSS-LHNSRY----IQSVLIVLPLGLSYTHYYLP
Ae.vex.293aa     VKAAIIRFLHKKSVMRSLS-LNKTRY----HQSILIVIELSLPYANYLLV
Ae.atr.271aa     VKIAYLRYEKYEHVKSSVQSLDNKQY----GQTILIVKQLEMPYWYYILP
Ae.tri.278aa     VKIAYVRFMNKCDAKNCTKKLNNKEY----RNSVLILNNLRSPYTYNVLT
Ae.tri.276aa     VKIAYVRFMNKCDAKNCTKKLNNKEY----RNSVLILNNLRSPYTYNVLT
Ae.tri.204aa     VKFAYVRFLHSRHARNCARILNNKQY----RQSILIVTPMSEPLSFNLLI
Ae.det.275aa     VKSAYVRFEKSEDAQICETKLNNARY----RQSVLIVRIMAKPYSYYVLP
Ae.jap.273aa     VKTAYVRFCDPESVDKCLE-KNNTTY----CESILIVKRMAMPYSYYLLP
Ps.col.277aa     TKVCFIKFKLPRSTIKALK-FNRSNF----EDSFIIVLPVN-DNNDYLLT
Wy.smi.319aa     RRCAFLRFCSLVSVNNTRR-LNGLKY----NESILIVVPLSSSYHERVLL
To.sp.287aa      FKVAYVRFDLPRSSRKAAK-LHGSIVKLLGLPYSIIVKLLGLPYSSYVLM
To.amb.285aa     HKIAYVRFRHARGARRAAG-LHGSYM---DDDSYLIVKLLGLPYSSHVLL
An.gam.420aa     TKVAFIRFRHSRHAKEAAK-WDRIRY----QDSILIVMQI---FKDQWFD
An.ste.429aa     TKVAFVRFRHSRHAKEAAK-WDRIRY----HDSILIVMQI---FKDQWFD
An.dar.414aa     TKIAFIRFRTARHAKEAAK-WDRVRY----FDSVLIVMQI---FKDQWFD
An.sin.420aa     TKVAFIRFRHARHAKEAAK-WDRVRY----HDSILIVMQI---LKDQLFD
An.far.422aa     TKVAFIRFRHSRHAKEAAK-WDRIRY----HDSILIVMQI---FKDQWFD
An.cul.417aa     TKVAFVRFRHSRHAKEAAK-WDRIRY----HDSILIVMQI---FKDQWFD
An.atr.422aa     TKVAFIRFRHTRHAKEAAK-WDRVRY----HDSVLIVMQI---IKDQWFD
An.alb.414aa     TKIAFIRFRTARHAKEAAK-WDRVRY----FDSVLIVMQI---FKDQWFD

Ae.aeg.288aa     YDTCIVVYI--------NDKNSCKTSMAELYDEFQKIGDIQNMFK-TTNN
Ae.mas.288aa     YDTCIVVYI--------NDKNSCKTSMAELYDEFQKIGDIQNMFK-TTNN
Ae.alb.282aa     YNTCVVVYT--------YN----KFGMVDFYQKFSKLGDIHAMKK-ATNV
Ae.alb.d.205aa   FHIKXSV----------------------------NLGDIH-----VTNV
Ae.pol.282aa     YNTCVAVYT--------NK----HFHMVELFENFKRFGDIQVIKK-TTNV
Ae.riv.282aa     YNTCVAVYT--------NK----NFDMVELSENLRRFGAIQVIKK-TTNV
Ae.vex.293aa     YNTLIVIYI--------REKTK-KYSLEQIYDEFKSFGAIRNILK-TTNL
Ae.atr.271aa     YETTCIVSN--------FKE---STELVEIKNVFGAFGKISQILK-TTHT
Ae.tri.278aa     YESTVVISN--------FK----DTHLAAIKNAYEAFGKIHTILK-TTNI
Ae.tri.276aa     YESTVVISN--------FK----NTHLAAIKNAYEAFGKIHTILK-TNN-
Ae.tri.204aa     YETTVAVSN--------FE----DASIAKIKRVYERFGPVHVILK-RTNT
Ae.det.275aa     YETTVVISN--------FEE---DTDLVKIKSAYETFGKIESVLK-TSNT
Ae.jap.273aa     VETTVLVST------ELQEK---NITLKDIHGIFKCFGELFCILQ-RTDT
Ps.col.277aa     YETTVVVSN--------IHP---GMTLWAIYLLFKPFG-VKTVLK-TTDT
Wy.smi.319aa     AECSISILN--------VYS---QITMQQLFDSLSIYGDIKAAVK-LTDT
To.sp.287aa      HDCTVAVFYCG------KGT---GITHLELYKYFSEYGHIETIWCSTYNI
To.amb.285aa     HDCTVAVSYCGGDKLKYKTK---GITHLELYKHFSDYGRIETILC-THPT
```

```
An.gam.420aa    MSVSIMVRN--------IRD---DTTDWQLYEAFRRFGKIYGILI-PTHG
An.ste.429aa    MSVSIMVRN--------IRD---DTSDWQLYEAFRRFGKIYGILI-PTHG
An.dar.414aa    MSVSIIVRN--------IRD---DTTDWQLYEAFRRFGKIYGILI-PTHG
An.sin.420aa    MSVSIIVRN--------IRD---DTTDWQLYEAFRRFGKIYGILI-PTHG
An.far.422aa    MSVSIMVRN--------IRD---DTTDWQLYEAFRRFGKIYGILI-PTHG
An.cul.417aa    MSVSIMVRN--------IRD---DTTDWQLYEAFRRFGKIYGILI-PTHG
An.atr.422aa    MSVSIIVRN--------IRD---DTTDWQLYEAFRRFGKIYGILI-PTHG
An.alb.414aa    MSVSIIVRN--------IRD---DTTDWQLYEAFRRFGKIYGILI-PTHG

Ae.aeg.288aa    MIYINFESEKSMQLSLATKPFLINN--------------NIFKIKKVERN
Ae.mas.288aa    MIYINFESEKSMQLSLATKPFLINN--------------NIFKINKVERN
Ae.alb.282aa    MVYISFVSERAARTILDTKPTDIHI--------------NVQTINHVTRN
Ae.alb.d.205aa  TVYISFVSERAARVILDTNPTDTNN--------------------YVSRN
Ae.pol.282aa    MGYICFTTENAARKMLVTKPTDIHK--------------NVQKINDVTRN
Ae.riv.282aa    MGYICFTSENAARNMLVTKPTDIHK--------------NVQKINDVTRN
Ae.vex.293aa    MVYINYYSKKAQESALKSKS--LPD--------------YVDKIITSHRN
Ae.atr.271aa    FVYVCFYTKASAQLVLSTYVN-------------------GWKVSSTRRN
Ae.tri.278aa    FVFVSYYKKNSAELALSKRVT------------------KDWKISSIRRN
Ae.tri.276aa    FVFVSYYKQNSTELALSKGVT------------------KDWKTSNIRRN
Ae.tri.204aa    FVLVSYCDPESAEKALAKELS-------------------GFRTTSIMGN
Ae.det.275aa    FVFLAFYHKSSAQLALTKPLP-------------------YWKTSSIKRN
Ae.jap.273aa    FVYVSFCSEESAQISLSRSIK-IGG--------------CPIKVGQIYRN
Ps.col.277aa    FVYLSLETEQKTKELLQCTTS-WTA--------------NTVKVCSIRRN
Wy.smi.319aa    SFIFSFQEPYQAEETLKKSGDIIRKIKFLVKSNDVNNMMLGVKLFNLFRN
To.sp.287aa     LQYVQFSNPQQAQRALLMTETMINH--------------RKIRVFSINKN
To.amb.285aa    YTYVSFSQPSQARRALLKTESTINH--------------QKIRVFSIARN
An.gam.420aa    TAYVGFYYEAETQCALEMNNNMFNG--------------NRMRVEMLRRN
An.ste.429aa    TAYVGFYYEGETQCALEMNNNMFNG--------------NRMRVEMLRRN
An.dar.414aa    TAYVGFYYEEETQRALEMDNNMFNG--------------NRMRVAMLRRN
An.sin.420aa    TAYVGFYYEEETQRALEMDNNMFNG--------------NRMRVAMLRRN
An.far.422aa    TAYVGFYYEGETQCALEMNNNMFNG--------------NRMRVEMLRRN
An.cul.417aa    TAYVGFYYETETQCALEMNNNMFNG--------------NRMRVEMLRRN
An.atr.422aa    TAYVGFYYEEETQRALEMDNNMFNG--------------NRMRVAMLRRN
An.alb.414aa    TAYVGFYYEEETQRALEMDNNMFNG--------------NRMRVAMLRRN

Ae.aeg.288aa    INM--CGLNSESQDFST---NLKKKLL-----------------------
Ae.mas.288aa    INM--CGLNLESQDFST---NLKKKLL-----------------------
Ae.alb.282aa    INV--CLIDFEKECTSNT--AIKLTLL-----------------------
Ae.alb.d.205aa  IYV--CLI------------------------------------------
Ae.pol.282aa    INV--CLIDFDNESSGNT--AIKLTLL-----------------------
Ae.riv.282aa    INV--CLIDFDNESSGNT--AIKLTLL-----------------------
Ae.vex.293aa    IHM--CCINLDSKDFSRNL-EIKIKLL-----------------------
Ae.atr.271aa    INVINDQFDVEYHTPVTI--KIIDHLL-----------------------
Ae.tri.278aa    INI--RIIDIEHDKNGKTE-RIIKNLL-----------------------
Ae.tri.276aa    INI--RVIDIVHNKKG-AE-KIIINLV-----------------------
Ae.tri.204aa    VSS--RIIDIEQDENQKTK-KIIKILS-----------------------
Ae.det.275aa    INV-HQTIDIQQEETKEERYKIMKILL-----------------------
Ae.jap.273aa    INC--RLFDIEYKINQTTE-AIIKEMV-----------------------
Ps.col.277aa    VRL--ILKDIFSDD-EEMR-EIRSYLKDED--------------------
Wy.smi.319aa    INI--CAYDIQFTKGNSKE-MLRQKTK-----------------------
To.sp.287aa     LNL-------PGTRNLAVK-TINDLMK-----------------------
To.amb.285aa    VLI-------PGSRNLAIK-TIDDLMK-----------------------
An.gam.420aa    LPL--QQIDIFDITKNEVR-LLRDMLLEVDEKLEQFYSAYDEAFNYSSKD
An.ste.429aa    LPL--QQIDIFDITKNEVR-LLRDMLLEVDEKLEQFYSAYDEAFNYSSKD
An.dar.414aa    LPL--QQIDIFDTARNDVR-LLRDMLLEVDEKLEQFYSAYDEAFNYSSKE
An.sin.420aa    LSL--QQIDIHDTAKNEVR-LLRDMLLEVDEKLEQFYTAYDDAFNYSSKE
An.far.422aa    LPL--QQIDIFDITKNEVR-LLRDMLLEVDEKLEQFYSAYDEAFNYSSKD
An.cul.417aa    LPL--QQIDIFDMSRNEVR-LLRDMLLEVDEKLEQFYSAYDEAFNYSSKD
An.atr.422aa    LPL--QQIDIFDTAKNGVR-LLRDMLLEVDEKLEQFYTAYDDAFNYSSKD
An.alb.414aa    LPL--QQIDIFDTARNDVR-LLRDMLLEVDEKLEQFYSAYDEAFNYSSKE

Ae.aeg.288aa    --------------------------------------------------
Ae.mas.288aa    --------------------------------------------------
```

```
Ae.alb.282aa     --------------------------------------------------
Ae.alb.d.205aa   --------------------------------------------------
Ae.pol.282aa     --------------------------------------------------
Ae.riv.282aa     --------------------------------------------------
Ae.vex.293aa     --------------------------------------------------
Ae.atr.271aa     --------------------------------------------------
Ae.tri.278aa     --------------------------------------------------
Ae.tri.276aa     --------------------------------------------------
Ae.tri.204aa     --------------------------------------------------
Ae.det.275aa     --------------------------------------------------
Ae.jap.273aa     --------------------------------------------------
Ps.col.277aa     -------------------------------------------VSCTT--
Wy.smi.319aa     ----------------------------------------IKAFLCEDFV
To.sp.287aa      --------------------------------------------------
To.amb.285aa     --------------------------------------------------
An.gam.420aa     YRRWKRKRRRVTPLLSDSS-SSSESSSSSVSEISNRSATEVRRILCGS--
An.ste.429aa     YRRWKRKRRRVTPLLSDSS-SSSESSSSSVSEMSNRSATEVRRILCGS--
An.dar.414aa     YRRWKRKRRRVTPLLSESSDTSSESSASTVSVISNRSVTDVRRILCAS--
An.sin.420aa     YRKWKRKRRRVTPLLSESS-SSSESSSSSVSVISNRSATEVKRIMCGS--
An.far.422aa     YRRWKRKRRRVTPLLSDSS-SSSESSSSSVSEISNRSATEMRRIMCGS--
An.cul.417aa     YRRWKRKRRRVTPLLSDSS-SSSESSSSSVSEMSNRSANEVRRILCGS--
An.atr.422aa     YRKWKRKRRRVTPLLSESSDSSSESSTSSVSVISNRSVTEVKRIMCGS--
An.alb.414aa     YRRWKRKRRRVTPLLSESSDTSSESSASTVSVISNRSVTDVRRILCAS--

Ae.aeg.288aa     ------YNRSIGIFGLPSNATEERIAQIFS-RFGDIQKITLICDVVGN-S
Ae.mas.288aa     ------YNRSIGIFGLPSNATEERIAQIFS-RFGDIQKITLICDIVGN-S
Ae.alb.282aa     ------YNRSIGIFGLPSNFTEAKLHDEFS-RYGRIEKNRLVYDSTGH-S
Ae.alb.d.205aa   -------VCSIEIFGLPSDFTESKLHDEFXRRYGSIEKSRLVYDSTGN-S
Ae.pol.282aa     ------YNRSIGVFGLPSDFTEEKLYDEFS-RYGRIEKNKLVYDSTGH-S
Ae.riv.282aa     ------YNRSIGIFGLPSDFTEAKLHDEFS-RYGRIEKNKLVYDSTGH-S
Ae.vex.293aa     ------YSSTIGIFGLPSDTKEAEITQICS-RFGDIDKIKLIYDKGGN-S
Ae.atr.271aa     ------HPRCLGIFGLSKATDEKSIEGLFK-RYGNIKSIKLVLNRQKI-S
Ae.tri.278aa     ------YSRCLGIFNLHQNTTAKRIIDFFE-KYGIITKTKLILSKEGN-S
Ae.tri.276aa     ------NSRCLGVFNLHQNATKNRINDCFE-KYGEITNTKLIVSKEGY-S
Ae.tri.204aa     ------DHRYLGQY------------------------------------
Ae.det.275aa     -G----YQRCVGVFGLSQQTTQTSIAEKMD-KYGKITNIKIPVTAQGI-S
Ae.jap.273aa     ------YPRCLGIFGLSKETTETSIAKHFS-KYAPIYKCVLIRTPEGN-S
Ps.col.277aa     ------TTRVLGCFGLDPKTSEDKIMQIFS-RYGHIENVRLIRNRKNSVS
Wy.smi.319aa     AGLKPNKYNCLGIFGLPSKTTEQILREKFC-CIDEKIKIVMIPDTN----
To.sp.287aa      ------IDVCIGFFGLSAQTTETDLRNLLQ-PYK-ILAVKLIRNS-----
To.amb.285aa     ------IDVCIGFFGLSPQCTENDLRDILQ-TYKLPTNLILIRDAVTSMS
An.gam.420aa     ----NEENRCLGIFGMNPDTTEKTLMKLFS-RYGHVKDIKLIYDGKTNVS
An.ste.429aa     ----NEENRCLGIFGMNPDTTEKTLMKLFS-RYGHVKDIKLIYDGKTNVS
An.dar.414aa     ----NEENRCLGIFGMSPETTENTLMKLFS-RYGQVKDIKLIYDGKTNVS
An.sin.420aa     ----NEENRCLGIFGMSPETTEKKLMKLFS-RYGQVKDIKLIYDGKTNVS
An.far.422aa     ----NEENRCLGIFGMNPDTTEKTLMKLFS-RYGHVKDIKLIYDGKTNVS
An.cul.417aa     ----NEENRCLGIFGMNPDTTEKTLMKLFS-RYGHVKDIKLIYDGKTNVS
An.atr.422aa     ----NEENRCLGIFGMSPETTEKKLMKLFS-RYGQVKDIKLIYDGKTNVS
An.alb.414aa     ----NEENRCLGIFGMSPETTENTLMKLFS-RYGQVKDIKLIYDGKTNVS

Ae.aeg.288aa     KQ-YGFIYYKKR--TSANAAKVIMDGENFEGNKISVRFVPEKK
Ae.mas.288aa     KQ-YGFIYYKKR--TSANAAKVIMDGENFEGNIISVRFVPEKK
Ae.alb.282aa     KQ-YGFVYYEKH--LSAQAAKQEMDRSDHTGRKIAVRFVPEKE
Ae.alb.d.205aa   KXTIWFCLLXNX--LFTRAEKQEMDRSYYTERKIAVRFVTEKE
Ae.pol.282aa     KQ-YGFVYYEKH--ISAQAAKEEMDRSDYTGGKISVRFVPEKE
Ae.riv.282aa     RQ-YGFVYYEKH--LSAQAAKEEMDRSDYTGGKISVRFVPEKE
Ae.vex.293aa     KQ-YCFVYYKNH--ISAIEAKHNLDKKPLQGREISVRFVPEKE
Ae.atr.271aa     KQ-YCFITFVDP--LSARMAQKALDGIVIDGRIISVRYTLER-
Ae.tri.278aa     LQ-YCFIYFDNP--LSAEKARKALDDTNIDGRVISVRSTAERE
Ae.tri.276aa     LQ-YCFIYFDDP--LSAEKARKALDETYIDGRVISVRCTAERE
Ae.tri.204aa     -------------------------------------------
Ae.det.275aa     KQ-YCFVYFDDP--RSAQAARLGLDKTIIDGRQISVRAISEKE
Ae.jap.273aa     RR-FGFLYFDDP--QSAQAALADLDESNIDGCKISVRFAPEKY
```

```
Ps.col.277aa     MG-YCFIFFDNS--DSASNAYSDLKELVIDDRK----------
Wy.smi.319aa     ---YCFLYFDTV--FKAVMLKYEPKNGFVDKENIRERNLF---
To.sp.287aa      ---YAFVKFEFANANFAKLAQEKLNGVEINGRVVRVELVAGNR
To.amb.285aa     RQ-YAFVKFSNSTIAMAKTIAHTLNGIEIKGRKVRVDLVVDNR
An.gam.420aa     RG-YSFIYFKHA--SDARRAQRKLNGTMLEGRKVRVDFSRSKP
An.ste.429aa     RG-YSFIYFKHA--SDARRAQRKLNGTMLEGRKVRVDFSRSKP
An.dar.414aa     RG-YSFIYFKHA--SDARRAQRKLNGTMLDGRKVRVDFSRSKP
An.sin.420aa     RG-YSFIYFKHA--SDARRAQRKLNGTMLDGRKVRVDFSRSKP
An.far.422aa     RG-YSFIYFKHA--SDARRAQRKLNGTMLEGRKVRVDFSRSKP
An.cul.417aa     RG-YSFIYFKHA--SDARRAQRKLNGTMLEGRKVRVDFSRSKP
An.atr.422aa     RG-YSFIYFKHA--SDARRAQRKLNGTMLDGRKVRVDFSRSKP
An.alb.414aa     RG-YSFIYFKHA--SDARRAQRKLNGTMLDGRKVRVDFSRSKP

;
END;
```

#### Nexus file of alignment used for phylogeny in Fig 8

```
#NEXUS

BEGIN DATA;
DIMENSIONS  NTAX=49 NCHAR=94;
FORMAT DATATYPE=PROTEIN GAP=- MISSING=?;
MATRIX

Ae_aeg_288aa             KLL-YNRSIGIFGLPSNATEERIAQIFS-RFGDIQKITLI----CDV-VGNSKQ-YGFIYYK--
KRTSANAAKVIMDGENFEGNKISVRFVPEK
Ae_mas_288aa             KLL-YNRSIGIFGLPSNATEERIAQIFS-RFGDIQKITLI----CDI-VGNSKQ-YGFIYYK--
KRTSANAAKVIMDGENFEGNIISVRFVPEK
Ae_alb_282aa             TLL-YNRSIGIFGLPSNFTEAKLHDEFS-RYGRIEKNRLV----YDS-TGHSKQ-YGFVYYE--
KHLSAQAAKQEMDRSDHTGRKIAVRFVPEK
Ae_alb_d_205aa           ----IVCSIEIFGLPSDFTESKLHDEFXRRYGSIEKSRLV----YDS-TGNSKXTIWFCLLX--
NXLFTRAEKQEMDRSYYTERKIAVRFVTEK
Ae_pol_282aa             TLL-YNRSIGVFGLPSDFTEEKLYDEFS-RYGRIEKNKLV----YDS-TGHSKQ-YGFVYYE--
KHISAQAAKEEMDRSDYTGGKISVRFVPEK
Ae_riv_282aa             TLL-YNRSIGIFGLPSDFTEAKLHDEFS-RYGRIEKNKLV----YDS-TGHSRQ-YGFVYYE--
KHLSAQAAKEEMDRSDYTGGKISVRFVPEK
Ae_vex_293aa             KLL-YSSTIGIFGLPSDTKEAEITQICS-RFGDIDKIKLI----YDK-GGNSKQ-YCFVYYK--
NHISAIEAKHNLDKKPLQGREISVRFVPEK
Ae_atr_271aa             HLL-HPRCLGIFGLSKATDEKSIEGLFK-RYGNIKSIKLV----LNR-QKISKQ-YCFITFV--
DPLSARMAQKALDGIVIDGRIISVRYTLER
Ae_tri_278aa             NLL-YSRCLGIFNLHQNTTAKRIIDFFE-KYGIITKTKLI----LSK-EGNSLQ-YCFIYFD--
NPLSAEKARKALDDTNIDGRVISVRSTAER
Ae_tri_276aa             NLV-NSRCLGVFNLHQNATKNRINDCFE-KYGEITNTKLI----VSK-EGYSLQ-YCFIYFD--
DPLSAEKARKALDETYIDGRVISVRCTAER
Ae_det_275aa             ILLGYQRCVGVFGLSQQTTQTSIAEKMD-KYGKITNIKIP----VTA-QGISKQ-YCFVYFD--
DPRSAQAARLGLDKTIIDGRQISVRAISEK
Ae_jap_273aa             EMV-YPRCLGIFGLSKETTETSIAKHFS-KYAPIYKCVLI----RTP-EGNSRR-FGFLYFD--
DPQSAQAALADLDESNIDGCKISVRFAPEK
Ps_col_277aa             CTT-TTRVLGCFGLDPKTSEDKIMQIFS-RYGHIENVRLI----RNRKNSVSMG-YCFIFFD--NSDSASNAYSDLKELVIDDRK-
--------
Wy_smi_319aa             KPN-KYNCLGIFGLPSKTTEQILREKFC-CIDEKIKIVMI----PDTN-------YCFLYFD--
TVFKAVMLKYEPKNGFVDKENIRERNLF--
To_sp_287aa              LMK-IDVCIGFFGLSAQTTETDLRNLLQ-PYK-ILAVKLI----RNS--------
YAFVKFEFANANFAKLAQEKLNGVEINGRVVRVELVAGN
To_amb_285aa             LMK-IDVCIGFFGLSPQCTENDLRDILQ-TYKLPTNLILI----RDAVTSMSRQ-
YAFVKFSNSTIAMAKTIAHTLNGIEIKGRKVRVDLVVDN
An_gam_420aa             SNE-ENRCLGIFGMNPDTTEKTLMKLFS-RYGHVKDIKLI----YDGKTNVSRG-YSFIYFK--
HASDARRAQRKLNGTMLEGRKVRVDFSRSK
An_ste_429aa             SNE-ENRCLGIFGMNPDTTEKTLMKLFS-RYGHVKDIKLI----YDGKTNVSRG-YSFIYFK--
HASDARRAQRKLNGTMLEGRKVRVDFSRSK
An_dar_414aa             SNE-ENRCLGIFGMSPETTENTLMKLFS-RYGQVKDIKLI----YDGKTNVSRG-YSFIYFK--
HASDARRAQRKLNGTMLDGRKVRVDFSRSK
An_sin_420aa             SNE-ENRCLGIFGMSPETTEKKLMKLFS-RYGQVKDIKLI----YDGKTNVSRG-YSFIYFK--
HASDARRAQRKLNGTMLDGRKVRVDFSRSK
An_far_422aa             SNE-ENRCLGIFGMNPDTTEKTLMKLFS-RYGHVKDIKLI----YDGKTNVSRG-YSFIYFK--
HASDARRAQRKLNGTMLEGRKVRVDFSRSK
An_cul_417aa             SNE-ENRCLGIFGMNPDTTEKTLMKLFS-RYGHVKDIKLI----YDGKTNVSRG-YSFIYFK--
HASDARRAQRKLNGTMLEGRKVRVDFSRSK
```

```
An_atr_422aa             SNE-ENRCLGIFGMSPETTEKKLMKLFS-RYGQVKDIKLI----YDGKTNVSRG-YSFIYFK--
HASDARRAQRKLNGTMLDGRKVRVDFSRSK
An_alb_414aa             SNE-ENRCLGIFGMSPETTENTLMKLFS-RYGQVKDIKLI----YDGKTNVSRG-YSFIYFK--
HASDARRAQRKLNGTMLDGRKVRVDFSRSK
Ae_aeg_tra2alpha_272aa   YDE-SKAVLAVFNLNIYTTESELYDVFT-KFGPLKKATIV----LDAKTGRSRG-FGFVYFE--
STEDARVAHTQANGIEIGDRPIRVDYSATE
Cu_qui_200aa             --------------------------FS-------KSKLISWKQHQVGTGRSRG-FGFVYFE--
SIEDARVAHVQANGIEIGDRRIRVDYSATD
An_gam_271aa             STS-GKVVLAVFNLSVYTTEAELYDTFS-KFGPLRKTTVV----LDAKTGRSRG-FGFVYFE--
SAEDAKVAHDQANGIEIGDRRIRVDFSATN
An_ste_271aa             GSS-GKLVLAVFNLSVYTTEAELYDTFS-KFGPLRKTTVV----LDAKTGRSRG-FGFVYFE--
SAEDAKVAHDQANGIEIGDRRIRVDFSATN
An_dar_255aa             YSS-GKLVLAVFNLSIYTTEAELYDIFS-KFGPVRKTTVV----LDAKTGRSRG-FGFVYFE--
SAEDAKIAHDQANGIEIGDRRIRVDFSATN
An_sin_257aa             AQS-GKVVLAVFNLSIYTTEAELYDIFS-KFGPLRKTTVV----LDAKTGRSRG-FGFVYFE--
SAEDAKVAHDQANGIEIGDRRIRVDFSATE
An_far_276aa             GSS-GKVVLAVFNLSIYTTEAELYDTFS-KFGPLRKTTVV----LDAKTGRSRG-FGFVYFE--
SAEDAKVAHDQANGIEIGDRRIRVDFSATN
An_fun_269aa             GSS-GKVVLAVFNLSIYTTEAELYDTFS-KFGPLRKTTVV----LDAKTGRSRG-FGFVYFE--
SAEDAKVAHDQANGIEIGDRRIRVDFSATN
A_aeg_tra2beta_244aa     DPP-KSKCLGVFGLSSYTNETSLMDVFA-PYGTIDKAMIV----YDAKTKVSRG-FGFVYFQ--
EQSAATEAKMQCNGMMLHERTIRVDYSVTE
Cu_qui_237aa             DPP-PSTCLGVFGLSNYTQEADLRTVFG-RFGLIEKVQIV----YDAKTKASRG-FGFVYFV--
NLEDASAAKVQCNGMVMHERTIRVDYSVTE
An_gam_215aa             SPE-PSRCLGVFGLSVYTTEPYLNDIFC-HFGTVEKSVVI----YDAKTRLSRG-FGFVYFK--
SQAEASIARANCNGLQIHGRRIRVDYSITD
An_ste_210aa             SPE-PSRCLGVFGLSVFTTEPYLNRIFC-SFGTVENTVVI----YDAKTRLSRG-FGFVYFK--
TKEEAAVARAHCNGLHIHGRRMRVDYSITE
An_sin_175aa             SPE-PSRCLGVFGLSVYTTEPYLKDIFD-QYGIVEDVFVV----YDAKTRLSRG-FGFVYFQ--
NVAEASWARMHCNGLHVHGRRIRVDFSISD
An_far_217aa             SPA-PSRCLGIFGLSVYTTEPYLEDIFG-HFGTVERTFVI----YDAKTRLSRG-FGFVYFK--
TKEEASVARTHCNGLQIHGRRIRVDYSITD
An_fun_248aa             SPE-PSRCLGIFGLSVFTTEPYLNRIFC-NFGTVENTVVI----YDAKTRLSRG-FGFVYFK--
TKEEAAVARAHCNGLHIHGRRMRVDYSITD
D_mel_264aa              HPQ-ASRCIGVFGLNTNTSQHKVRELFN-KYGPIERIQMV----IDAQTQRSRG-FCFIYFE--
KLSDARAAKDSCSGIEVDGRRIRVDFSITQ
C_cap_251aa              KPV-QNRCIGVFGLSVYTTQQKIRDIFS-RFGPIERIQVV----IDAQTGRSRG-FCFIYYD--
DIADAKAAKDACSGMEIDDRRIRVDYSTTQ
M_dom_232aa              KPS-PCRCLGVFGLSVHTTQQQIREIFS-KYGPIERIQVV----VDAQTGRSRG-FCFIYYK--
HLADAEVARDQCCGQEVDGRRIRVAYSITE
B_mor_284aa              NPT-PSRCLGVFGLSLYTTEQQINHIFS-KYGPVDKVQVV----IDAKTGRSRG-FCFVYFE--
DMEDAKIAKNECTGMEIDGRRIRVDYSITQ
A_mel_252aa              NPS-PSRCLGVFGLSIFTTEQQVHHIFS-KYGPVERIQVV----IDAKTGHSKG-YCFVYFE--
SLEDAKVAKEQCAGMEIDGRRMRVDYSITQ
T_cas_275aa              NPK-PSRCLGVFGLSVYTTEDELYHIFS-KYGPLERVQVV----IDAKTGRSRG-FSFVYFE--
NTDDAKVAKDQCSGMKINGKNIRVDYSITE
D_vir_315aa              HPQ-ASRCIGVFGLNTNTTQQKVRELFN-KFGPIERIQMV----IDAHTHRSRG-FCFIYFE--
NLGDARVAKDACTGMEVDGRRIRVDYSITQ
L_cup_271aa              KPL-PCRCIGVFGLSVYTTQLKIREIFS-KFGPIERIQVV----IDAQTGRSRG-SCFIYYE--
NLADAKAACDNCCGMEIEGRRIRVAYSITE
A_fra_249aa              KPV-QNRCIGVFGLSVYTTQQKIRDIFS-RFGPIERIQVV----IDAQTGRSRG-FCFIYYQ--
DIADAKAAKDACSGMEIDDRRIRVDYSTTQ
B_ole_251aa              KPV-QNRCIGVFGLSVYTTQQKIRDIFS-RFGPIERIQVV----IDAQTGRSRG-FCFIYYE--
DIADAKAAKDACSGMEIDDRRIRVDYSTTQ
;

END;

BEGIN ASSUMPTIONS;
EXSET * UNTITLED  = ;
END;

BEGIN CODONS;
CODONPOSSET * CodonPositions =
 N:,
 1: 1-94\3,
 2: 2-92\3,
 3: 3-93\3;
CODESET  * UNTITLED = Universal: all ;
END;

BEGIN SETS;
END;
```

**Supplemental Data 2. Plasmid sequences used in this study**

**>187_Ae.jpn.Nix**
TTTCCCGACTGGAAAGCGGGCAGTGAGCGCAACGCAATTAATGTGAGTTAGCTCACTCATTAGGCACCCCAGGC
TTTACACTTTATGCTTCCGGCTCGTATGTTGTGTGGAATTGTGAGCGGATAACAATTTCACACAGGAAACAGCTAT
GACATGATTACGAATTCGAGCTCGGTACCCGGGGATCCTCTAGAGTCGACGCTCGCGCGACTTGGTTTGCCATTC
TTTAGCGCGCGTCGCGTCACACAGCTTGGCCACAATGTGGTTTTTGTCAAACGAAGATTCTATGACGTGTTTAAA
GTTTAGGTCGAGTAAAGCGCAAATCTTTTTTAACCCTAGAAAGATAGTCTGCGTAAAATTGACGCATGCATTCTT
GAAATATTGCTCTCTCTTTCTAAATAGCGCGAATCCGTCGCTGTGCATTTAGGACATCTCAGTCGCCGCTTGGAGC
TCCCGTGAGGCGTGCTTGTCAATGCGGTAAGTGTCACTGATTTTGAACTATAACGACCGCGTGAGTCAAAATGAC
GCATGATTATCTTTTACGTGACTTTTAAGATTTAACTCATACGATAATTATATTGTTATTTCATGTTCTACTTACGTG
ATAACTTATTATATATATATTTTCTTGTTATAGATATCGTGACTAATATATAATAAAATGGGTAGTTCTTTAGACGA
TGAGCATATCCTCTCTGCTCTTCTGCAAAGCGATGACGAGCTTGTTGGTGAGGATTCTGACAGTGAAATATCAGA
TCACGTAAGTGAAGATGACGTCCAGAGCGATACAGAAGAAGCGTTTATAGATGAGGTACATGAAGTGCAGCCA
ACGTCAAGCGGTAGTGAAATATTAGACGAACAAAATGTTATTGAACAACCAGGTTCTTCATTGGCTTCTAACAGA
ATCTTGACCTTGCCACAGAGGACTATTAGAGGTAAGAATAAACATTGTTGGTCAACTTCAAAGTCCACGAGGCGT
AGCCGAGTCTCTGCACTGAACATTGTCAGATCTaacccccttgtgtcatgtcggcgaccctacgcccccaactgagagaactcaaaggt
taccccagttggggcactactcccgaaaaccgcttctgacctgggTAAGATACATTGATGAGTTTGGACAAACCACAACTAGAAT
GCAGTGAAAAAAATGCTTTATTTGTGAAATTTGTGATGCTATTGCTTTATTTGTAACCATTATAAGCTGCAATAAA
CAAGTTAACAACAACAATTGCATTCATTTTATGTTTCAGGTTCAGGGGGAGGTGTGGGAGGTTTTTTAAAGCAAG
TAAAACCTCTACAAATGTGGTATGGCTGATTATGATCTAGCGGCCGCTTTACTTGTACAGCTCGTCCATGCCGAG
AGTGATCCCGGCGGCGGTCACGAACTCCAGCAGGACCATGTGATCGCGCTTCTCGTTGGGGTCTTTGCTCAGGG
CGGACTGGGTGCTCAGGTAGTGGTTGTCGGGCAGCAGCACGGGGCCGTCGCCGATGGGGGTGTTCTGCTGGTA
GTGGTCGGCGAGCTGCACGCTGCCGTCCTCGATGTTGTGGCGGATCTTGAAGTTCACCTTGATGCCGTTCTTCTG
CTTGTCGGCCATGATATAGACGTTGTGGCTGTTGTAGTTGTACTCCAGCTTGTGCCCCAGGATGTTGCCGTCCTCC
TTGAAGTCGATGCCCTTCAGCTCGATGCGGTTCACCAGGGTGTCGCCCTCGAACTTCACCTCGGCGCGGGTCTTG
TAGTTGCCGTCGTCCTTGAAGAAGATGGTGCGCTCCTGGACGTAGCCTTCGGGCATGGCGGACTTGAAGAAGTC
GTGCTGCTTCATGTGGTCGGGGTAGCGGCTGAAGCACTGCACGCCGTAGGTCAGGGTGGTCACGAGGGTGGGC
CAGGGCACGGGCAGCTTGCCGGTGGTGCAGATGAACTTCAGGGTCAGCTTGCCGTAGGTGGCATCGCCCTCGCC
CTCGCCGGACACGCTGAACTTGTGGCCGTTTACGTCGCCGTCCAGCTCGACCAGGATGGGCACCACCCCGGTGA
ACAGCTCCTCGCCCTTGCTCACCATGGTTGAAATCTCTGTTGAGCAGAAAAAGAAACGAGGAAACGCTTgAGTAA
TTGGTTGTGAAATGCAAACTCTCATTTGATATTGATTCATTGCCTTTGGCTTCGAGCACGACACGACAGGTTTTAA
ACTTGTTTTGCTTTGTCTGCGTTTGCAGTCGCAGGCCAAGTGAAAAATATACACTTGAAGGTGATGACGTCACAA
CAACGCCCTACTTTTaAGTGAAAAATTAACTTGTTTTCGACTTTGAACTACGTAGTTTtGAAATTGCGTATCTTCAA
GTTTTCACGATTTCCTCAAGGTTTTTCTCGATATGTGTTAATATTACCTTAATGGGTAATTACCATCAAAATATTTA
TTTTAGATATGTGACGGAGCAAATACGTTATTCTTATTATTCTAGAAATTTAATTCAATTAGtAGCGATGATTCAAC
GAAATATGATTATCGCtGTGAATCACAATTGGGTTTTATCAATGATGATGAAACTGCGTTGCAAATTTTCACTAAT
CACTCAAAGCTCAATAGTCGCCATCTTGGAAAATAGTTTGCTCATTCAAAGACAAAGGAATCATTCACCAAACTA
GTTTTCGCTCATAGCTATAATTTCATCAaTTAATTTACCTACCTTCACTAGAAGATTCCCTTCACCGAAATGCACTTT
CACcAATTTAATAGTAATTGTCCTTTGAAtAAAGCTTTGTTCACTCTGAAATTTTCTCCTCTGGCTAATTGGATCACT
CTTTTTCACTAGAGACTTCACTTCACTTGCACTGGCACTGCTTACTTGGgCCGCGTAATGTTCACTCCACTAGGAAA
CGTATTCGATTGAGCTGGTTTcGCCTTTGCAGGGGGCGTTTTATAGACACTGtCGTAGTGGTGGTTGTACTTCTAGA
AAATTTCaGCAATaCATTCATACATATACCTCTGTTCGGTTGGATGGCTCTAtATCGATCGAaTATGGGTACCATCC
CTGTGTCTGAATGGACAGCAAAACGTGCTTGTGTCTGTTAGTCGTTCATTCTGTACCTTGAACGATGCAGTTCAAC

TTCTGGCAAAGACGTCAATGTACCTACCcTTCGTGTATATGGCATAgAGAGgAAGATGTGCGAATGCCTTTTTCGA
TAGAGAAAGGATTTCCTGATTTGATCGACAATTTCGGTAGCTAaTCTTAGCTTCGAATGTAATCTGATAACCAAAT
CCAGAGAaAAAtCAGTAATTTGGGAATTTCACTTGAATCATTCTAATTGcATCATTTGCTGTATTAGTGCAGTCAGC
AAGTGACGTCAACCCTTCTAAATCGATATACTTCTGGGAAGCTTTCTTTCTTGTCTGGCTCAGCTGGTGCCAAGGC
AAATTATAATTGGATTCAATGCACAAGCTACATGTAAAGATActcgagAGAACAAAAACCGCTCCAGAAACTACTT
ACCTTGAAATGATATTTCAAATATTTTTGCTAGAAGGGTGTAGATCCGAATCCATACCCATATACGAATAGTTTCA
GAGTCGAAAAAGAATATTAAACATTCAAAACTAAATTTTAGTTAACAAAAAAACGCCACCGCAAAGTCAACTTGA
AAATGGCGATGCTGCTCGTCAAAGTATCGTGCGTATTATGTACCCATGATTAGTAAACGAATGGAAACTTATTTA
TGTGGATTAAGTACCCTTTTTATGACATTTGAAAAATACCCTCTTTCTGACAATTCAGTACTGGTTGGGAAAATGG
GTACATGGAACCCATTTAATGGGTACTTCCAAGTTAGCGTGAAGTTTACTGTCAATGTCATTCCAATCGAGCAGG
AGACTTGTCAAGCACTTTTTCACACAAGCTGAAAACGGCTCACACAAAAATGTTACCGCCCACGAAAATTTTGCTT
CTTCTGAAGTAGCATATTTCTGAATACTTGAGTGTCGGTTGCAATTTTCATGCTTTGCATAAAGAATGCTCTACTC
ATTACATTGGTTTTCCACACATTCAGTCAATTTTTCATTCGTTACCCAAATCTGTGAAAATTCAACACATAAAACTT
ATGACTCACTTTTAGCGTGGGATTCAACATTGGCAGTGCTGCGTAATAGATTTGGCACAGTTGCTAGCTGATTTA
TTTTGAATATGATCAGGTTTTAGGTAACGCTCGTAACGCTTTTTGTATGAATATTCTACAAATTTTGTATGAGCCGT
AACACCAGCAAAACACCCACCCACCCCCTTCAGCGTTATGAAATTTGTAAATAAGCCCATATGGGTAAAATATAC
CAATAAAAATTGGTAAATTTCACCCACATTATTAGTTTACTGGATTTACCCAAATATGGGTAGATGCGGTTACCCA
TATTTGGATGAAAAATTAGAATTGTTGTCAGTCAGAGTGCGGCGCTCAAGATGGAAATTCATTTTTCAGCAGTAC
TGAATAAATTGTTTTAGGATCAAGAAGAAAAATGATGATTCCTATCGGAGGTGAGTAGGTATAGTCATTTATTCA
TTGTTATTAGGTGTTATATATTCCAGTTTCTTCATCGATGTGCACCAATGTTTTTCTTCATGGCGACAAAAAACTGA
ACATTTCGGGCTGGTTAAATTGTGCTGTACTAACCAAGATGCAACAACTACAAAACTGAATCGAAATTGGGAGAC
AGCAATTCCAAATAAGAATGTCGTTCCGTTTATTTGTCACGATATACATATTTCACAACTCTTGGTTGACAATAATT
TTACTCTATTTTATAATATATCTTTATTTCTCAATAAAGTTTGCATTTGTATGTTAATGACAAAATTCAATTTAATTC
AAAATAGTCATAAAGAAATTTAATAAAAAAGCTAGTCGAAAAGTCTATGAAACTACTCATTTTTGCGTATTTTTTA
GAATAATTGCTTCCAAGCTTGAATACCCAAATATGGGTATTGTCAGTTTACCTATTTATGAGTAAACCCGCTTTTT
GGCGATTATGGGCAAACTTTACCCATATTTGGGTAGACTGTTCTTAGCGTGTACATCTAGGGCACCCATACCCAA
TGTGATCTATAAGAGGTGAAGAAAAAGTCATTGTGTGATTGTATTAGTACAATAAAAATCAAAACAAACATTGAG
GGGAAAATGAGTGACGTCATACCGTTTCAACGAAATGCTGTGAGTGGATGTAAAACACTCCATATCGTGGAACA
AATCAAGATCACACCGTATAAATCTGCAACTAACACCTAGATGAAAGAGTCGAATGGGTAGATTCCAGGAATTAT
GTTTTTTCACTATATTCACGGATGAAAGAAAGCACTTTTCTTACTTTTTTCTAATTATATTCAATTTGAGCACTACAT
TTATGGATGAATTACGCAAAATATCCTTTGGAAGAAAGAACAATTTCAGCCTATTCCAGCACATTATCAGTTTTGA
TGATATTTGGGTTGCCTTCCGTTGTTATATTTCCTCTGTTTTATAATAGATTTCATCCTAAAATAAAGCAAAACTTA
GCATGAAGCAAACAACGCATGAATTACCTGAAATGCCATCTTTCTGCGACGAATTTTTTCATGTTGTTGTTTGAAC
AACAGCACACAGGACAACCTTGGTAACGATAGGGTTGCTAGTAACGATACCTCGAAATAAATCGATAACAAACT
GTATGACGTCACGCAGTGGTGGAATGCAATGGGTTGCTGTGTGAAAATTTAAGCGACCCACTAACACATGCAAG
CAGTTGTGTATTACACAAAATCATCTCCTCACCTCTAATAAAGCACATTGACCCATACCATTGGGCGTGATAACCA
CTATAATCGAGTAAAAATAAGAGAGAACAGTGGCACGCAACATGATTTTTCCCATTTTTCCACCTGAGCAAACTA
AGAACATTGACTACGGAGCTCAGTCCAGCAATAATGTTAATCATAAAAAGATGTGATCTTTTTCAAAGAAAATTT
TATTAGTGTTTTCTAAGTTTTATATTGATTTTTGTTTTTGTCGTGCAAATGGAAAAATTCGATCAAATGAAAAATAA
AATAGTTTATATTGGAAATATTCCCCTAGGAGCTTCGAAAAAAGATATCTTTATGTTATTGGAGGAATATGGTAA
AATATTTACTATAAGTGAGAATCAAAACCATGCAGTGAAAACAGCCTATGTGCGGTTTTGTGATCCAGAATCTGT
AGATAAGTGTTTAGAAAAAAATAACACTACGTATTGTGAATCTATATTGATAGTAAAGAGAATGGCTATGCCATA
TTCATACTATTTATTACCCGTTGAAACAACGGTGTTAGTATCTACAGAACTACAGGAGAAAAACATAACGCTAAA
AGACATTCATGGCATATTTAAGTGTTTTGGCGAGTTATTTTGCATTTTACAAAGAACAGATACGTTTGTATACGTA

TCCTTTTGCTCTGAGGAAAGTGCACAGATTTCTTTATCACGATCTATTAAAATAGGTGGATGTCCAATTAAGGTTG
GTCAAATCTATAGAAACATCAATTGCCGGCTATTCGACATCGAATATAAAATCAACCAAACGACTGAAGCAATTA
TTAAAGAAATGGTGTATCCCCGATGCTTAGGGATTTTTGGGCTCTCCAAAGAGACAACAGAAACGAGTATTGCCA
AACATTTTTCCAAGTATGCACCCATATACAAGTGTGTTTTGATAAGAACACCAGAAGGAAATTCAAGAAGATTCG
GTTTTTTGTACTTCGACGATCCACAATCCGCACAAGCTGCCCTTGCTGACCTTGATGAAAGTAATATTGATGGATG
CAAAATTTCGGTACGTTTTGCTCCCGAAAAATATGATTAATAAAGTGATATATGACGCGGCCGCGACTCTAGATC
ATAATCAGCCATACCACATTTGTAGAGGTTTTACTTGCTTTAAAAAACCTCCCACACCTCCCCCTGAACCTGAAAC
ATAAAATGAATGCAATTGTTGTTGTTAACTTGTTTATTGCAGCTTATAATGGTTACAAATAAAGCAATAGCATCAC
AAATTTCACAAATAAAGCATTTTTTTCACTGCATTCTAGTTGTGGTTTGTCCAAACTCATCAATGTatcttaaTTAACC
ATTGTGGGAACCGTGCGATCAAACAAACGCGAGATACCGGAAGTACTGAAAAACAGTCGCTCCAGGCCAGTGG
GAACATCGATGTTTTGTTTTGACGGACCCCTTACTCTCGTCTCATATAAACCGAAGCCAGCTAAGATGGTATACTT
ATTATCATCTTGTGATGAGGATGCTTCTATCAACGAAAGTACCGGTAAACCGCAAATGGTTATGTATTATAATCAA
ACTAAAGGCGGAGTGGACACGCTAGACCAAATGTGTTCTGTGATGACCTGCAGTAGGAAGACGAATAGGTGGC
CTATGGCATTATTGTACGGAATGATAAACATTGCCTGCATAAATTCTTTTATTATATACAGCCATAATGTCAGTAG
CAAGGGAGAAAAGGTTCAAAGTCGCAAAAAATTTATGAGAAACCTTTACATGAGCCTGACGTCATCGTTTATGC
GTAAGCGTTTGGAAGCTCCTACTTTGAAGAGATATTTGCGCGATAATATCTCTAATATTTTGCCAAATGAAGTGCC
TGGTACATCAGATGACAGTACTGAAGAGCCAGTAATGAAAAAACGTACTTACTGTACTTACTGCCCCTCTAAAAT
AAGGCGAAAGGCAAATGCATCGTGCAAAAAATGCAAAAAAGTTATTTGTCGAGAGCATAATATTGATATGTGCC
AAAGTTGTTTCTGACTGACTAATAAGTATAATTTGTTTCTATTATGTATAAGTTAAGCTAATTACTTATTTTATAAT
ACAACATGACTGTTTTTAAAGTACAAAATAAGTTTATTTTTGTAAAAGAGAGAATGTTTAAAAGTTTTGTTACTTT
ATAGAAGAAATTTTGAGTTTTTGTTTTTTTTTAATAAATAAATAAACATAAATAAATTGTTTGTTGAATTTATTATT
AGTATGTAAGTGTAAATATAATAAAACTTAATATCTATTCAAATTAATAAATAAACCTCGATATACAGACCGATAA
AACACATGCGTCAATTTTACGCATGATTATCTTTAACGTACGTCACAATATGATTATCTTTCTAGGGTTAAATAATA
GTTTCTAATTTTTTTATTATTCAGCCTGCTGTCGTGAATACCGTATATCTCAACGCTGTCTGTGAGATTGTCGTATT
CTAGCCTTTTTAGTTTTTCGCTCATCGACTTGATATTGTCCGACACATTTTCGTCGATTTGCGTTTTGATCAAAGAC
TTGAGCAGAGACACGTTAATCAACTGTTCAAATTGATCCATATTAACGATATCAACCCGATGCGTATATGGTGCG
TAAAATATATTTTTTAACCCTCTTATACTTTGCACTCTGCGTTAATACGCGTTCGTGTACAGACGTAATCATGTTTT
CTTTTTTGGATAAAACTCCTACTGAGTTTGACCTCATATTAGACCCTCACAAGTTGCAAAACGTGGCATTTTTTACC
AATGAAGAATTTAAAGTTATTTTAAAAAATTTCATCACAGATTTAAAGAAGAACCAAAAATTAAATTATTTCAACA
GTTTAATCGACCAGTTAATCAACGTGTACACAGACGCGTCGGCAAAAAACACGCAGCCCGACGTGTTGGCTAAA
ATTATTAAATCAACTTGTGTTATAGTCACGGATTTGCCGTCCAACGTGTTCCTCAAAAAGTTGAAGACCAACAAGT
TTACGGACACTATTAATTATTTGATTTTGCCCCACTTCATTTTGTGGGATCACAATTTTGTTATATTTTTAAACAAA
GCTTGGCACTGGCCGTCGTTTTACAACGTCGTGACTGGGAAAACCCTGGCGTTACCCAACTTAATCGCCTTGCAG
CACATCCCCCTTTCGCCAGCTGGCGTAATAGCGAAGAGGCCCGCACCGATCGCCCTTCCCAACAGTTGCGCAGCC
TGAATGGCGAATGGCGCCTGATGCGGTATTTTCTCCTTACGCATCTGTGCGGTATTTCACACCGCATATGGTGCA
CTCTCAGTACAATCTGCTCTGATGCCGCATAGTTAAGCCAGCCCCGACACCCGCCAACACCCGCTGACGCGCCCT
GACGGGCTTGTCTGCTCCCGGCATCCGCTTACAGACAAGCTGTGACCGTCTCCGGGAGCTGCATGTGTCAGAGG
TTTTCACCGTCATCACCGAAACGCGCGAGACGAAAGGGCCTCGTGATACGCCTATTTTTATAGGTTAATGTCATG
ATAATAATGGTTTCTTAGACGTCAGGTGGCACTTTTCGGGGAAATGTGCGCGGAACCCCTATTTGTTTATTTTTCT
AAATACATTCAAATATGTATCCGCTCATGAGACAATAACCCTGATAAATGCTTCAATAATATTGAAAAAGGAAGA
GTATGAGTATTCAACATTTCCGTGTCGCCCTTATTCCCTTTTTTGCGGCATTTTGCCTTCCTGTTTTTGCTCACCCAG
AAACGCTGGTGAAAGTAAAAGATGCTGAAGATCAGTTGGGTGCACGAGTGGGTTACATCGAACTGGATCTCAAC
AGCGGTAAGATCCTTGAGAGTTTTCGCCCCGAAGAACGTTTTCCAATGATGAGCACTTTTAAAGTTCTGCTATGT
GGCGCGGTATTATCCCGTATTGACGCCGGGCAAGAGCAACTCGGTCGCCGCATACACTATTCTCAGAATGACTTG

GTTGAGTACTCACCAGTCACAGAAAAGCATCTTACGGATGGCATGACAGTAAGAGAATTATGCAGTGCTGCCAT
AACCATGAGTGATAACACTGCGGCCAACTTACTTCTGACAACGATCGGAGGACCGAAGGAGCTAACCGCTTTTT
TGCACAACATGGGGGATCATGTAACTCGCCTTGATCGTTGGGAACCGGAGCTGAATGAAGCCATACCAAACGAC
GAGCGTGACACCACGATGCCTGTAGCAATGGCAACAACGTTGCGCAAACTATTAACTGGCGAACTACTTACTCTA
GCTTCCCGGCAACAATTAATAGACTGGATGGAGGCGGATAAAGTTGCAGGACCACTTCTGCGCTCGGCCCTTCC
GGCTGGCTGGTTTATTGCTGATAAATCTGGAGCCGGTGAGCGTGGGTCTCGCGGTATCATTGCAGCACTGGGGC
CAGATGGTAAGCCCTCCCGTATCGTAGTTATCTACACGACGGGGAGTCAGGCAACTATGGATGAACGAAATAGA
CAGATCGCTGAGATAGGTGCCTCACTGATTAAGCATTGGTAACTGTCAGACCAAGTTTACTCATATATACTTTAGA
TTGATTTAAAACTTCATTTTTAATTTAAAAGGATCTAGGTGAAGATCCTTTTTGATAATCTCATGACCAAAATCCCT
TAACGTGAGTTTTCGTTCCACTGAGCGTCAGACCCCGTAGAAAAGATCAAAGGATCTTCTTGAGATCCTTTTTTTC
TGCGCGTAATCTGCTGCTTGCAAACAAAAAAACCACCGCTACCAGCGGTGGTTTGTTTGCCGGATCAAGAGCTAC
CAACTCTTTTTCCGAAGGTAACTGGCTTCAGCAGAGCGCAGATACCAAATACTGTTCTTCTAGTGTAGCCGTAGTT
AGGCCACCACTTCAAGAACTCTGTAGCACCGCCTACATACCTCGCTCTGCTAATCCTGTTACCAGTGGCTGCTGCC
AGTGGCGATAAGTCGTGTCTTACCGGGTTGGACTCAAGACGATAGTTACCGGATAAGGCGCAGCGGTCGGGCT
GAACGGGGGGTTCGTGCACACAGCCCAGCTTGGAGCGAACGACCTACACCGAACTGAGATACCTACAGCGTGA
GCTATGAGAAAGCGCCACGCTTCCCGAAGGGAGAAAGGCGGACAGGTATCCGGTAAGCGGCAGGGTCGGAAC
AGGAGAGCGCACGAGGGAGCTTCCAGGGGGAAACGCCTGGTATCTTTATAGTCCTGTCGGGTTTCGCCACCTCT
GACTTGAGCGTCGATTTTTGTGATGCTCGTCAGGGGGGCGGAGCCTATGGAAAAACGCCAGCAACGCGGCCTTT
TTACGGTTCCTGGCCTTTTGCTGGCCTTTTGCTCACATGTTCTTTCCTGCGTTATCCCCTGATTCTGTGGATAACCG
TATTACCGCCTTTGAGTGAGCTGATACCGCTCGCCGCAGCCGAACGACCGAGCGCAGCGAGTCAGTGAGCGAGG
AAGCGGAAGAGCGCCCAATACGCAAACCGCCTCTCCCCGCGCGTTGGCCGATTCATTAATGCAGCTGGCACGAC
AGG

**>188_Ae.pol.Nix**
TTTCCCGACTGGAAAGCGGGCAGTGAGCGCAACGCAATTAATGTGAGTTAGCTCACTCATTAGGCACCCCAGGC
TTTACACTTTATGCTTCCGGCTCGTATGTTGTGTGGAATTGTGAGCGGATAACAATTTCACACAGGAAACAGCTAT
GACATGATTACGAATTCGAGCTCGGTACCCGGGGATCCTCTAGAGTCGACGCTCGCGCGACTTGGTTTGCCATTC
TTTAGCGCGCGTCGCGTCACACAGCTTGGCCACAATGTGGTTTTTGTCAAACGAAGATTCTATGACGTGTTTAAA
GTTTAGGTCGAGTAAAGCGCAAATCTTTTTTAACCCTAGAAAGATAGTCTGCGTAAAATTGACGCATGCATTCTT
GAAATATTGCTCTCTCTTTCTAAATAGCGCGAATCCGTCGCTGTGCATTTAGGACATCTCAGTCGCCGCTTGGAGC
TCCCGTGAGGCGTGCTTGTCAATGCGGTAAGTGTCACTGATTTTGAACTATAACGACCGCGTGAGTCAAAATGAC
GCATGATTATCTTTTACGTGACTTTTAAGATTTAACTCATACGATAATTATATTGTTATTTCATGTTCTACTTACGTG
ATAACTTATTATATATATATATTTTCTTGTTATAGATATCGTGACTAATATATAATAAAATGGGTAGTTCTTTAGACGA
TGAGCATATCCTCTCTGCTCTTCTGCAAAGCGATGACGAGCTTGTTGGTGAGGATTCTGACAGTGAAATATCAGA
TCACGTAAGTGAAGATGACGTCCAGAGCGATACAGAAGAAGCGTTTATAGATGAGGTACATGAAGTGCAGCCA
ACGTCAAGCGGTAGTGAAATATTAGACGAACAAAATGTTATTGAACAACCAGGTTCTTCATTGGCTTCTAACAGA
ATCTTGACCTTGCCACAGAGGACTATTAGAGGTAAGAATAAACATTGTTGGTCAACTTCAAAGTCCACGAGGCGT
AGCCGAGTCTCTGCACTGAACATTGTCAGATCTaacccc ttgtgtcatgtcggcgaccctacgcccccaactgagagaactcaaaggt
taccccagttggggcactactcccgaaaaccgcttctgacctgggTAAGATACATTGATGAGTTTGGACAAACCACAACTAGAAT
GCAGTGAAAAAAATGCTTTATTTGTGAAATTTGTGATGCTATTGCTTTATTTGTAACCATTATAAGCTGCAATAAA
CAAGTTAACAACAACAATTGCATTCATTTTATGTTTCAGGTTCAGGGGGAGGTGTGGGAGGTTTTTTAAAGCAAG
TAAAACCTCTACAAATGTGGTATGGCTGATTATGATCTAGCGGCCGCTTTACTTGTACAGCTCGTCCATGCCGAG
AGTGATCCCGGCGGCGGTCACGAACTCCAGCAGGACCATGTGATCGCGCTTCTCGTTGGGGTCTTTGCTCAGGG
CGGACTGGGTGCTCAGGTAGTGGTTGTCGGGCAGCAGCACGGGGCCGTCGCCGATGGGGGTGTTCTGCTGGTA

GTGGTCGGCGAGCTGCACGCTGCCGTCCTCGATGTTGTGGCGGATCTTGAAGTTCACCTTGATGCCGTTCTTCTG
CTTGTCGGCCATGATATAGACGTTGTGGCTGTTGTAGTTGTACTCCAGCTTGTGCCCCAGGATGTTGCCGTCCTCC
TTGAAGTCGATGCCCTTCAGCTCGATGCGGTTCACCAGGGTGTCGCCCTCGAACTTCACCTCGGCGCGGGTCTTG
TAGTTGCCGTCGTCCTTGAAGAAGATGGTGCGCTCCTGGACGTAGCCTTCGGGCATGGCGGACTTGAAGAAGTC
GTGCTGCTTCATGTGGTCGGGGTAGCGGCTGAAGCACTGCACGCCGTAGGTCAGGGTGGTCACGAGGGTGGGC
CAGGGCACGGGCAGCTTGCCGGTGGTGCAGATGAACTTCAGGGTCAGCTTGCCGTAGGTGGCATCGCCCTCGCC
CTCGCCGGACACGCTGAACTTGTGGCCGTTTACGTCGCCGTCCAGCTCGACCAGGATGGGCACCACCCCGGTGA
ACAGCTCCTCGCCCTTGCTCACCATGGTTGAAATCTCTGTTGAGCAGAAAAAGAAACGAGGAAACGCTTgAGTAA
TTGGTTGTGAAATGCAAACTCTCATTTGATATTGATTCATTGCCTTTGGCTTCGAGCACGACACGACAGGTTTTAA
ACTTGTTTTGCTTTGTCTGCGTTTGCAGTCGCAGGCCAAGTGAAAAATATACACTTGAAGGTGATGACGTCACAA
CAACGCCCTACTTTTaAGTGAAAAATTAACTTGTTTTCGACTTTGAACTACGTAGTTTtGAAATTGCGTATCTTCAA
GTTTTCACGATTTCCTCAAGGTTTTTCTCGATATGTGTTAATATTACCTTAATGGGTAATTACCATCAAAATATTTA
TTTTAGATATGTGACGGAGCAAATACGTTATTCTTATTATTCTAGAAATTTAATTCAATTAGtAGCGATGATTCAAC
GAAATATGATTATCGCtGTGAATCACAATTGGGTTTTATCAATGATGATGAAACTGCGTTGCAAATTTTCACTAAT
CACTCAAAGCTCAATAGTCGCCATCTTGGAAAATAGTTTGCTCATTCAAAGACAAAGGAATCATTCACCAAACTA
GTTTTCGCTCATAGCTATAATTTCATCAaTTAATTTACCTACCTTCACTAGAAGATTCCCTTCACCGAAATGCACTTT
CACcAATTTAATAGTAATTGTCCTTTGAAtAAAGCTTTGTTCACTCTGAAATTTTCTCCTCTGGCTAATTGGATCACT
CTTTTTCACTAGAGACTTCACTTCACTTGCACTGGCACTGCTTACTTGGgCCGCGTAATGTTCACTCCACTAGGAAA
CGTATTCGATTGAGCTGGTTTcGCCTTTGCAGGGGCGTTTTATAGACACTGtCGTAGTGGTGGTTGTACTTCTAGA
AAATTTCaGCAATaCATTCATACATATACCTCTGTTCGGTTGGATGGCTCTAtATCGATCGAaTATGGGTACCATCC
CTGTGTCTGAATGGACAGCAAAACGTGCTTGTGTCTGTTAGTCGTTCATTCTGTACCTTGAACGATGCAGTTCAAC
TTCTGGCAAAGACGTCAATGTACCTACCcTTCGTGTATATGGCATAgAGAGgAAGATGTGCGAATGCCTTTTTCGA
TAGAGAAAGGATTTCCTGATTTGATCGACAATTTCGGTAGCTAaTCTTAGCTTCGAATGTAATCTGATAACCAAAT
CCAGAGAaAAAtCAGTAATTTGGGAATTTCACTTGAATCATTCTAATTGcATCATTTGCTGTATTAGTGCAGTCAGC
AAGTGACGTCAACCCTTCTAAATCGATATACTTCTGGGAAGCTTTCTTTCTTGTCTGGCTCAGCTGGTGCCAAGGC
AAATTATAATTGGATTCAATGCACAAGCTACATGTAAAGATActcgagAGAACAAAAACCGCTCCAGAAACTACTT
ACCTTGAAATGATATTTCAAATATTTTTGCTAGAAGGGTGTAGATCCGAATCCATACCCATATACGAATAGTTTCA
GAGTCGAAAAAGAATATTAAACATTCAAAACTAAATTTAGTTAACAAAAAAACGCCACCGCAAAGTCAACTTGA
AAATGGCGATGCTGCTCGTCAAAGTATCGTGCGTATTATGTACCCATGATTAGTAAACGAATGGAAACTTATTTA
TGTGGATTAAGTACCCTTTTTATGACATTTGAAAAATACCCTCTTTCTGACAATTCAGTACTGGTTGGGAAAATGG
GTACATGGAACCCATTTAATGGGTACTTCCAAGTTAGCGTGAAGTTTACTGTCAATGTCATTCCAATCGAGCAGG
AGACTTGTCAAGCACTTTTTCACACAAGCTGAAAACGGCTCACACAAAAATGTTACCGCCCACGAAAATTTTGCTT
CTTCTGAAGTAGCATATTTTCTGAATACTTGAGTGTCGGTTGCAATTTTCATGCTTTGCATAAAGAATGCTCTACTC
ATTACATTGGTTTTCCACACATTCAGTCAATTTTTCATTCGTTACCCAAATCTGTGAAAATTCAACACATAAAACTT
ATGACTCACTTTTAGCGTGGGATTCAACATTGGCAGTGCTGCGTAATAGATTTGGCACAGTTGCTAGCTGATTTA
TTTTGAATATGATCAGGTTTTAGGTAACGCTCGTAACGCTTTTTGTATGAATATTCTACAAATTTTGTATGAGCCGT
AACACCAGCAAAACACCCACCCACCCCCTTCAGCGTTATGAAATTTGTAAATAAGCCCATATGGGTAAAATATAC
CAATAAAAAATTGGTAAATTTCACCCACATTATTAGTTTACTGGATTTACCCAAATATGGGTAGATGCGGTTACCCA
TATTTGGATGAAAAATTAGAATTGTTGTCAGTCAGAGTGCGGCGCTCAAGATGGAAATTCATTTTTCAGCAGTAC
TGAATAAATTGTTTTAGGATCAAGAAGAAAAATGATGATTCCTATCGGAGGTGAGTAGGTATAGTCATTTATTCA
TTGTTATTAGGTGTTATATATTCCAGTTTCTTCATCGATGTGCACCAATGTTTTTCTTCATGGCGACAAAAAACTGA
ACATTTCGGGCTGGTTAAATTGTGCTGTACTAACCAAGATGCAACAACTACAAAACTGAATCGAAATTGGGAGAC
AGCAATTCCAAATAAGAATGTCGTTCCGTTTATTTGTCACGATATACATATTTCACAACTCTTGGTTGACAATAATT
TTACTCTATTTTATAATATATCTTTATTTCTCAATAAAGTTTGCATTTGTATGTTAATGACAAAATTCAATTTAATTC

AAAATAGTCATAAAGAAATTTAATAAAAAAGCTAGTCGAAAAGTCTATGAAACTACTCATTTTTGCGTATTTTTTA
GAATAATTGCTTCCAAGCTTGAATACCCAAATATGGGTATTGTCAGTTTACCTATTTATGAGTAAACCCGCTTTTT
GGCGATTATGGGCAAACTTTACCCATATTTGGGTAGACTGTTCTTAGCGTGTACATCTAGGGCACCCATACCCAA
TGTGATCTATAAGAGGTGAAGAAAAAGTCATTGTGTGATTGTATTAGTACAATAAAAATCAAAACAAACATTGAG
GGGAAAATGAGTGACGTCATACCGTTTCAACGAAATGCTGTGAGTGGATGTAAAACACTCCATATCGTGGAACA
AATCAAGATCACACCGTATAAATCTGCAACTAACACCTAGATGAAAGAGTCGAATGGGTAGATTCCAGGAATTAT
GTTTTTCACTATATTCACGGATGAAAGAAAGCACTTTTCTTACTTTTTTCTAATTATATTCAATTTGAGCACTACAT
TTATGGATGAATTACGCAAAATATCCTTTGGAAGAAAGAACAATTTCAGCCTATTCCAGCACATTATCAGTTTTGA
TGATATTTGGGTTGCCTTCCGTTGTTATATTTCCTCTGTTTTATAATAGATTTCATCCTAAAATAAAGCAAAACTTA
GCATGAAGCAAACAACGCATGAATTACCTGAAATGCCATCTTTCTGCGACGAATTTTTTCATGTTGTTGTTTGAAC
AACAGCACACAGGACAACCTTGGTAACGATAGGGTTGCTAGTAACGATACCTCGAAATAAATCGATAACAAACT
GTATGACGTCACGCAGTGGTGGAATGCAATGGGTTGCTGTGTGAAAATTTAAGCGACCCACTAACACATGCAAG
CAGTTGTGTATTACACAAAATCATCTCCTCACCTCTAATAAAGCACATTGACCCATACCATTGGGCGTGATAACCA
CTATAATCGAGTAAAAATAAGAGAGAACAGTGGCACGCAACATGATTTTTCCCATTTTTCCACCTGAGCAAACTA
AGAACATTGACTACGGAGCTCAGTCCAGCAATAATGTTAATCATAAAAAGATGTGATCTTTTTCAAAGAAAATTT
TATTAGTGTTTTCTAAGTTTTATATTGATTTTTGTTTTTGTCGTGCAAATGTACAGCAAAAATGAGATCAATCTCAT
TAATAAGCAATTTGAATATATTAAAAAATATTGCATATATATTGGAAACATTCCTGCCGAAGTGTCAAAAAGAGA
TTTAGTTGCAAAATTTTCCGAATTCGGGGAAATATCCAACTTATATTTGAAGTCATTCGTAAAGTTCTGTGATGTG
AAACATGCTGTAATTCGTTACAGATTGGAAACAAGTGTAAAGGAATCTTCAAGTTTACACAATAATCGATATATT
CAATCGGTTTTAATAGTTCTGCCACTAGACTCGTCGTACACCCATTACTATCTTCCTTACAACACTTGTGTTGCGGT
GTACACTAACAAACATTTTCATATGGTAGAACTTTTTGAAAATTTTAAAAGATTTGGAGACATTCAAGTCATAAAG
AAAACTACAAACGTCATGGGATACATTTGTTTTACAACAGAAAATGCTGCAAGAAAAAATGTTGGTTACTAAGCCC
ACAGATATCCATAAAAATGTACAAAAAATTAATGATGTTACGCGAAACATTAACGTTTGCTTAATAGATTTCGATA
ACGAATCTTCTGGAAATACGGCGATAAAACTAACACTTTTATATAATCGCTCGATAGGAGTATTCGGGCTGCCCT
CAGATTTCACAGAAGAAAAACTGTACGATGAATTTTCAAGGTATGGCAGAATAGAAAAAAATAAACTAGTGTAC
GACTCTACCGGACACTCTAAACAATACGGTTTTGTTTATTATGAAAAGCACATCTCTGCTCAAGCTGCCAAAGAG
GAAATGGACCGCAGTGATTATACAGGAGGTAAAATTTCCGTCCGTTTTGTTCCAGAAAAAGAGTAATAAAGTGA
TATATGACGCGGCCGCGACTCTAGATCATAATCAGCCATACCACATTTGTAGAGGTTTTACTTGCTTTAAAAAACC
TCCCACACCTCCCCCTGAACCTGAAACATAAAATGAATGCAATTGTTGTTGTTAACTTGTTTATTGCAGCTTATAAT
GGTTACAAATAAAGCAATAGCATCACAAATTTCACAAATAAAGCATTTTTTTCACTGCATTCTAGTTGTGGTTTGT
CCAAACTCATCAATGTatcttaaTTAACCATTGTGGGAACCGTGCGATCAAACAAACGCGAGATACCGGAAGTACT
GAAAAACAGTCGCTCCAGGCCAGTGGGAACATCGATGTTTTGTTTTGACGGACCCCTTACTCTCGTCTCATATAA
ACCGAAGCCAGCTAAGATGGTATACTTATTATCATCTTGTGATGAGGATGCTTCTATCAACGAAAGTACCGGTAA
ACCGCAAATGGTTATGTATTATAATCAAACTAAAGGCGGAGTGGACACGCTAGACCAAATGTGTTCTGTGATGA
CCTGCAGTAGGAAGACGAATAGGTGGCCTATGGCATTATTGTACGGAATGATAAACATTGCCTGCATAAATTCTT
TTATTATATACAGCCATAATGTCAGTAGCAAGGGAGAAAAGGTTCAAAGTCGCAAAAAATTTATGAGAAACCTTT
ACATGAGCCTGACGTCATCGTTTATGCGTAAGCGTTTGGAAGCTCCTACTTTGAAGAGATATTTGCGCGATAATA
TCTCTAATATTTTGCCAAATGAAGTGCCTGGTACATCAGATGACAGTACTGAAGAGCCAGTAATGAAAAAACGTA
CTTACTGTACTTACTGCCCCTCTAAAATAAGGCGAAAGGCAAATGCATCGTGCAAAAAATGCAAAAAAGTTATTT
GTCGAGAGCATAATATTGATATGTGCCAAAGTTGTTTCTGACTGACTAATAAGTATAATTTGTTTCTATTATGTAT
AAGTTAAGCTAATTACTTATTTTATAATACAACATGACTGTTTTTAAAGTACAAAATAAGTTTATTTTTGTAAAAGA
GAGAATGTTTAAAAGTTTTGTTACTTTATAGAAGAAATTTTGAGTTTTTGTTTTTTTTTAATAAATAAATAAACATA
AATAAATTGTTTGTTGAATTTATTATTAGTATGTAAGTGTAAATATAATAAAACTTAATATCTATTCAAATTAATAA
ATAAACCTCGATATACAGACCGATAAAACACATGCGTCAATTTTACGCATGATTATCTTTAACGTACGTCACAATA

TGATTATCTTTCTAGGGTTAAATAATAGTTTCTAATTTTTTTATTATTCAGCCTGCTGTCGTGAATACCGTATATCTC
AACGCTGTCTGTGAGATTGTCGTATTCTAGCCTTTTTAGTTTTTCGCTCATCGACTTGATATTGTCCGACACATTTT
CGTCGATTTGCGTTTTGATCAAAGACTTGAGCAGAGACACGTTAATCAACTGTTCAAATTGATCCATATTAACGAT
ATCAACCCGATGCGTATATGGTGCGTAAAATATATTTTTTAACCCTCTTATACTTTGCACTCTGCGTTAATACGCGT
TCGTGTACAGACGTAATCATGTTTTCTTTTTTGGATAAAACTCCTACTGAGTTTGACCTCATATTAGACCCTCACAA
GTTGCAAAACGTGGCATTTTTTACCAATGAAGAATTTAAAGTTATTTTAAAAAAATTTCATCACAGATTTAAAGAAG
AACCAAAAATTAAATTATTTCAACAGTTTAATCGACCAGTTAATCAACGTGTACACAGACGCGTCGGCAAAAAAC
ACGCAGCCCGACGTGTTGGCTAAAATTATTAAATCAACTTGTGTTATAGTCACGGATTTGCCGTCCAACGTGTTCC
TCAAAAAGTTGAAGACCAACAAGTTTACGGACACTATTAATTATTTGATTTTGCCCCACTTCATTTTGTGGGATCA
CAATTTTGTTATATTTTTAAACAAAGCTTGGCACTGGCCGTCGTTTTACAACGTCGTGACTGGGAAAACCCTGGCG
TTACCCAACTTAATCGCCTTGCAGCACATCCCCCTTTCGCCAGCTGGCGTAATAGCGAAGAGGCCCGCACCGATC
GCCCTTCCCAACAGTTGCGCAGCCTGAATGGCGAATGGCGCCTGATGCGGTATTTTCTCCTTACGCATCTGTGCG
GTATTTCACACCGCATATGGTGCACTCTCAGTACAATCTGCTCTGATGCCGCATAGTTAAGCCAGCCCCGACACCC
GCCAACACCCGCTGACGCGCCCTGACGGGCTTGTCTGCTCCCGGCATCCGCTTACAGACAAGCTGTGACCGTCTC
CGGGAGCTGCATGTGTCAGAGGTTTTCACCGTCATCACCGAAACGCGCGAGACGAAAGGGCCTCGTGATACGCC
TATTTTTATAGGTTAATGTCATGATAATAATGGTTTCTTAGACGTCAGGTGGCACTTTTCGGGGAAATGTGCGCG
GAACCCCTATTTGTTTATTTTTCTAAATACATTCAAATATGTATCCGCTCATGAGACAATAACCCTGATAAATGCTT
CAATAATATTGAAAAAGGAAGAGTATGAGTATTCAACATTTCCGTGTCGCCCTTATTCCCTTTTTTGCGGCATTTT
GCCTTCCTGTTTTTGCTCACCCAGAAACGCTGGTGAAAGTAAAAGATGCTGAAGATCAGTTGGGTGCACGAGTG
GGTTACATCGAACTGGATCTCAACAGCGGTAAGATCCTTGAGAGTTTTCGCCCCGAAGAACGTTTTCCAATGATG
AGCACTTTTAAAGTTCTGCTATGTGGCGCGGTATTATCCCGTATTGACGCCGGGCAAGAGCAACTCGGTCGCCGC
ATACACTATTCTCAGAATGACTTGGTTGAGTACTCACCAGTCACAGAAAAGCATCTTACGGATGGCATGACAGTA
AGAGAATTATGCAGTGCTGCCATAACCATGAGTGATAACACTGCGGCCAACTTACTTCTGACAACGATCGGAGG
ACCGAAGGAGCTAACCGCTTTTTTGCACAACATGGGGGATCATGTAACTCGCCTTGATCGTTGGGAACCGGAG
CTGAATGAAGCCATACCAAACGACGAGCGTGACACCACGATGCCTGTAGCAATGGCAACAACGTTGCGCAAACT
ATTAACTGGCGAACTACTTACTCTAGCTTCCCGGCAACAATTAATAGACTGGATGGAGGCGGATAAAGTTGCAG
GACCACTTCTGCGCTCGGCCCTTCCGGCTGGCTGGTTTATTGCTGATAAATCTGGAGCCGGTGAGCGTGGGTCTC
GCGGTATCATTGCAGCACTGGGGCCAGATGGTAAGCCCTCCCGTATCGTAGTTATCTACACGACGGGGAGTCAG
GCAACTATGGATGAACGAAATAGACAGATCGCTGAGATAGGTGCCTCACTGATTAAGCATTGGTAACTGTCAGA
CCAAGTTTACTCATATATACTTTAGATTGATTTAAAACTTCATTTTTAATTTAAAAGGATCTAGGTGAAGATCCTTT
TTGATAATCTCATGACCAAAATCCCTTAACGTGAGTTTTCGTTCCACTGAGCGTCAGACCCCGTAGAAAAGATCAA
AGGATCTTCTTGAGATCCTTTTTTTCTGCGCGTAATCTGCTGCTTGCAAACAAAAAAACCACCGCTACCAGCGGTG
GTTTGTTTGCCGGATCAAGAGCTACCAACTCTTTTTCCGAAGGTAACTGGCTTCAGCAGAGCGCAGATACCAAAT
ACTGTTCTTCTAGTGTAGCCGTAGTTAGGCCACCACTTCAAGAACTCTGTAGCACCGCCTACATACCTCGCTCTGC
TAATCCTGTTACCAGTGGCTGCTGCCAGTGGCGATAAGTCGTGTCTTACCGGGTTGGACTCAAGACGATAGTTAC
CGGATAAGGCGCAGCGGTCGGGCTGAACGGGGGGTTCGTGCACACAGCCCAGCTTGGAGCGAACGACCTACAC
CGAACTGAGATACCTACAGCGTGAGCTATGAGAAAGCGCCACGCTTCCCGAAGGGAGAAAGGCGGACAGGTAT
CCGGTAAGCGGCAGGGTCGGAACAGGAGAGCGCACGAGGGAGCTTCCAGGGGGAAACGCCTGGTATCTTTATA
GTCCTGTCGGGTTTCGCCACCTCTGACTTGAGCGTCGATTTTTGTGATGCTCGTCAGGGGGGCGGAGCCTATGGA
AAAACGCCAGCAACGCGGCCTTTTTACGGTTCCTGGCCTTTTGCTGGCCTTTTGCTCACATGTTCTTTCCTGCGTTA
TCCCCTGATTCTGTGGATAACCGTATTACCGCCTTTGAGTGAGCTGATACCGCTCGCCGCAGCCGAACGACCGAG
CGCAGCGAGTCAGTGAGCGAGGAAGCGGAAGAGCGCCCAATACGCAAACCGCCTCTCCCCGCGCGTTGGCCGA
TTCATTAATGCAGCTGGCACGACAGG

**>189_Ae.vex.Nix**
TTTCCCGACTGGAAAGCGGGCAGTGAGCGCAACGCAATTAATGTGAGTTAGCTCACTCATTAGGCACCCCAGGC
TTTACACTTTATGCTTCCGGCTCGTATGTTGTGTGGAATTGTGAGCGGATAACAATTTCACACAGGAAACAGCTAT
GACATGATTACGAATTCGAGCTCGGTACCCGGGGATCCTCTAGAGTCGACGCTCGCGCGACTTGGTTTGCCATTC
TTTAGCGCGCGTCGCGTCACACAGCTTGGCCACAATGTGGTTTTTGTCAAACGAAGATTCTATGACGTGTTTAAA
GTTTAGGTCGAGTAAAGCGCAAATCTTTTTTAACCCTAGAAAGATAGTCTGCGTAAAATTGACGCATGCATTCTT
GAAATATTGCTCTCTCTTTCTAAATAGCGCGAATCCGTCGCTGTGCATTTAGGACATCTCAGTCGCCGCTTGGAGC
TCCCGTGAGGCGTGCTTGTCAATGCGGTAAGTGTCACTGATTTTGAACTATAACGACCGCGTGAGTCAAAATGAC
GCATGATTATCTTTTACGTGACTTTTAAGATTTAACTCATACGATAATTATATTGTTATTTCATGTTCTACTTACGTG
ATAACTTATTATATATATATATTTTCTTGTTATAGATATCGTGACTAATATATAATAAAATGGGTAGTTCTTTAGACGA
TGAGCATATCCTCTCTGCTCTTCTGCAAAGCGATGACGAGCTTGTTGGTGAGGATTCTGACAGTGAAATATCAGA
TCACGTAAGTGAAGATGACGTCCAGAGCGATACAGAAGAAGCGTTTATAGATGAGGTACATGAAGTGCAGCCA
ACGTCAAGCGGTAGTGAAATATTAGACGAACAAAATGTTATTGAACAACCAGGTTCTTCATTGGCTTCTAACAGA
ATCTTGACCTTGCCACAGAGGACTATTAGAGGTAAGAATAAACATTGTTGGTCAACTTCAAAGTCCACGAGGCGT
AGCCGAGTCTCTGCACTGAACATTGTCAGATCTaacccccttgtgtcatgtcggcgaccctacgcccccaactgagagaactcaaaggt
taccccagttggggcactactcccgaaaaccgcttctgacctgggTAAGATACATTGATGAGTTTGGACAAACCACAACTAGAAT
GCAGTGAAAAAAATGCTTTATTTGTGAAATTTGTGATGCTATTGCTTTATTTGTAACCATTATAAGCTGCAATAAA
CAAGTTAACAACAACAATTGCATTCATTTTATGTTTCAGGTTCAGGGGGAGGTGTGGGAGGTTTTTTAAAGCAAG
TAAAACCTCTACAAATGTGGTATGGCTGATTATGATCTAGCGGCCGCTTTACTTGTACAGCTCGTCCATGCCGAG
AGTGATCCCGGCGGCGGTCACGAACTCCAGCAGGACCATGTGATCGCGCTTCTCGTTGGGGTCTTTGCTCAGGG
CGGACTGGGTGCTCAGGTAGTGGTTGTCGGGCAGCAGCACGGGGCCGTCGCCGATGGGGGTGTTCTGCTGGTA
GTGGTCGGCGAGCTGCACGCTGCCGTCCTCGATGTTGTGGCGGATCTTGAAGTTCACCTTGATGCCGTTCTTCTG
CTTGTCGGCCATGATATAGACGTTGTGGCTGTTGTAGTTGTACTCCAGCTTGTGCCCCAGGATGTTGCCGTCCTCC
TTGAAGTCGATGCCCTTCAGCTCGATGCGGTTCACCAGGGTGTCGCCCTCGAACTTCACCTCGGCGCGGGTCTTG
TAGTTGCCGTCGTCCTTGAAGAAGATGGTGCGCTCCTGGACGTAGCCTTCGGGCATGGCGGACTTGAAGAAGTC
GTGCTGCTTCATGTGGTCGGGGTAGCGGCTGAAGCACTGCACGCCGTAGGTCAGGGTGGTCACGAGGGTGGGC
CAGGGCACGGGCAGCTTGCCGGTGGTGCAGATGAACTTCAGGGTCAGCTTGCCGTAGGTGGCATCGCCCTCGCC
CTCGCCGGACACGCTGAACTTGTGGCCGTTTACGTCGCCGTCCAGCTCGACCAGGATGGGCACCACCCCGGTGA
ACAGCTCCTCGCCCTTGCTCACCATGGTTGAAATCTCTGTTGAGCAGAAAAAGAAACGAGGAAACGCTTgAGTAA
TTGGTTGTGAAATGCAAACTCTCATTTGATATTGATTCATTGCCTTTGGCTTCGAGCACGACACGACAGGTTTTAA
ACTTGTTTTGCTTTGTCTGCGTTTGCAGTCGCAGGCCAAGTGAAAAATATACACTTGAAGGTGATGACGTCACAA
CAACGCCCTACTTTTaAGTGAAAAATTAACTTGTTTTCGACTTTGAACTACGTAGTTTtGAAATTGCGTATCTTCAA
GTTTTCACGATTTCCTCAAGGTTTTTCTCGATATGTGTTAATATTACCTTAATGGGTAATTACCATCAAAATATTTA
TTTTAGATATGTGACGGAGCAAATACGTTATTCTTATTATTCTAGAAATTTAATTCAATTAGtAGCGATGATTCAAC
GAAATATGATTATCGCtGTGAATCACAATTGGGTTTTATCAATGATGATGAAACTGCGTTGCAAATTTTCACTAAT
CACTCAAAGCTCAATAGTCGCCATCTTGGAAAATAGTTTGCTCATTCAAAGACAAAGGAATCATTCACCAAACTA
GTTTTCGCTCATAGCTATAATTTCATCAaTTAATTTACCTACCTTCACTAGAAGATTCCCTTCACCGAAATGCACTTT
CACcAATTTAATAGTAATTGTCCTTTGAAtAAAGCTTTGTTCACTCTGAAATTTTCTCCTCTGGCTAATTGGATCACT
CTTTTTCACTAGAGACTTCACTTCACTTGCACTGGCACTGCTTACTTGGgCCGCGTAATGTTCACTCCACTAGGAAA
CGTATTCGATTGAGCTGGTTTcGCCTTTGCAGGGGGCGTTTTATAGACACTGtCGTAGTGGTGGTTGTACTTCTAGA
AAATTTCaGCAATaCATTCATACATATACCTCTGTTCGGTTGGATGGCTCTAtATCGATCGAaTATGGGTACCATCC
CTGTGTCTGAATGGACAGCAAAACGTGCTTGTGTCTGTTAGTCGTTCATTCTGTACCTTGAACGATGCAGTTCAAC
TTCTGGCAAAGACGTCAATGTACCTACCcTTCGTGTATATGGCATAgAGAGgAAGATGTGCGAATGCCTTTTTCGA
TAGAGAAAGGATTTCCTGATTTGATCGACAATTTCGGTAGCTAaTCTTAGCTTCGAATGTAATCTGATAACCAAAT

CCAGAGAaAAAtCAGTAATTTGGGAATTTCACTTGAATCATTCTAATTGcATCATTTGCTGTATTAGTGCAGTCAGC
AAGTGACGTCAACCCTTCTAAATCGATATACTTCTGGGAAGCTTTCTTTCTTGTCTGGCTCAGCTGGTGCCAAGGC
AAATTATAATTGGATTCAATGCACAAGCTACATGTAAAGATActcgagAGAACAAAAACCGCTCCAGAAACTACTT
ACCTTGAAATGATATTTCAAATATTTTTGCTAGAAGGGTGTAGATCCGAATCCATACCCATATACGAATAGTTTCA
GAGTCGAAAAAGAATATTAAACATTCAAAACTAAATTTTAGTTAACAAAAAAACGCCACCGCAAAGTCAACTTGA
AAATGGCGATGCTGCTCGTCAAAGTATCGTGCGTATTATGTACCCATGATTAGTAAACGAATGGAAACTTATTTA
TGTGGATTAAGTACCCTTTTTATGACATTTGAAAAATACCCTCTTTCTGACAATTCAGTACTGGTTGGGAAAATGG
GTACATGGAACCCATTTAATGGGTACTTCCAAGTTAGCGTGAAGTTTACTGTCAATGTCATTCCAATCGAGCAGG
AGACTTGTCAAGCACTTTTTCACACAAGCTGAAAACGGCTCACACAAAAATGTTACCGCCCACGAAAATTTTGCTT
CTTCTGAAGTAGCATATTTTCTGAATACTTGAGTGTCGGTTGCAATTTTCATGCTTTGCATAAAGAATGCTCTACTC
ATTACATTGGTTTTCCACACATTCAGTCAATTTTTCATTCGTTACCCAAATCTGTGAAAATTCAACACATAAAACTT
ATGACTCACTTTTAGCGTGGGATTCAACATTGGCAGTGCTGCGTAATAGATTTGGCACAGTTGCTAGCTGATTTA
TTTTGAATATGATCAGGTTTTAGGTAACGCTCGTAACGCTTTTTGTATGAATATTCTACAAATTTTGTATGAGCCGT
AACACCAGCAAAACACCCACCCACCCCCTTCAGCGTTATGAAATTTGTAAATAAGCCCATATGGGTAAAATATAC
CAATAAAAATTGGTAAATTTCACCCACATTATTAGTTTACTGGATTTACCCAAATATGGGTAGATGCGGTTACCCA
TATTTGGATGAAAAATTAGAATTGTTGTCAGTCAGAGTGCGGCGCTCAAGATGGAAATTCATTTTTCAGCAGTAC
TGAATAAATTGTTTTAGGATCAAGAAGAAAAATGATGATTCCTATCGGAGGTGAGTAGGTATAGTCATTTATTCA
TTGTTATTAGGTGTTATATATTCCAGTTTCTTCATCGATGTGCACCAATGTTTTTCTTCATGGCGACAAAAAACTGA
ACATTTCGGGCTGGTTAAATTGTGCTGTACTAACCAAGATGCAACAACTACAAAACTGAATCGAAATTGGGAGAC
AGCAATTCCAAATAAGAATGTCGTTCCGTTTATTTGTCACGATATACATATTTCACAACTCTTGGTTGACAATAATT
TTACTCTATTTTATAATATATCTTTATTTCTCAATAAAGTTTGCATTTGTATGTTAATGACAAAATTCAATTTAATTC
AAAATAGTCATAAAGAAATTTAATAAAAAAGCTAGTCGAAAAGTCTATGAAACTACTCATTTTTGCGTATTTTTTA
GAATAATTGCTTCCAAGCTTGAATACCCAAATATGGGTATTGTCAGTTTACCTATTTATGAGTAAACCCGCTTTTT
GGCGATTATGGGCAAACTTTACCCATATTTGGGTAGACTGTTCTTAGCGTGTACATCTAGGGCACCCATACCCAA
TGTGATCTATAAGAGGTGAAGAAAAAGTCATTGTGTGATTGTATTAGTACAATAAAAATCAAAACAAACATTGAG
GGGAAAATGAGTGACGTCATACCGTTTCAACGAAATGCTGTGAGTGGATGTAAAACACTCCATATCGTGGAACA
AATCAAGATCACACCGTATAAATCTGCAACTAACACCTAGATGAAAGAGTCGAATGGGTAGATTCCAGGAATTAT
GTTTTTTCACTATATTCACGGATGAAAGAAAGCACTTTTCTTACTTTTTTCTAATTATATTCAATTTGAGCACTACAT
TTATGGATGAATTACGCAAAATATCCTTTGGAAGAAAGAACAATTTCAGCCTATTCCAGCACATTATCAGTTTTGA
TGATATTTGGGTTGCCTTCCGTTGTTATATTTCCTCTGTTTTATAATAGATTTCATCCTAAAATAAAGCAAAACTTA
GCATGAAGCAAACAACGCATGAATTACCTGAAATGCCATCTTTCTGCGACGAATTTTTTCATGTTGTTGTTTGAAC
AACAGCACACAGGACAACCTTGGTAACGATAGGGTTGCTAGTAACGATACCTCGAAATAAATCGATAACAAACT
GTATGACGTCACGCAGTGGTGGAATGCAATGGGTTGCTGTGTGAAAATTTAAGCGACCCACTAACACATGCAAG
CAGTTGTGTATTACACAAAATCATCTCCTCACCTCTAATAAAGCACATTGACCCATACCATTGGGCGTGATAACCA
CTATAATCGAGTAAAAATAAGAGAGAACAGTGGCACGCAACATGATTTTTCCCATTTTTCCACCTGAGCAAACTA
AGAACATTGACTACGGAGCTCAGTCCAGCAATAATGTTAATCATAAAAAGATGTGATCTTTTTCAAAGAAAATTT
TATTAGTGTTTTCTAAGTTTTATATTGATTTTTGTTTTTGTCGTGCAAATGTTTACTCCTTTACATTATGCCGCAAGT
CATCGCCACCTAACGAGTGAAGTAATCAACACAGAATTTGAAATTATTGCTCAGAATTGTATTTACATTGGAAAC
ATTCCAAATTTTGTTTCAAAAAAAAATATTGTTTGATTTGTTTACTCAATCTGGCGGTGAAATATTTAACATTTATAT
ACAGCAAAATGAACACTGCGATGTAAAAGCAGCAATCATTAGATTTCTACATAAAAAAAGTGTAATGCGATCCTT
AAGTTTGAACAAAACTCGATATCATCAGTCAATACTAATAGTAATAGAACTAAGTTTGCCATATGCAAACTATCTA
CTAGTATATAATACATTGATAGTGATTTATATCCGAGAGAAAACAAAAAAATATTCACTGGAGCAAATTTACGAT
GAGTTTAAAAGTTTTGGAGCAATTCGAAATATTTTGAAAACAACGAATTTAATGGTTTATATTAATTATTACTCAA
AAAAAGCCCAAGAATCAGCCCTTAAATCAAAGTCACTTCCTGATTATGTTGATAAAATAATTACATCTCATAGAAA

TATACACATGTGTTGCATAAATTTAGACAGTAAGGATTTCTCGAGAAATTTAGAAATCAAAATTAAACTTTTATAT
AGCAGTACCATCGGCATATTTGGATTGCCATCTGATACCAAGGAAGCAGAAATTACACAAATATGTTCAAGGTTT
GGCGACATTGATAAAATTAAGCTTATCTATGACAAAGGTGGAAACTCGAAGCAATACTGCTTCGTATATTACAAA
AATCACATTTCTGCAATTGAGGCAAAACACAATCTGGACAAAAAACCACTTCAAGGACGCGAAATTTCAGTGCGT
TTTGTTCCAGAAAAGGAGTAATAAAGTGATATATGACGCGGCCGCGACTCTAGATCATAATCAGCCATACCACAT
TTGTAGAGGTTTTACTTGCTTTAAAAAACCTCCCACACCTCCCCCTGAACCTGAAACATAAAATGAATGCAATTGT
TGTTGTTAACTTGTTTATTGCAGCTTATAATGGTTACAAATAAAGCAATAGCATCACAAATTTCACAAATAAAGCA
TTTTTTTCACTGCATTCTAGTTGTGGTTTGTCCAAACTCATCAATGTatcttaaTTAACCATTGTGGGAACCGTGCGAT
CAAACAAACGCGAGATACCGGAAGTACTGAAAAACAGTCGCTCCAGGCCAGTGGGAACATCGATGTTTTGTTTT
GACGGACCCCTTACTCTCGTCTCATATAAACCGAAGCCAGCTAAGATGGTATACTTATTATCATCTTGTGATGAGG
ATGCTTCTATCAACGAAAGTACCGGTAAACCGCAAATGGTTATGTATTATAATCAAACTAAAGGCGGAGTGGACA
CGCTAGACCAAATGTGTTCTGTGATGACCTGCAGTAGGAAGACGAATAGGTGGCCTATGGCATTATTGTACGGA
ATGATAAACATTGCCTGCATAAATTCTTTTATTATATACAGCCATAATGTCAGTAGCAAGGGAGAAAAGGTTCAA
AGTCGCAAAAAATTTATGAGAAACCTTTACATGAGCCTGACGTCATCGTTTATGCGTAAGCGTTTGGAAGCTCCT
ACTTTGAAGAGATATTTGCGCGATAATATCTCTAATATTTTGCCAAATGAAGTGCCTGGTACATCAGATGACAGT
ACTGAAGAGCCAGTAATGAAAAAACGTACTTACTGTACTTACTGCCCCTCTAAAATAAGGCGAAAGGCAAATGC
ATCGTGCAAAAAATGCAAAAAAGTTATTTGTCGAGAGCATAATATTGATATGTGCCAAAGTTGTTTCTGACTGAC
TAATAAGTATAATTTGTTTCTATTATGTATAAGTTAAGCTAATTACTTATTTTATAATACAACATGACTGTTTTTAAA
GTACAAAATAAGTTTATTTTTGTAAAAGAGAGAATGTTTAAAAGTTTTGTTACTTTATAGAAGAAATTTTGAGTTT
TTGTTTTTTTTTAATAAATAAATAAACATAAATAAATTGTTTGTTGAATTTATTATTAGTATGTAAGTGTAAATATA
ATAAAACTTAATATCTATTCAAATTAATAAATAAACCTCGATATACAGACCGATAAAACACATGCGTCAATTTTAC
GCATGATTATCTTTAACGTACGTCACAATATGATTATCTTTCTAGGGTTAAATAATAGTTTCTAATTTTTTTATTATT
CAGCCTGCTGTCGTGAATACCGTATATCTCAACGCTGTCTGTGAGATTGTCGTATTCTAGCCTTTTTAGTTTTTCGC
TCATCGACTTGATATTGTCCGACACATTTTCGTCGATTTGCGTTTTGATCAAAGACTTGAGCAGAGACACGTTAAT
CAACTGTTCAAATTGATCCATATTAACGATATCAACCCGATGCGTATATGGTGCGTAAAATATATTTTTTAACCCTC
TTATACTTTGCACTCTGCGTTAATACGCGTTCGTGTACAGACGTAATCATGTTTTCTTTTTTGGATAAAACTCCTAC
TGAGTTTGACCTCATATTAGACCCTCACAAGTTGCAAAACGTGGCATTTTTTACCAATGAAGAATTTAAAGTTATT
TTAAAAAATTTCATCACAGATTTAAAGAAGAACCAAAAATTAAATTATTTCAACAGTTTAATCGACCAGTTAATCA
ACGTGTACACAGACGCGTCGGCAAAAAACACGCAGCCCGACGTGTTGGCTAAAATTATTAAATCAACTTGTGTTA
TAGTCACGGATTTGCCGTCCAACGTGTTCCTCAAAAAGTTGAAGACCAACAAGTTTACGGACACTATTAATTATTT
GATTTTGCCCCACTTCATTTTGTGGGATCACAATTTTGTTATATTTTTAAACAAAGCTTGGCACTGGCCGTCGTTTT
ACAACGTCGTGACTGGGAAAACCCTGGCGTTACCCAACTTAATCGCCTTGCAGCACATCCCCCTTTCGCCAGCTG
GCGTAATAGCGAAGAGGCCCGCACCGATCGCCCTTCCCAACAGTTGCGCAGCCTGAATGGCGAATGGCGCCTGA
TGCGGTATTTTCTCCTTACGCATCTGTGCGGTATTTCACACCGCATATGGTGCACTCTCAGTACAATCTGCTCTGAT
GCCGCATAGTTAAGCCAGCCCCGACACCCGCCAACACCCGCTGACGCGCCCTGACGGGCTTGTCTGCTCCCGGCA
TCCGCTTACAGACAAGCTGTGACCGTCTCCGGGAGCTGCATGTGTCAGAGGTTTTCACCGTCATCACCGAAACGC
GCGAGACGAAAGGGCCTCGTGATACGCCTATTTTTATAGGTTAATGTCATGATAATAATGGTTTCTTAGACGTCA
GGTGGCACTTTTCGGGGAAATGTGCGCGGAACCCCTATTTGTTTATTTTTCTAAATACATTCAAATATGTATCCGC
TCATGAGACAATAACCCTGATAAATGCTTCAATAATATTGAAAAAGGAAGAGTATGAGTATTCAACATTTCCGTG
TCGCCCTTATTCCCTTTTTTGCGGCATTTTGCCTTCCTGTTTTTGCTCACCCAGAAACGCTGGTGAAAGTAAAAGAT
GCTGAAGATCAGTTGGGTGCACGAGTGGGTTACATCGAACTGGATCTCAACAGCGGTAAGATCCTTGAGAGTTT
TCGCCCCGAAGAACGTTTTCCAATGATGAGCACTTTTAAAGTTCTGCTATGTGGCGCGGTATTATCCCGTATTGAC
GCCGGGCAAGAGCAACTCGGTCGCCGCATACACTATTCTCAGAATGACTTGGTTGAGTACTCACCAGTCACAGA
AAAGCATCTTACGGATGGCATGACAGTAAGAGAATTATGCAGTGCTGCCATAACCATGAGTGATAACACTGCGG

CCAACTTACTTCTGACAACGATCGGAGGACCGAAGGAGCTAACCGCTTTTTTGCACAACATGGGGGATCATGTA
ACTCGCCTTGATCGTTGGGAACCGGAGCTGAATGAAGCCATACCAAACGACGAGCGTGACACCACGATGCCTGT
AGCAATGGCAACAACGTTGCGCAAACTATTAACTGGCGAACTACTTACTCTAGCTTCCCGGCAACAATTAATAGA
CTGGATGGAGGCGGATAAAGTTGCAGGACCACTTCTGCGCTCGGCCCTTCCGGCTGGCTGGTTTATTGCTGATA
AATCTGGAGCCGGTGAGCGTGGGTCTCGCGGTATCATTGCAGCACTGGGGCCAGATGGTAAGCCCTCCCGTATC
GTAGTTATCTACACGACGGGGAGTCAGGCAACTATGGATGAACGAAATAGACAGATCGCTGAGATAGGTGCCTC
ACTGATTAAGCATTGGTAACTGTCAGACCAAGTTTACTCATATATACTTTAGATTGATTTAAAACTTCATTTTTAAT
TTAAAAGGATCTAGGTGAAGATCCTTTTTGATAATCTCATGACCAAAATCCCTTAACGTGAGTTTTCGTTCCACTG
AGCGTCAGACCCCGTAGAAAAGATCAAAGGATCTTCTTGAGATCCTTTTTTTCTGCGCGTAATCTGCTGCTTGCAA
ACAAAAAAACCACCGCTACCAGCGGTGGTTTGTTTGCCGGATCAAGAGCTACCAACTCTTTTTCCGAAGGTAACT
GGCTTCAGCAGAGCGCAGATACCAAATACTGTTCTTCTAGTGTAGCCGTAGTTAGGCCACCACTTCAAGAACTCT
GTAGCACCGCCTACATACCTCGCTCTGCTAATCCTGTTACCAGTGGCTGCTGCCAGTGGCGATAAGTCGTGTCTTA
CCGGGTTGGACTCAAGACGATAGTTACCGGATAAGGCGCAGCGGTCGGGCTGAACGGGGGGTTCGTGCACACA
GCCCAGCTTGGAGCGAACGACCTACACCGAACTGAGATACCTACAGCGTGAGCTATGAGAAAGCGCCACGCTTC
CCGAAGGGAGAAAGGCGGACAGGTATCCGGTAAGCGGCAGGGTCGGAACAGGAGAGCGCACGAGGGAGCTT
CCAGGGGGAAACGCCTGGTATCTTTATAGTCCTGTCGGGTTTCGCCACCTCTGACTTGAGCGTCGATTTTTGTGAT
GCTCGTCAGGGGGGCGGAGCCTATGGAAAAACGCCAGCAACGCGGCCTTTTTACGGTTCCTGGCCTTTTGCTGG
CCTTTTGCTCACATGTTCTTTCCTGCGTTATCCCCTGATTCTGTGGATAACCGTATTACCGCCTTTGAGTGAGCTGA
TACCGCTCGCCGCAGCCGAACGACCGAGCGCAGCGAGTCAGTGAGCGAGGAAGCGGAAGAGCGCCCAATACGC
AAACCGCCTCTCCCCGCGCGTTGGCCGATTCATTAATGCAGCTGGCACGACAGG
